# Heavy Chain CDR 3 and Junctional Length Biases in Human Antibody Repertoires Associated with Heavy and Light Chain Germline Utilization

**DOI:** 10.1101/664714

**Authors:** Kannan Sankar, Kam Hon Hoi, Isidro Hötzel

**Author notes:** Corresponding author: Isidro Hötzel, Ph.D., Department of Antibody Engineering, Genentech, South San Francisco, CA 94080, USA.

## Abstract

Antibody variable domain sequence diversity is generated by recombination of germline segments. The third complementarity-determining region of the heavy chain (CDR H3) is the region of highest sequence diversity and is formed by the joining of heavy chain V_H_, D_H_ and J_H_ germline segments combined with random nucleotide trimming and additions between these segments. We show that CDR H3 length distribution is biased in human antibody repertoires as a function of V_H_, V_L_ and J_H_ germline segment utilization. Most length biases are apparent in the naïve B cell compartment, with a significant bias towards shorter CDR H3 sequences observed in association with a subset of V_H_ and V_L_ germlines in the antigen experienced compartment. Similar biases were not observed in nonproductive heavy chain recombination products, indicating selection of the repertoire during B cell maturation as a major driver of the length biases. Some V_H_-associated CDR H3 length biases are dependent on utilization of specific J_H_ germline segments in a manner not directly linked to J_H_ segment length in the germline, but are rather associated with selection of differentially trimmed J_H_ segments in the naïve compartment. In addition, D_H_ segment and N-region random nucleotide insertion lengths within CDR H3 in the naïve compartment were also biased by specific V_H_/J_H_ germline combinations, indicating a complex set of constraints between germline segments selected during repertoire maturation. Our findings reveal biases in the antibody diversity landscape shaped by V_H_, V_L_, and J_H_ germline features with implications for mechanisms of naïve and immune repertoire selection.

## Introduction

The diversity of sequences in the variable regions of immunoglobulins is the basis for the ability of these molecules to bind a virtually unlimited number of antigenic structures. Sequence diversity in the primary repertoire is created by recombination of germline segments for both the heavy and light chains which results in the formation of full-length immunoglobulin variable region exons (1). The light chain variable region is created by the joining of V_L_ and J_L_ germlines while the V_H_ region is created by recombination of V_H_, D_H_ (or D) and J_H_ germlines. The process of recombination starts with the heavy chain in progenitor B cells, initiated by D/J_H_ recombination followed by V_H_/DJ_H_ recombination (2, 3). Light chain recombination occurs in pre-B cells after successful completion of the heavy chain recombination. Germline segments in both chains are also trimmed and extended by a variable number of nucleotides by exonucleolytic nibbling of germline segments and random nucleotide incorporation in the N-regions flanking the D germline mediated by terminal deoxynucleotidyl transferase and germline palindromic duplications (3). B cell clones with full-length, in-frame variable regions are further selected to remove or induce receptor editing of self-reactive clones to form the naïve immune repertoire (4, 5).

The third complementarity determining region (CDR) of the heavy chain (CDR H3) is the region of highest overall sequence and length diversity in antibody repertoires (1). CDR H3 length approximates a Gaussian distribution (6). Average CDR H3 length varies as a function of species, age, isotype, B cell development stage and disease state (6-13). CDR H3 amino acid composition is also biased in a CDR H3 length-dependent manner, associated with differential incorporation of D and J_H_ germline sequences of different lengths and sequence composition into CDR H3 of different lengths (6). Beyond the germline-associated biases, CDR H3 has been shown to undergo different biases during B cell maturation. In particular, a bias towards shorter average CDR H3 lengths is observed in mature relative to immature B cells (9). This is accompanied by a reduction of positively charged residue content and hydrophobicity within CDR H3 associated with negative selection of self-reactive clones in the repertoire (9, 11, 14, 15). A similar reduction in CDR H3 length occurs in isotype-switched memory B cells relative to naïve to B cells (10, 16).

The analyses of CDR H3 diversity and biases in health and disease have been mostly performed independently of the V regions contributed by V_H_ and V_L_ germlines (6-11, 17, 18). Except for sequences that are directly incorporated into CDR H3, the impact of V germline segments on CDR H3 properties has not been addressed. In part this is due to the absence of any expectation for systematic CDR H3 biases as a function of V_H_ germline, especially in the naïve B cell compartment prior to selection associated with adaptive immune responses. Analysis of the impact of the V_L_ on CDR H3 has been limited to properties of the third CDR of the light chain, without any evidence of biases (16). Finally, analysis of the impact of J_H_ germlines on CDR H3 biases has been confined to the expected effects of differential J_H_ germline length and sequence composition. A recent analysis of a large dataset of isotype-switched human antibody sequences with paired chain information revealed an unexpected preferential pairing of IGHV3-7 (V_H_3-7) and Vκ2-30 germlines (19) which, in subsequent detailed analysis, was determined to be linked to biases towards shorter CDR H3 lengths associated with both germlines. This prompted us to investigate the extent to which CDR H3 length might be biased as a function of germline use in human immunoglobulin repertoires. Here we describe high-dimensional analyses of CDR H3 sequences from several independently generated human antibody repertoire sequence datasets. Our results uncover biases in CDR H3 and junctional length distributions associated with V_H_, V_L_ and J_H_ germline segment utilization that shape naïve and immune repertoires in unexpected and unpredictable patterns.

## Results

### Sequence datasets

In the present study we analyzed sequences from four previously described B cell repertoire deep sequencing datasets including 3 donors each and a fifth dataset with 8 donors (16, 19-22). We refer to these datasets as TX, WA, CA, MA and SRI. These datasets were sequenced and bioinformatically parsed using a diversity of methods (Table 1), minimizing the impact of methodological biases. For simplicity we refer to the TX CD27^pos^/IgG/IgA, CA, MA IgG/IgA and WA CD27^pos^ subsets as “antigen-experienced” or “AE”, the TX CD27^pos^/IgM as “AE IgM” and the TX and WA CD27^neg^ subsets as “naïve”. The TX and CA datasets include V_H_/V_L_ chain pairing information. The SRI dataset was analyzed separately in this study to avoid over-representation of donors from a single source in pooled data. No antigen-specific selection of B cells was performed for any of the datasets, although the CA and MA datasets include both pre- and post-vaccination samples.

**Table 1.** Datasets used for analysis.

|  | Dataset |  |  |  |  |
| --- | --- | --- | --- | --- | --- |
|  | CA | MA | TX | WA | SRI |
| Reference | 19 | 21,33<sup>a</sup> | 16 | 20 | 22 |
| Donors | 3 | 3 | 3 | 3 | 8 |
| CD27 marker | NA<sup>b</sup> | NA | CD27<sup>neg</sup> (3)<br>CD27<sup>pos</sup> (2) | CD27<sup>neg</sup> (3)<br>CD27<sup>pos</sup> (3) | NA |
| Isotypes | IgG | IgG/IgA | IgM (CD27<sup>neg</sup>)<sup>c</sup><br>IgG/IgA/IgM (CD27<sup>pos</sup>) | NA | IgG/IgM |
| Chain | V<sub>H</sub>/V<sub>L</sub><sup>d</sup> | V<sub>H</sub> | V<sub>H</sub>/V<sub>L</sub><sup>d</sup> | V<sub>H</sub> | V<sub>H</sub> |
| V/J region parsing | Absolve | IgBlast | As published | As published | As published |
| V-region coverage | Full-length | Full-length | Partial | Partial | Partial |
| Unproductive sequences | NA | NA | NA | CD27<sup>neg</sup>, frameshifted<sup>e</sup> | NA |

<sup>a</sup> Illumina re-sequencing data (31) of the original MA samples (19) was used. See Table S3 for details.
<sup>b</sup> Not available
<sup>c</sup> Isotype information ignored for CD27<sup>neg</sup> IgM clones in the TX dataset and includes undefined isotype sequences.
<sup>d</sup> Paired V<sub>H</sub>/V<sub>L</sub> variable region sequence data
<sup>e</sup> Stop codon unproductive sequences not used to minimize impact of sequencing errors. Unproductive sequences in the CD27<sup>pos</sup> subset not used to minimize clonal expansion effects. CD27<sup>neg</sup>, frameshifted unproductive sequences were not filtered for clonotype.

We aimed at identifying properties that are shared among donors and not influenced by clonal expansion related to specific immune responses. To minimize the impact of clonal expansion, the datasets were processed to retain a random sequence from each lineage, or clonotype. Clonotype definition varied according to sequencing and parsing method used for each dataset (Table S1). Nonproductive sequences were not grouped by clonotype. Overall distribution of CDR H3 lengths was not significantly affected by removal of redundant sequences in most datasets except for the WA and MA AE compartments, which had subtle but noticeable shifts (Fig. S1A). Germline-specific analyses were performed with germlines having at least 240 counts for all donors in a dataset, which represent 94 to 98% of the repertoire in the CA, TX and MA datasets. Due to ambiguities in V_H_ germline calls in the WA dataset, germline-specific analyses in this dataset were performed with 16 V_H_ germlines that had fewer than 10% ambiguous (“in ties”) calls in the dataset, totaling about one third of the repertoire (Table S2). Analyses of whole repertoire data included all sequences filtered by clonotype in specific B cell subsets, regardless of germline classification. The overall AE CDR H3 length distributions are similar among datasets, allowing pooling data from different datasets for the AE B cell subset (Fig. S1B). However, the relative CDR H3 length distributions of the WA and TX naïve B cell subsets differed by an average of 0.9 residues (Fig. S1B) and were thus analyzed separately.

### Average CDR H3 length varies with V_H_ and V_L_ germline use

As a first step we analyzed average CDR H3 length as a function of V_H_ or V_L_ germline use. Average CDR H3 length in the AE subset varied by up to 3 amino acid residues as a function of V_H_ germline use and correlated well for different datasets, with relatively little deviation from length equivalence for different germlines between datasets (Fig. 1A). Average CDR H3 length also varied as a function of V_L_ germline use by up to 4 amino acid residues in the AE compartment and correlated well between the CA and TX datasets (Fig. 1B). The naïve compartment showed a more limited spread in average CDR H3 lengths relative to the AE compartment (Fig. 1C-F, blue squares). Significant reductions in average CDR H3 length in the AE relative to naïve compartments were associated with a subset of V_H_ and V_L_ germlines (Fig. 1C-F). The TX AE IgM subset showed similar trends as the TX AE IgG/IgA subset except that average CDR H3 length was decreased in association with most V_H_ germlines relative to the naïve compartment (Fig. 1C and E).

**Fig. 1.**
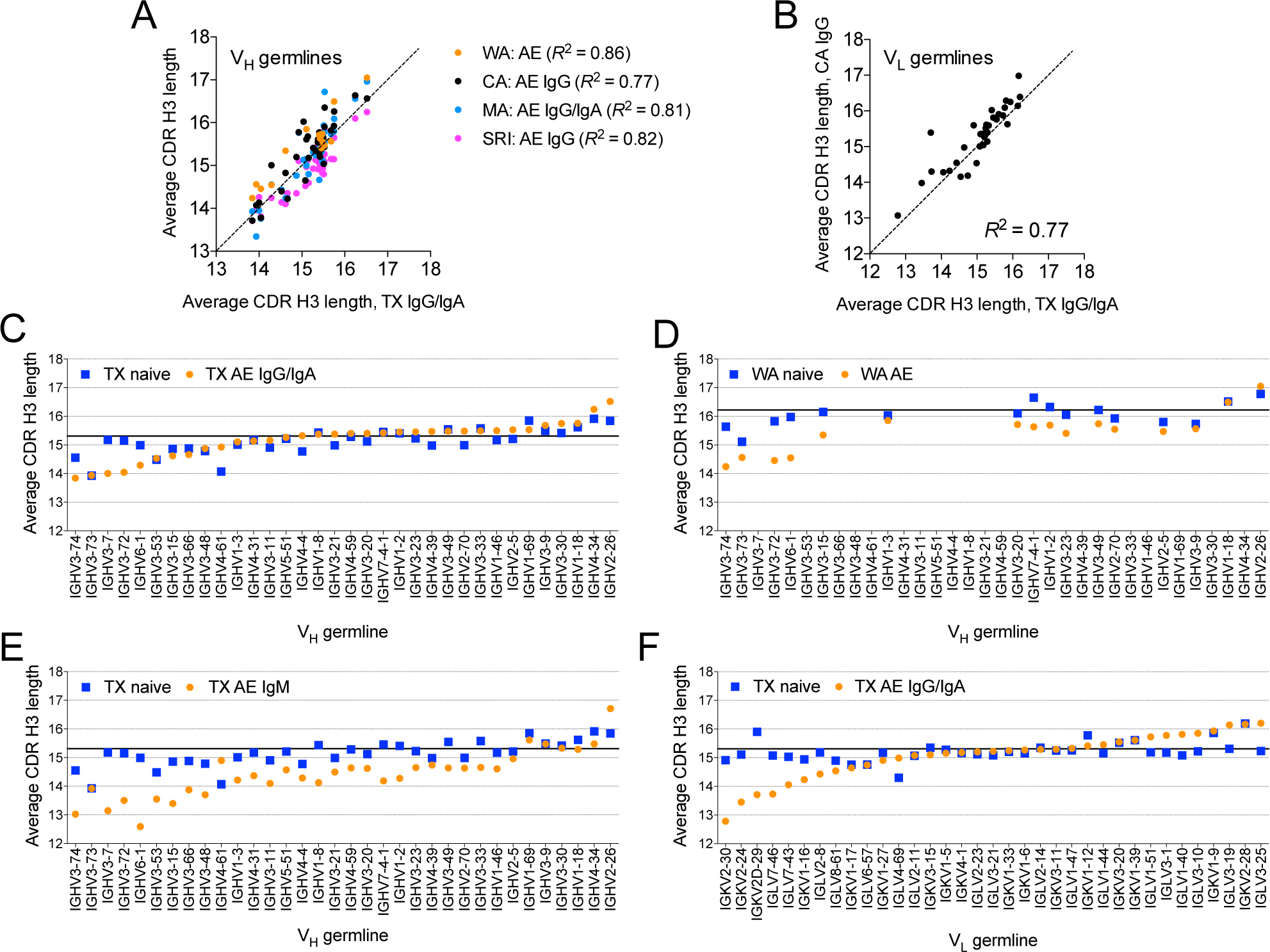
Average CDR H3 length associated with V_H_ and V_L_ segments. Average CDR H3 lengths of TX dataset clonotypes (abscissa) correlated with the CDR H3 of clonotypes with V_H_ (A) and CA V_L_ germlines (B) in the AE compartment. Diagonal lines show 1:1 length correspondence between datasets. (C-F) Average CDR H3 length of TX and WA AE and naïve B cell compartments, segregated by isotype for the TX dataset. Solid horizontal bars indicate average CDR H3 length in the naïve B cell compartment. The order of germlines is the same in panels C, D and E. Lengths are shown in amino acid residues according to the IMGT® CDR definition.

### CDR H3 length distribution varies as a function of V_H_ germline use

We next determined whether CDR H3 length distribution varies with germline use. Overall CDR H3 length distribution of the respective B cell compartment was used as a relative standard to which germline-specific CDR H3 length distributions were compared. This was done to facilitate comparison of biases between samples and also because useful objective reference distributions are not available to determine bias types in naïve compartment sequences. Therefore, most biases described here, including all in the naïve compartment, are relative to the average of the repertoire in each B cell compartment. Overall and germline-specific CDR H3 length distributions were determined by averaging the frequency of each CDR H3 length for all donors across the TX, CA, MA and WA datasets, with the SRI dataset analyzed separately. Statistical analysis of biases was performed in the AE compartment by a two-tailed paired (by donor) *t*-test of length frequencies with a sliding window of two consecutive CDR H3 lengths to minimize the significance of local distribution fluctuations. Observed length distribution biases included overall shifts in average CDR H3 length for sequences with different V_H_ germlines and also obvious and subtle deviations from the overall CDR H3 distribution in discrete ranges of the CDR H3 length spectrum (Fig. 2A, Fig. S2A).

**Fig. 2.**
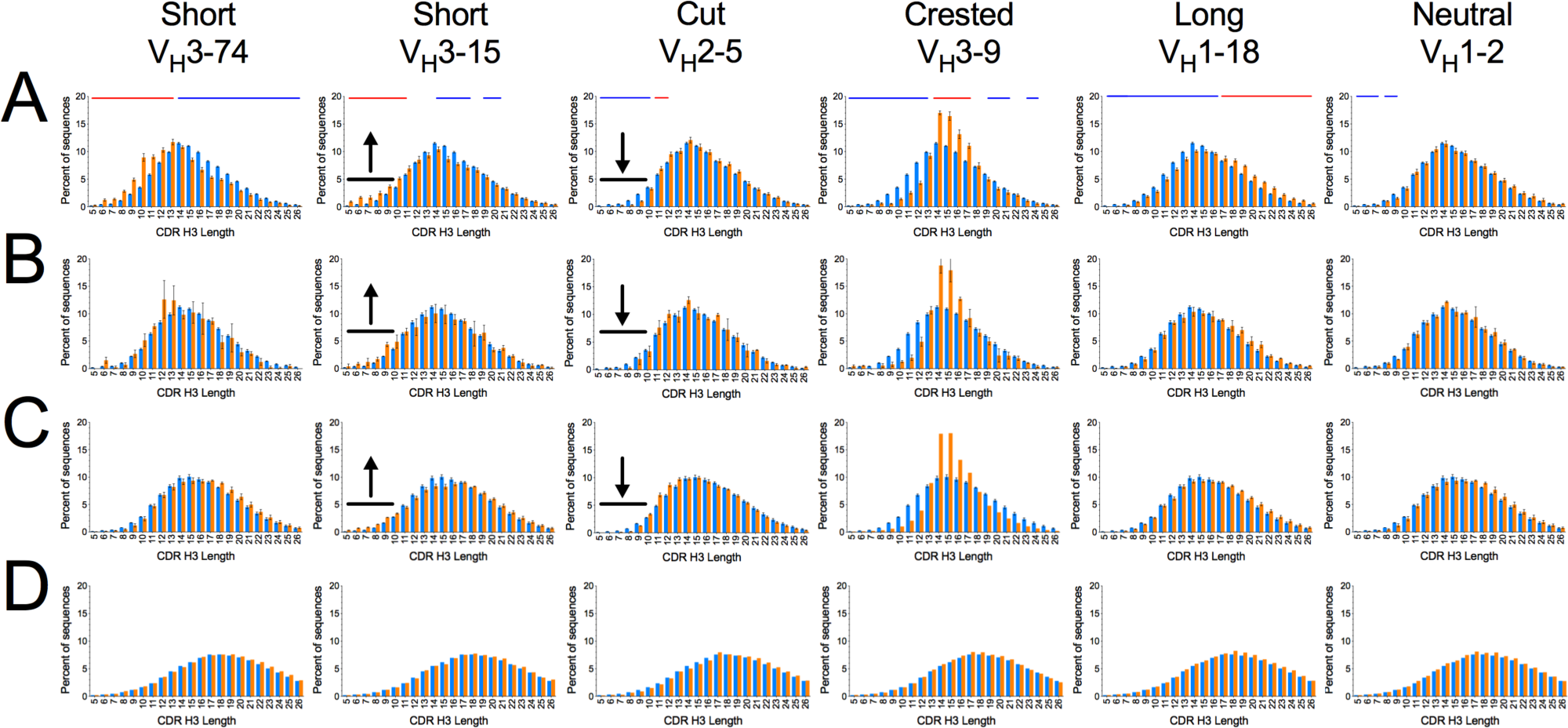
CDR H3 length distribution groups associated with V_H_ germlines. Characteristic examples of each V_H_ bias group are shown, averaged for TX, CA, MA and WA donors in the AE (and/or isotype switched) (A), TX naïve (B), WA naïve (C) and WA nonproductive compartments (D). Orange bars are germline-specific CDR H3 length distributions of unique clonotypes. Blue bars are overall CDR H3 length distributions of unique clonotypes. Blue and red lines above the distributions indicate range of CDR H3 lengths statistically significant different between germline-specific and overall length distributions in a paired (within donor) *t*-test (*P* < 10^-4^) with a sliding window of two contiguous CDR H3 lengths, with red and blue indicating relative enrichment and depletion in the germline-specific distributions. Arrows and black bars highlight subtle depletion of CDR H3 sequence lengths consistent across datasets and B cell compartments. Error bars indicate S.E.M. (A and B) or range (C and D). The full set of distributions is shown in Fig. S2.

To further discern the CDR H3 length biases quantitatively, we performed a principal component (PC) analysis of the length distributions (lengths 5 to 26) associated with different V_H_ germlines to capture the trends causing highest variations in the observed length distributions. Results from the PC analysis were visualized by projecting each germline onto the most important trends, to obtain the so-called PC scores (Fig. 3A). Interpretation of the PC score plots was aided by a visual analysis of the corresponding distributions. Overall, the most significant component, PC1, corresponded to apparent skewness towards shorter or longer lengths, whereas PC2 corresponded to apparent kurtosis, or relative enrichment or depletion of sequences in the mid-range lengths, of the distributions. Using the PC analysis results in conjunction with visual inspection of V_H_ germline-associated CDR H3 distributions compared to the overall CDR H3 distribution in the AE compartment, germlines were categorized by bias type in discrete groups as “Short”, “Neutral”, “Long”, “Cut” and “Crested” (Fig. 2 and 3A, Fig. S2). These groups have different degrees of shifts towards longer or shorter lengths and kurtosis relative to the overall distribution. Differences between the distributions of member of different groups can be subtle, both visually and in the PC analysis. One example is the difference between the Neutral and Cut groups, the latter showing depletion of sequences limited to a narrow band of short CDR H3 lengths. The magnitude of the biases and the details of distribution shapes within each group varied for different V_H_ germlines. However, these were consistent across datasets for each germline, V_H_1-69 being a notable exception (Fig. S3). Germlines in the same V_H_ subfamily did not always belong to the same bias groups. The three major V_H_ subfamilies, V_H_1, V_H_3 and V_H_4, are represented in the Neutral group. The range of germline prevalence in the repertoire was similar for different groups except for the higher prevalence of some germlines in the Crested group (Fig. S4).

**Fig. 3.**
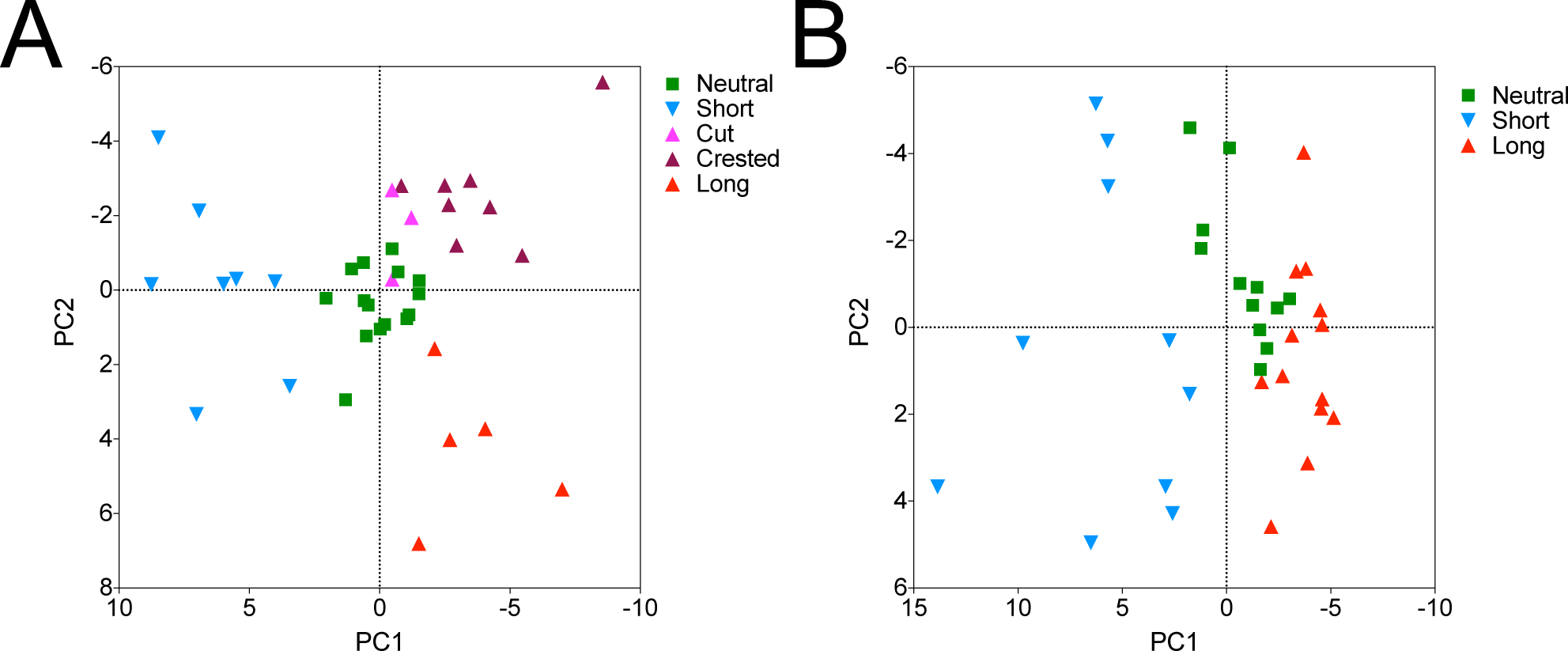
Differentiation of CDR H3 length distributions by PCA analysis. PCA analyses of V_H_ (A) and V_L_ (B) germline-associated CDR H3 length distributions in the AE compartments are shown. Analysis in (A) excludes the WA dataset due to limited germline coverage. Analysis in (B) includes both CA and TX datasets. Each data point indicates a germline for which minimum count requirements were met for the analysis. Points are color-coded by bias group determined by visual inspection of CDR H3 length distributions (Fig. S2 and S6). Axes are oriented to position distributions skewed towards long lengths and with high apparent kurtosis to the right and top respectively.

We determined whether the observed distribution biases were also present in the naïve B cell subset. The Long, Crested and Cut biases were also observed in the naïve B cell compartment, without apparent differences relative to the distributions in the AE compartments (Fig. 2B and C and Fig. S2B, C, E and F). All the germlines in the Neutral group showed average CDR H3 length distribution in the naïve subset as well. However, distribution biases of the Short group in the naïve compartment were variable (Fig. S2B and C), consistent with the average CDR H3 length analysis (Fig. 1E and F). Short biases in the naïve compartment were mostly limited to the V_H_3-73 and V_H_3-15 germlines in the TX and WA datasets. Despite the differences in overall CDR H3 length between the TX and WA naïve datasets, the biases in the naïve compartment had the same trends in both datasets (Fig. 2B and C and Fig. S2B and C).

The data analysis was performed in datasets aggressively filtered for sequences likely to belong to the same lineage. To confirm that biases are not due to pockets of clonal expansion, we performed a repertoire similarity index (RSI) analysis with the CA, TX and MA datasets similar to a recently described method (23). The RSI analysis computes CDR H3 identities within each donor for sequences with the same germline, CDR length and V_H_/V_L_ pairing (for the paired CA and TX datasets) or V_H_ and J_H_ combination (for the unpaired MA dataset). Clonal expansion would thus be reflected by higher than average RSI values. Overall, no significant increase in RSI scores was associated with regions of positive prevalence biases in different parts of the CDR H3 length spectrum for different bias groups (Fig. S5A), confirming that clonal expansion does not account for the observed CDR H3 length biases.

### CDR H3 length distribution bias as a function of V_H_ germline is not generated by VDJ recombination

We next determined whether the biases observed in the naïve compartment are a direct consequence of biases in the VDJ recombination process for each germline. For this we analyzed frameshifted, nonproductive V_H_ sequences that were part of the naïve WA dataset. Nonproductive recombination products are not directly subject to selection and therefore provide information about recombination products prior to any selection of the repertoire. As previously reported, the CDR H3 length of nonproductive V_H_ genes is significantly longer than the productively recombined genes in mature B cell subsets (15). However, CDR H3 length for the nonproductive sequences associated with different V_H_ germlines approximated a Gaussian distribution, with no observable biases associated with different V_H_ germlines relative to the overall repertoire except for minor anomalies associated with some germlines (Fig. 2D and Fig. S2D). Therefore, heavy chain recombination mechanisms do not account for the CDR length distribution biases observed in the naïve repertoire.

### CDR H3 length distribution varies as a function of V_L_ germline use

We performed a similar analysis of CDR H3 length distribution as a function of V_L_ germline and B cell compartment using PC and visual analysis. Similar to the V_H_ germline-associated biases, V_L_-associated biases in the AE compartment could be classified into three groups, named here “Short” (high value of PC1), “Long” (low value of PC1) and “Neutral” (intermediate values of PC1), present in both the CA and TX datasets, each group including a diverse set of V_κ_ and V_λ_ germlines (Fig. 3B and 4A, Fig. S6A). PC1 and PC2 for the light chain were also associated with apparent skewness and kurtosis. The V_L_ Long bias group has Gaussian CDR H3 length distributions, whereas the V_L_ Short bias group includes distribution shapes with significant deviations from Gaussian, including localized frequency spikes in discrete sections in the short range. Only V_κ_ germlines in the Long group were associated with similar CDR H3 length biases in the TX naïve compartment (Fig. 4B, Fig. S6B and C). The magnitude of the V_L_-associated biases varied for different germlines within each bias group but were consistent between datasets (Fig. S7). As above, the RSI analysis results indicated that clonal expansion does not account for the V_L_ germline-associated CDR H3 length biases (Fig. S5B). The prevalence of germlines in the Short group in the repertoire was lower than for germlines of the other two groups (Fig. S4)

**Fig. 4.**
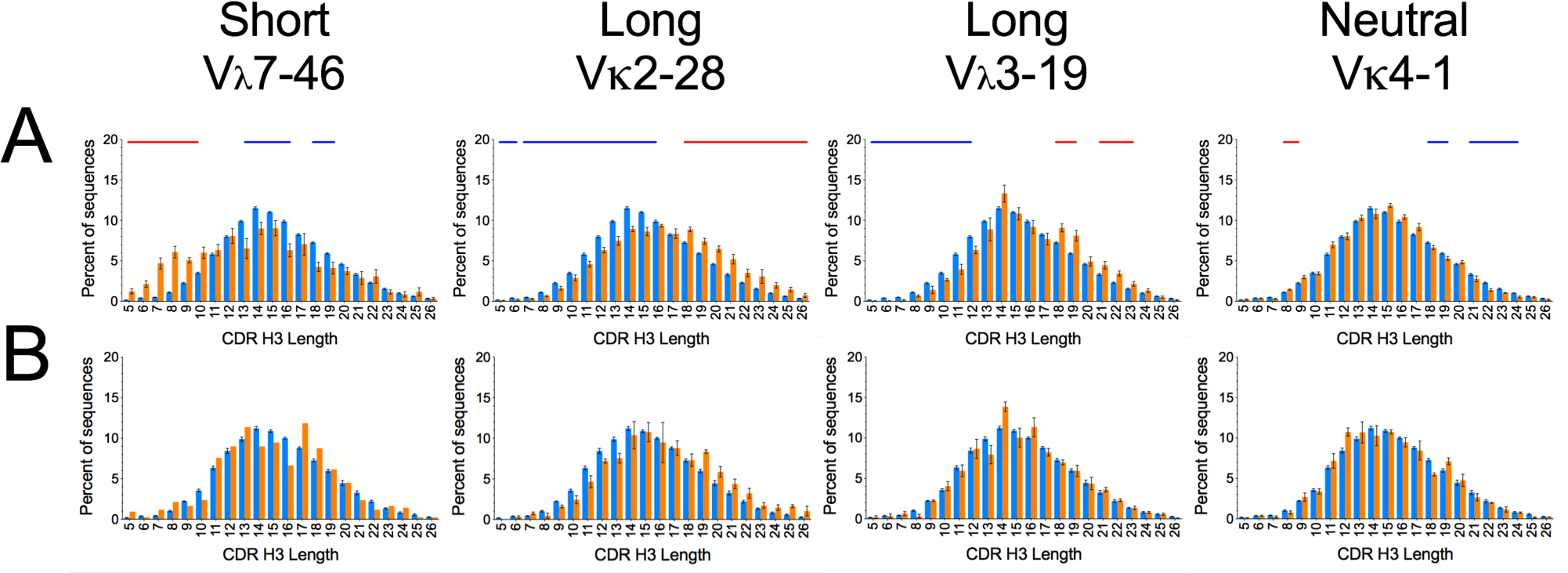
CDR H3 length distribution groups associated with V_L_ germlines. Characteristic examples of each group are shown, averaged for all CA and TX donors in the AE and/or isotype switched compartments (A) and the naïve compartment of TX donors (B). Bar and line colors as in Fig. 1. Error bars indicate S.E.M. for data available from more than 2 donors. The full set of distributions is shown in Fig. S6.

### CDR H3 length is biased as a function of V_H_/J_H_ combination

J_H_ germlines vary in the number of amino acid residues that can be potentially contributed to CDR H3 from 4 in J_H_4 to 9 in J_H_6. We assessed whether differential J_H_ germline usage as a function of V_H_ and V_L_ germline is the basis for V segment-associated CDR H3 length biases. No significant deviations from average J_H_ usage were observed in association with most V_H_ germlines in the WA unproductive sequences (Fig. S8A). Although deviations in J_H_ prevalence linked to some germlines were observed in the naïve compartment of both datasets (e.g., V_H_2-5, V_H_3-9), those deviations do not readily explain CDR H3 distribution biases associated with these V_H_ germlines (Fig. S8B, C and D). One exception was a single V_L_ germline with a bias for longer CDR H3 lengths, Vκ2-28, which was associated with a higher prevalence of the longer J_H_6 and lower prevalence of the shorter J_H_4 germline segments (Fig. S8D). Some of these biases were observed in a reciprocal analysis of J_H_ germline usage as a function of V_H_ germline and CDR H3 length in the naïve compartment but not in nonproductive sequences (Fig. S9). These include not only the generally skewed J_H_ usage proportional to J_H_ germline length, as expected, but also higher or lower than average prevalence of the J_H_5 germline and different J_H_4/J_H_6 germline usage ratios in the average to longer CDR H3 lengths.

We next analyzed CDR H3 length distributions associated with different V_H_/J_H_ germline combinations, comparing these to CDR H3 length distribution of all sequences with the corresponding J_H_ germline. As expected, CDR H3 length distributions were generally shifted according to length of J_H_ the segment in the germline regardless of V_H_ germline (Fig. 5, Fig. S10 and S11). However, a subset of V_H_-associated CDR H3 length biases were impacted by J_H_ germline in a manner independent of length of the J_H_ segment in the germline, with very similar patterns in the naïve WA and IgM/naïve SRI subsets (Fig. 5, Fig. S10 and S11). These included a short CDR H3 length bias associated with sequences with the V_H_3-72, V_H_3-73 and V_H_3-15 germlines and a long bias associated with V_H_1-2, V_H_1-3, V_H_2-5 and V_H_7-4-1 combined with the J_H_5 germline (Fig. 5, Fig. S10). Other CDR H3 length biases associated with specific V_H_/J_H_ germline combinations were also observed, including V_H_3-73/J_H_1, V_H_3-73/J_H_4, V_H_7-4-1/J_H_6 and other minor biases. Two other members of the Crested group, V_H_3-20 and V_H_3-23, showed their characteristic bias when combined with all J_H_ germlines except J_H_2 and J_H_6 (Fig. S10), the longest J_H_ germlines. The Cut group was apparent only when overall population distribution is shifted toward shorter CDR H3 lengths in association with the short J_H_4 germline, making the reductions in the number of sequences in very short range in this group more apparent (Fig. 5). In contrast, the two V_H_ germlines in the Long group that were analyzed, V_H_1-18 and V_H_2-26, were associated with long CDR H3 length biases in the context of most or all J_H_ segments. Distributions associated with V_H_3-9 were unique in that the same peak of sequences with length 14 to 17 occurred independently of J_H_ segment in both datasets. Our results indicate that CDR H3 length distribution biases are not necessarily uniform for each V_H_ germline but may vary as a function of J_H_ germline. In addition, the effect of J_H_ on CDR H3 length distribution is not necessarily similar within V_H_ bias groups, indicating some degree of heterogeneity within bias groups.

**Fig. 5.**
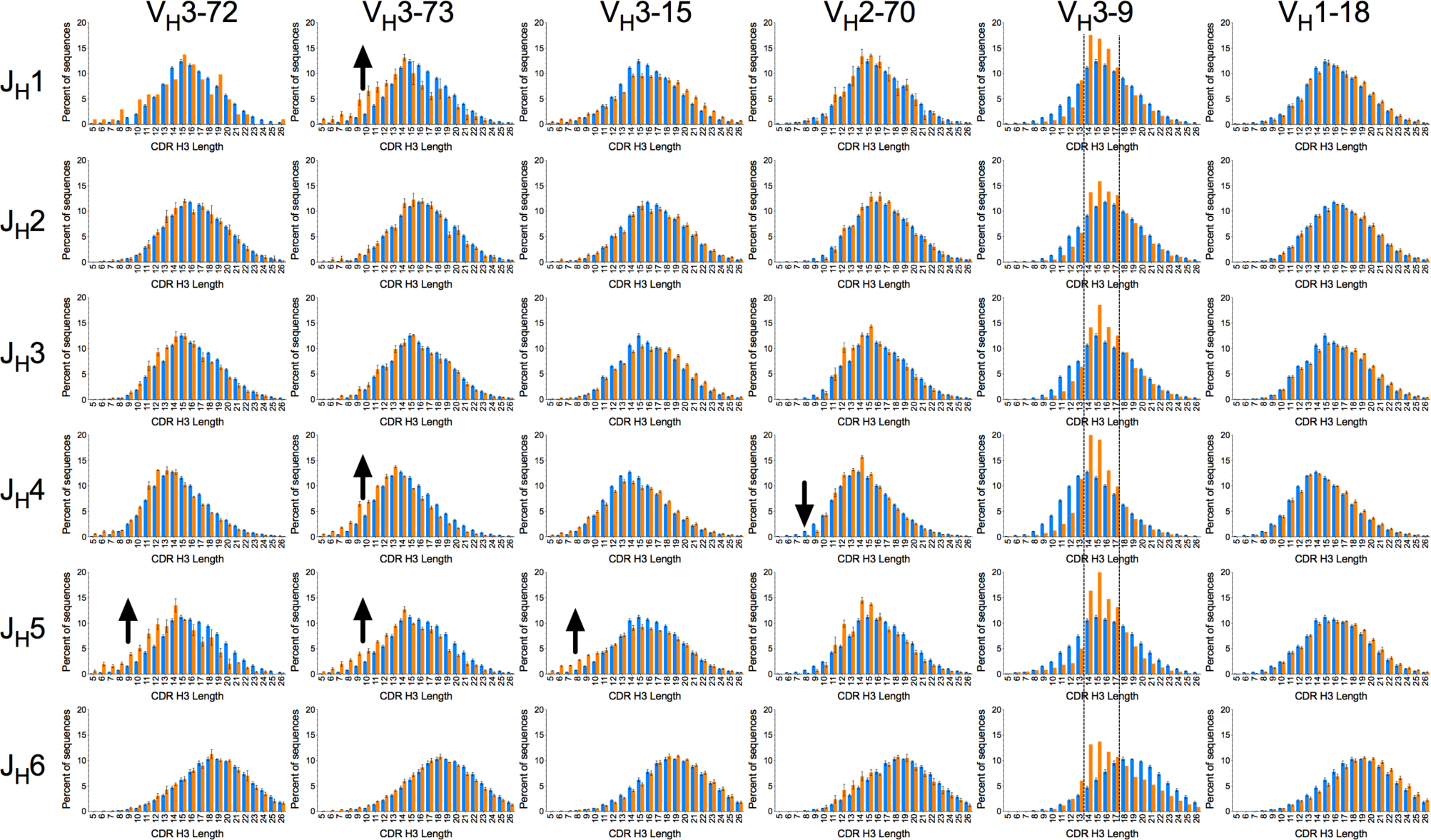
Modulation of V_H_-associated CDR H3 length biases by J_H_ segments in the WA naïve compartment. CDR H3 length distributions associated with individual V_H_/J_H_ combinations (orange bars) are compared to overall repertoire distributions associated with J_H_ segments (blue bars). Arrows highlight biases dependent on J_H_ segment use. Dotted lines indicate the of peaks in the distributions associated with V_H_3-9 and all J_H_ germlines in the same part of the length spectrum. The number of CDR H3 amino acid residues potentially encoded by J_H_1-J_H_6 segments are 6, 7, 6, 4, 6 and 9 respectively. Error bars indicate S.E.M. The full set of distributions is shown in Fig. S10.

### Selection of differentially trimmed J_H_ segments associated with different V_H_ germlines in the naïve compartment

The CDR H3 length distribution biases associated with a subset of V_H_/J_H_ germline combinations may be a consequence of biases in J_H_ trimming as a function of V_H_ germline. J_H_ residue occupancy in the last CDR H3 positions of J_H_4 and J_H_5 sequences was used to indirectly determine J_H_ trimming. The J_H_1, 2, 3 and 6 germlines were not analyzed due to lack of sufficient data or, in the case of J_H_6, absence of apparent J_H_ segment-associated CDR H3 length biases. No apparent biases in J_H_ residue occupancy relative to overall repertoire were observed for any of the analyzed V_H_/J_H_ combinations in the nonproductive WA sequences (Fig. S12). However, J_H_ residue trimming biases were observed for different V_H_/J_H_ combinations in the naïve WA compartment (Fig. 6 and Fig. S12). General trends in residue occupancy in J_H_4 were similar in IgM/naïve SRI sequences for the V_H_/J_H_4 germline combinations with sufficient numbers for analysis (Fig. S13). Residue-specific trimming biases were found to be mostly coordinated for different J_H_ residues in each analyzed V_H_/J_H_ combination, as expected due to the directional nature of trimming. However, closely related V_H_ germlines can be associated with distinct trimming biases of different J_H_4 residues. For instance, V_H_2-5/J_H_4 sequences are associated mostly with reduced trimming of IMGT® residues 114 and 115 (Tyr and Phe) whereas in the case of V_H_2-70/J_H_4 strongly reduced trimming of residue 116 (Asp) was also observed. The results indicate a complex set of constraints leading to selection of differentially trimmed J_H_ segments in the context of certain V_H_ and J_H_ germlines during naïve repertoire maturation.

**Fig. 6.**
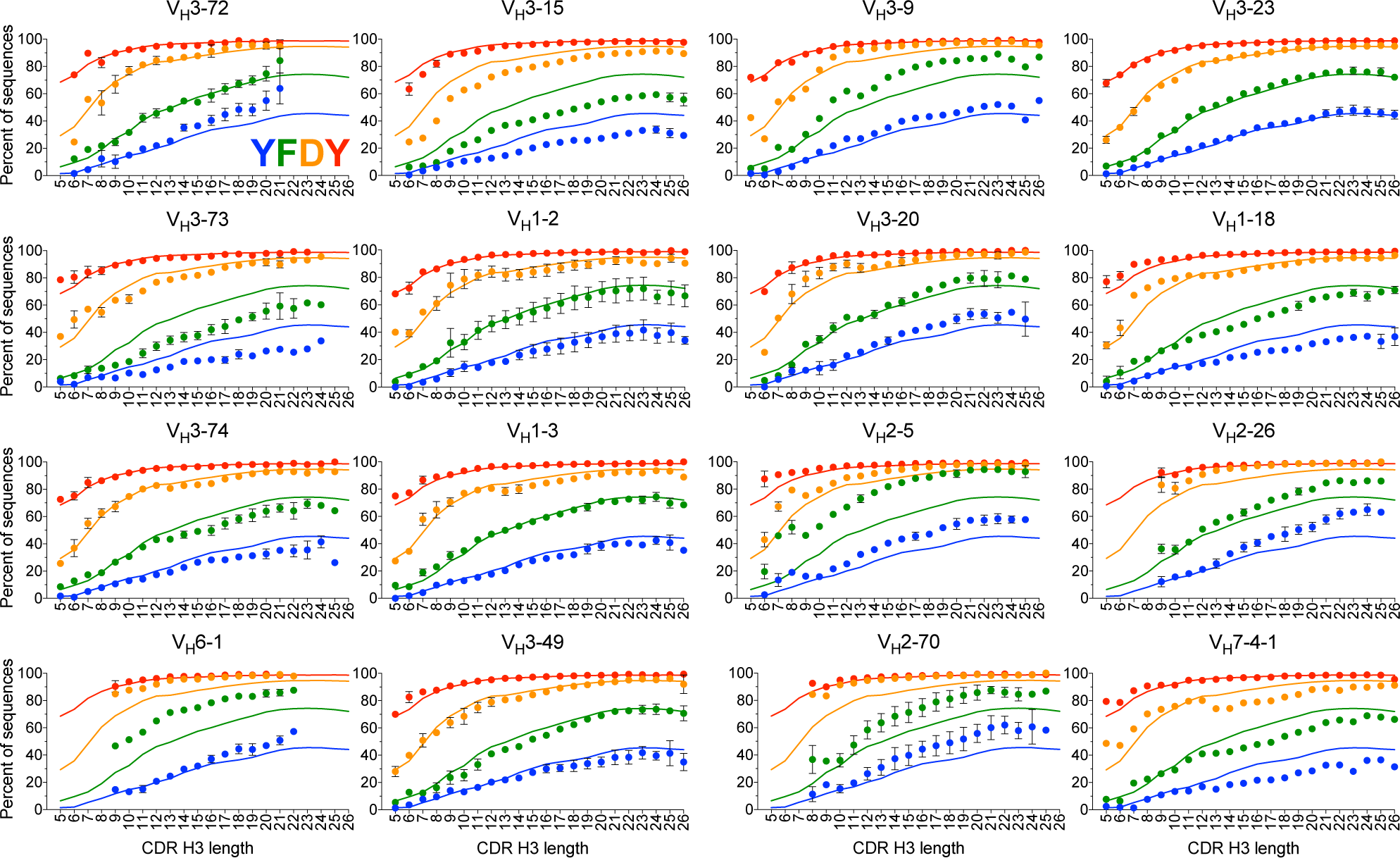
Biases in CDR H3 J_H_4 residue occupancy associated with different V_H_ germlines and CDR H3 length in WA naïve sequences. Occupancy of J_H_ germline-encoded residues in the last CDR H3 positions (IMGT 114 to 117) is shown. Only sequences with J_H_4 are included in the analysis. J_H_ residues are color-coded by position as indicated in the top left panel. Solid lines indicate residue occupancy for all sequences in the naïve repertoire with the J_H_4 germline. Dots indicate average residue occupancy with each V_H_ germline and CDR H3 length. Bars indicate S.E.M. for three donors except for V_H_3-9 and V_H_7-4-1, which are present in one donor each. Data points with fewer than 60 sequences were excluded.

### Biases in D segment and N-region lengths within CDR H3 sequences as a function of V_H_ and J_H_ germline use

The observed J_H_ segment length biases in CDR H3 sequences of specific lengths could be an indirect consequence of biases elsewhere in CDR H3, including the length of V_H_ and D sequences and number of N-region and palindromic nucleotide insertions (NP-region) flanking the D region in CDR H3. Naïve sequences from the 3 donors of the WA dataset and IgM/naïve sequences from 3 of the donors with higher number of sequences in the SRI dataset were parsed for V_H_, J_H_, D and NP-region lengths within CDR H3. In general, no obvious differences on the prevalence of D germlines with different average lengths were observed in association with different V_H_ and the J_H_4 and J_H_5 germlines that would account for J_H_ segment length biases (Fig. S14). One possible exception is V_H_6-1 which was associated with shorter D germlines. In addition, the number of nucleotides that V_H_ can contribute to CDR H3 did not correlate with J_H_ length biases (Fig. 7 and S15). However, different classes of biases in the lengths of D segments and NP-regions were observed for different V_H_/J_H_ combinations, even for clones with the same V_H_ germline (Fig. 7 and S15). Biases had similar trends in the WA and SRI datasets, with differences between datasets observed mostly in the magnitude of the biases. No similar biases were observed in the nonproductive WA sequences with exception of differences in average V_H_-derived sequence lengths associated with V_H_ germline length and a generally shorter NP-region length in V_H_2 clones (Fig. S15), indicating that the observed D and NP-region length biases are mostly selected in naïve repertoire maturation.

**Fig. 7.**
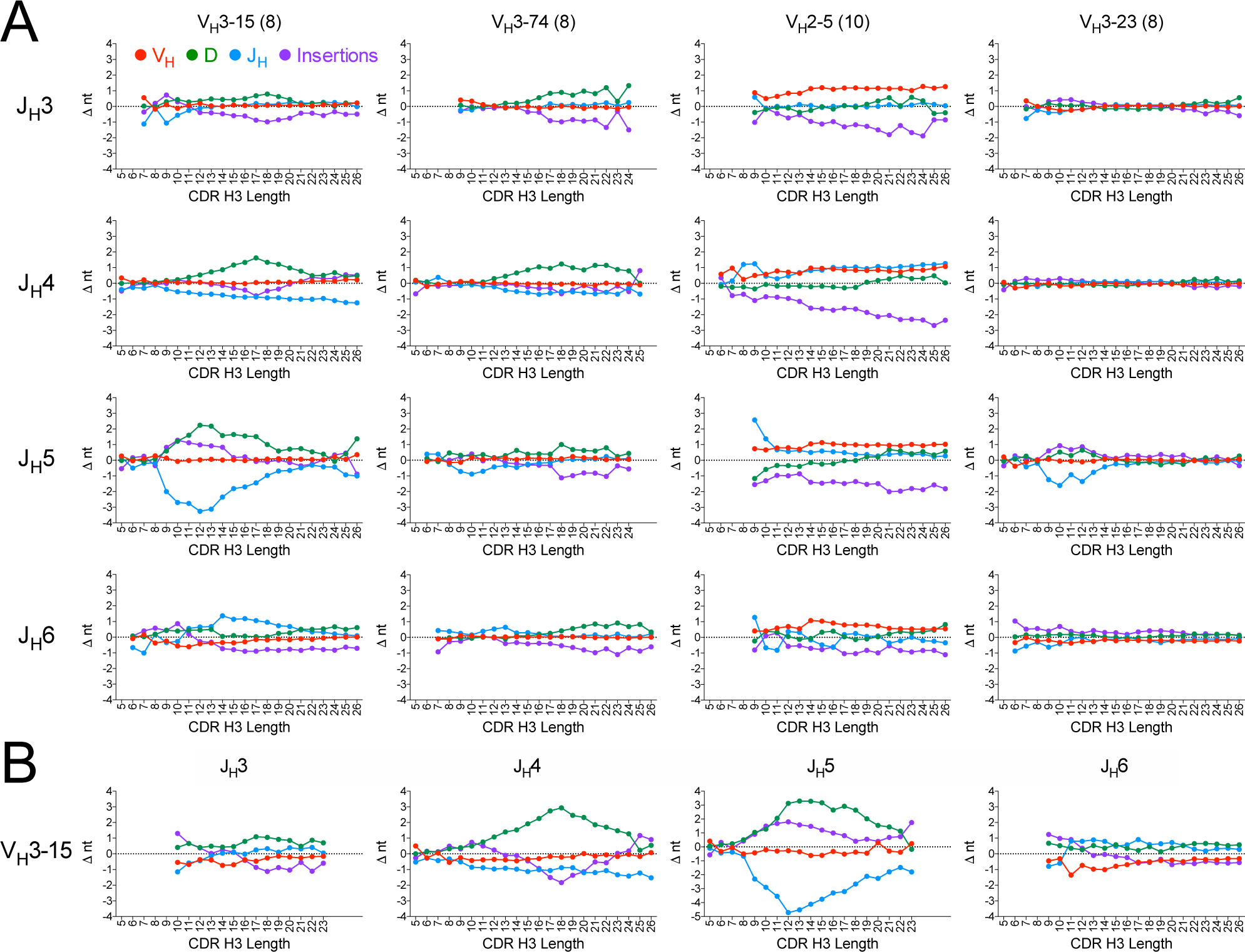
Length biases of regions within CDR H3 as a function of V_H_ and J_H_ germline use and CDR H3 length. (A) Deviations in nucleotides (Δnt) from whole repertoire averages are shown for the V_H_, D, J_H_ segments and NP-region (insertions) in CDR H3 in WA naïve sequences grouped by V_H_ and J_H_ germlines. Colors for each segment are indicated in the top left panel. Values in parentheses indicate the maximum number of V_H_ germline nucleotides that can be included in CDR H3. (B) As in panel (A), showing CDR H3 segment length deviations associated with V_H_3-15 sequences and different J_H_ germlines in the SRI dataset (donors 326650, 326797 and 327059). Sequences with undefined D segments were omitted. Values shown are averages of deviations from overall repertoire, with deviations calculated within donors. Data points with fewer than 60 counts within donors were excluded. Error bars not shown for clarity. CDR H3 lengths are shown in amino acid residues. The full dataset is shown in Fig. S15.

Different classes of junctional biases were observed for different V_H_/J_H_ combinations (Fig. 7A and B and S15). In several V_H_/J_H_ combinations, J_H_ length biases were inversely correlated with both D segment and NP-region length biases including, for instance, V_H_3-23/J_H_5, V_H_3-15/J_H_4, V_H_3-73/J_H_4/J_H_5 and V_H_3-73/J_H_4 combinations. In these cases, the factor determining junctional biases is likely to be the J_H_ segment, with the other junctional components compensating for the J_H_ biases. In V_H_3-74/J_H_4, biases in the D sequence are compensated by both NP-region and J_H_ lengths, suggesting the D segments as the bias determinant. In other cases, such as V_H_3-15/J_H_4, V_H_3-7/J_H_4 and V_H_3-9/J_H_4, D segment and NP-region lengths were inversely correlated with each other without correlations with J_H_ segment length across the CDR H3 length spectrum. In V_H_2 sequences, the reduction in NP-region length observed in nonproductive sequences is maintained in several J_H_ germline combinations in the naïve repertoire and compensates for the longer V_H_ sequences. However, as the J_H_ segments of nonproductive V_H_2 sequences do not appear to be biased relative to the overall repertoire, the observed J_H_4 and J_H_5 trimming biases associated with V_H_2 naïve sequences are presumably due to J_H_ trimming rather than NP-region selection. Overall, the results show different classes of biases in D segment, N-region and J_H_ lengths within CDR H3 of naïve sequences that vary among V_H_/J_H_ germline combinations.

## Discussion

Understanding antibody CDR H3 diversity generation, a process critical for antigen binding, has long been a goal in the immunology and antibody engineering fields. Numerous reports have described overall CDR H3 length and amino acid composition biases in health and disease as well as in different B cell differentiation and developmental stages and species. Here we describe detailed, high-dimensional analyses of CDR H3 and junctional segment length distributions and show a complex set of biases determined by V_H_, V_L_, and J_H_ germline use and B cell maturation state. Most of the length biases we describe are evident in the naïve B cell compartment but not, with very few exceptions, in the nonproductive subset, indicating a major role of naïve B cell repertoire maturation in shaping those biases. In addition, only a subset of V_H_ or V_L_ germlines is associated with biases towards shorter CDR H3 lengths in the antigen-experienced compartment, indicating general germline-specific adaptive immunity selection processes shared among individuals. The V_H_ and V_L_-associated CDR H3 length biases parallel observations of patterns of T cell receptor β chain CDR3 length distribution biases with different TRBV germlines in repertoires that arise in the process of T cell maturation (24), indicating that at least the broader biases are a general phenomenon in immune repertoires. Our results extend these observations by showing biases in the length of junctional segments within CDR H3 determined by V_H_, and J_H_ germline usage in patterns that are not predictable from germline repertoire sequence properties. A recurring theme in the results presented here was that biases observed at one level (e.g., V_H_ germline) were only partly explained by biases at higher dimensional levels (e.g., V_H_/J_H_ combinations), with additional unexpected biases observed in the higher-dimensional levels. It is expected that analyses including other repertoire descriptors not included here will uncover additional biases, one example being germline allelic variation (Fig. S16). Our results provide a framework and baseline for repertoire analyses in disease and immune states beyond the germline usage and average CDR H3 length and composition properties commonly analyzed in repertoire deep sequencing studies.

Special consideration was given to the repeatability and robustness of the findings. The results are based on a total of 12 donors with AE or isotype-switched sequences, 5 of which also had naïve B cell sequences, and confirmed by analysis of 8 additional donors from the SRI dataset. These datasets were obtained and parsed with different sequencing methods and bioinformatic pipelines, which minimizes the impact of technical artifacts. Some of the biases, such as those associated with V_L_, J_H_ and D germlines and NP-regions, cannot be easily generated by sequencing or parsing artifacts. However, subtle differences were observed between datasets which may be due to technical reasons, such as the slightly longer average CDR H3 length of sequences in the WA AE dataset compared to other datasets and the significant differences in average CDR H3 length between the naïve compartments of the TX and WA datasets (Fig. 1). However, these differences in baseline average CDR H3 length, which were accounted for in our naïve compartment analyses, did not affect the overall results. The V germline-associated CDR H3 length biases are not related to clonal expansion as sequences were clustered by clonotype. The stringency of clonotype clustering criteria had limited impact on results. This is exemplified by the WA and SRI datasets, which yielded results comparable with other datasets despite having been clustered by clonotype using a higher CDR H3 sequence identity threshold than other datasets. Limited reliability of D germline classification, especially when sequence identity length is short, is well known. However, any D germline assignment errors would not be expected to be associated with particular V_H_/J_H_ germline combinations sequences. In addition, errors in D parsing should still result in similar length D germline sequence matches, minimizing the impact on D and NP-region length assessments. Thus, the observed D segment length biases associated with sequences with different V_H_/J_H_ germline combinations, generally similar in the WA and SRI dataset subsets analyzed, are not expected to be a consequence of systematic junction parsing errors. This is further supported by the lack of similar biases in nonproductive sequences of similar lengths parsed by the same method, except for the expected germline-specific V_H_ region length biases in CDR H3.

One factor that was not addressed here is haplotype variations within and between donors. Haplotypes could potentially affect CDR H3 length distributions through differences in D germline composition and differential recombination frequencies of D or J_H_ germlines of different lengths in different chromosomes (25). This, combined with differential recombination frequencies of V_H_ alleles (26-28), may impact CDR H3 distributions associated with certain V_H_ germlines. However, heavy chain variable region haplotype differences would not be expected to impact CDR H3 distributions associated with V_L_ germlines and the AE compartment-specific short CDR H3 length biases. In addition, the observation of essentially the same CDR H3 length distribution biases in several donors from five different sources and junctional segment length biases in six donors from two of these sources, along with a lack of systematic associations between V_H_ and D and J_H_ alleles across donors (27, 28), indicates that haplotype variations are unlikely to be a major factor in the CDR H3 and junctional length distribution biases described here.

The analyses shown here use germline information as a proxy for undefined sequence features that ultimately determine the observed biases. Therefore, the selected CDR H3 sequence and structural properties that result in the observed biases and the germline sequence properties that determine those biases remain to be identified. Analysis of V_H_ germline residues that can directly encode or bias CDR H3 IMGT® positions 105 to 107 did not reveal clear correlations between the number or type of encoded residues and most CDR H3 bias groups or junctional segment length biases (Fig. S17). One exception may be the Cut group, which includes two V_H_2 germlines with slightly longer extensions into CDR H3. The extended V_H_ sequences, along with the observed reduced J_H_4 trimming associated with these germlines, may contribute to the low frequency of short CDR H3 sequences in the Cut group. In addition, no obvious correlations between J_H_ trimming biases and variations in V_H_ germline residues in positions 40 to 42 generally contacting the differentially trimmed J_H_ residues 115 and 116 were observed. The differentially trimmed residue 116 is located in a region at the base of CDR H3 that can adopt either a “bulged” or “extended” conformation (29, 30). The factors that determine the more common bulged conformation are not clear and appear to depend on the Ig domain, encoded mostly by V_H_ germlines (30, 31). Whether J_H_ trimming biases reflect biases in the structure of the CDR H3 base as a function of V_H_ germline remains to be determined. The biases in D segment and NP-region lengths appear to be, in several cases, secondary to J_H_ length biases. In other cases, opposite length biases in D segment and NP-region lengths that are independent of J_H_ trimming biases in the context of specific V_H_/J_H_ combinations are observed, indicating selection of differentially trimmed D germline segments. These may be due to selection of different amino acid compositions associated with D and NP-region sequences. Finally, CDR H3 length-specific selection independent of sequence appears to occur in V_H_3-9, in which sequences of lengths 14-17 are enriched regardless of the underlying J_H_ germline length-associated biases. One challenge in determining how different germline regions interact to give rise to the observed biases is the relatively limited number of non-redundant human antibody structures with different germline combinations, extent of J_H_ germline trimming and CDR H3 lengths.

The CDR H3 biases described here pose questions about the functional properties that might shape those biases and the functional consequences of these biases for adaptive immunity. The emergence of some biases in the naïve repertoire suggests selection against self-reactivity, selection for structural integrity, expression or a combination of these factors as possible mechanisms. One factor that seems not to contribute significantly to most or all of these biases is chain pairing, except perhaps for the association between Vκ2-28 and J_H_6 in the naïve compartment. In agreement with previous reports, no significant association between V_H_ and V_L_ germlines of similar bias types was observed, with one exception being the previously described preferential pairing of the V_H_3-7/Vκ2-30 germlines in the Short V_H_ and V_L_ bias groups in the AE compartment (19, 32). If related to selection against self-reactivity, the different biases indicate either that features other than CDR H3 charge and hydrophobicity significantly contribute to self-reactivity or that V segments modulate the self-reactivity mediated by these factors. The bias towards shorter CDR H3 lengths associated with a subset of V_H_ and V_L_ germlines in the AE compartment may be attributable to these same mechanisms or to immune selection. The latter would suggest widespread convergences in human repertoires associated with certain V_H_ and V_L_ germlines or, possibly, some degree of functional specialization in the germline repertoire linked to short CDR H3 sequences, analogous to the association between CDR H3 length and recognition of different antigen classes (33). Our results point to unexpected cross-constraints between V_H_, V_L_, J_H_ and other junctional elements selected at different stages of B cell development that significantly shape antibody repertoires.

## Materials and methods

### Datasets and analysis

Sequences were obtained from the original publications (16, 19, 20) except for the MA dataset. The sequences in the MA dataset were obtained from a re-sequencing by Illumina MiSeq (34) of the same set of samples previously described by Laserson *et al.* (21). A summary of the samples used here is given in Table S3. Sequencing methods for the MA dataset are described in the experiment design section associated with sample data (see https://www.ncbi.nlm.nih.gov/sra/SRX2251687). Sequences were used as parsed in the original publications except for the MA dataset, where the raw sequencing files were processed and germlines annotated with a custom pipeline. Briefly, paired-end reads were merged using FLASH (35) to reconstruct the full-length variable domain sequences using the following parameters: read length at 300 bps, expected fragment length at 530 bps, standard deviation at 50 bps. The full-length sequences were subsequently processed to identify the frameworks and CDR regions using position-weighted motifs as previously described (36). IgBlast (37) was used to supplement the region parsed data with germline annotation for each sequence, including nucleotide somatic mutations. Isotypes of the sequences were determined by finding the closest matching human CH1 isotypes on the available CH1 sequences. Each sequence was processed and annotated with the frameworks, CDRs, germline use and clonotype grouping (see below). Nonproductive sequences in the WA dataset used for analyses were limited to frameshifted sequences in the naïve compartment to minimize the indirect effect of clonal expansion. CDR H3 length of nonproductive, frameshifted sequences in amino acids was set as the nearest integer of CDR H3 length in nucleotides divided by 3. For naïve compartment sequences of Donor 1 of the WA dataset only the D1a repeat was used for most analyses (20). Parsing of D segments was done using Blast (38) after removing the sequences corresponding to V_H_ and J_H_ regions from CDR H3 sequences. An identity of 100% over a span of at least 5 contiguous nucleotides was required for D germline matches. The samples from the D1Nb subset were also included for D segment parsing, removing redundant sequences as described below. All CDR H3 length distributions and germline prevalence analyses were determined using custom scripts and Microsoft Excel 2016. Paired *t*-tests of CDR H3 length distributions were performed using Microsoft Excel 2016. Mann-Whitney tests for distributions were done using GraphPad Prism version 6. The IMGT® CDR definition and numbering system is used throughout (39).

### Clonotype clustering

Clonotypes in the CA and TX datasets were defined as sequences from the same donor, V_H_ and V_L_ germlines and CDR H3 length with a nominal 57% or greater CDR H3 amino acid sequence identity, which better approximates an average 60% CDR H3 sequence identity across the range of CDR H3 lengths. For the TX dataset IgG/IgA, CD27^pos^/IgM and CD27^neg^ sequences were segregated prior to clonotype clustering. The minor fraction of sequences without germline information in the TX dataset was not clustered into clonotypes. Clonotypes in the MA dataset were defined as sequences from the same donor, V_H_ and J_H_ germlines and CDR H3 length with a nominal 57% or greater CDR H3 amino acid sequence identity as above. IgG/IgA and IgM sequences were also segregated prior to clonotype clustering. Clonotypes in the WA dataset were defined as sequences from the same donor with the same V_H_ and J_H_ germline and same CDR H3 length and sequence. If V_H_ germline information was not available then V_H_ subfamily information was used in lieu, retaining as a representative for the clonotype a sequence with V_H_ germline information if available. If J_H_ germline information was not available then this parameter was ignored, also retaining otherwise identical sequences with available J_H_ information as representatives for clonotypes, if available. Nonproductive sequences in the WA dataset were not processed for clonotype clustering. Clonotypes in the SRI dataset were defined as sequences from the same donor with the same V_H_ and J_H_ germline and same CDR H3 length and sequence. Only sequences labeled as “productive” in the SRI dataset were analyzed.

### Repertoire Similarity Index Analysis

RSI was computed in a manner similar to a previously described method (23). For a given set ***s*** of CDR H3 sequences, all of the same length *n*, RSI is measured as follows:

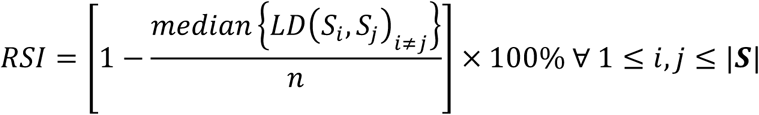

where *S*_*j*_ and *S*_*j*_ refer to any two sequences in the set of CDRH3 sequences and *LD*(*S*_*i*_, *S*_*j*_) refers to the Levenshtein distance function, which measures the number of amino acid changes necessary to convert *S*_*i*_ to *S*_*j*_. For a given V_H_ germline and CDR H3 length, RSI values were computed for those sequences that shared the same V_L_ germline (for the paired CA and TX datasets) or the same J_H_ germline (for the unpaired MA datasets) and the same CDR H3 length. Values were computed separately for each donor in the datasets and averaged for each length. Values shown in graphs in Fig. S5 are the averages in each length for different datasets. All calculations were performed using custom scripts in R.

### Principal component analysis of CDR H3 length distributions

The length distribution of each germline was captured as a vector of length 22 containing the percentage of sequences of length 5 to 26. For V_H_, the values for each germline were averaged over all the AE datasets except the WA dataset due to limited germline coverage. For V_L_, the values were averaged over the CA and TX datasets. The distributions of each germline were consolidated into a matrix ***X***_*n*×22_ where *n* is the number of germlines considered for analysis (*n* = 39 for V_H_ and *n* = 35 for V_L_). In order to capture the differences in behaviors of the germlines the naïve and AE sets, they were considered separately (if at least 240 sequences were available when pooled from all donors in each set). The variance covariance matrix ***S***_**22**×**22**_ of X was computed with elements *S*_*ij*_ as

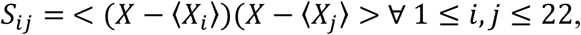

where <> refers to average across all germlines. Eigen decomposition of the matrix ***S*** results in 22 eigenvectors, each of which capture a trend in the distribution as a function of the CDR H3 lengths and are sorted in decreasing order of the variance they capture. Each germline was then projected onto these eigenvectors to obtain the PC scores which enabled visualization of the different trends and comparisons among the different germlines.

## Acknowledgments

We thank Steve Guerrero and Qing Zhang for helpful discussions.

