## Supplementary Figures and Tables for "Heavy Chain CDR 3 and Junctional Length Biases in Human Antibody Repertoires Associated with Heavy and Light Chain Germline Utilization"

Corresponding author:

Isidro Hötzel, Ph.D.

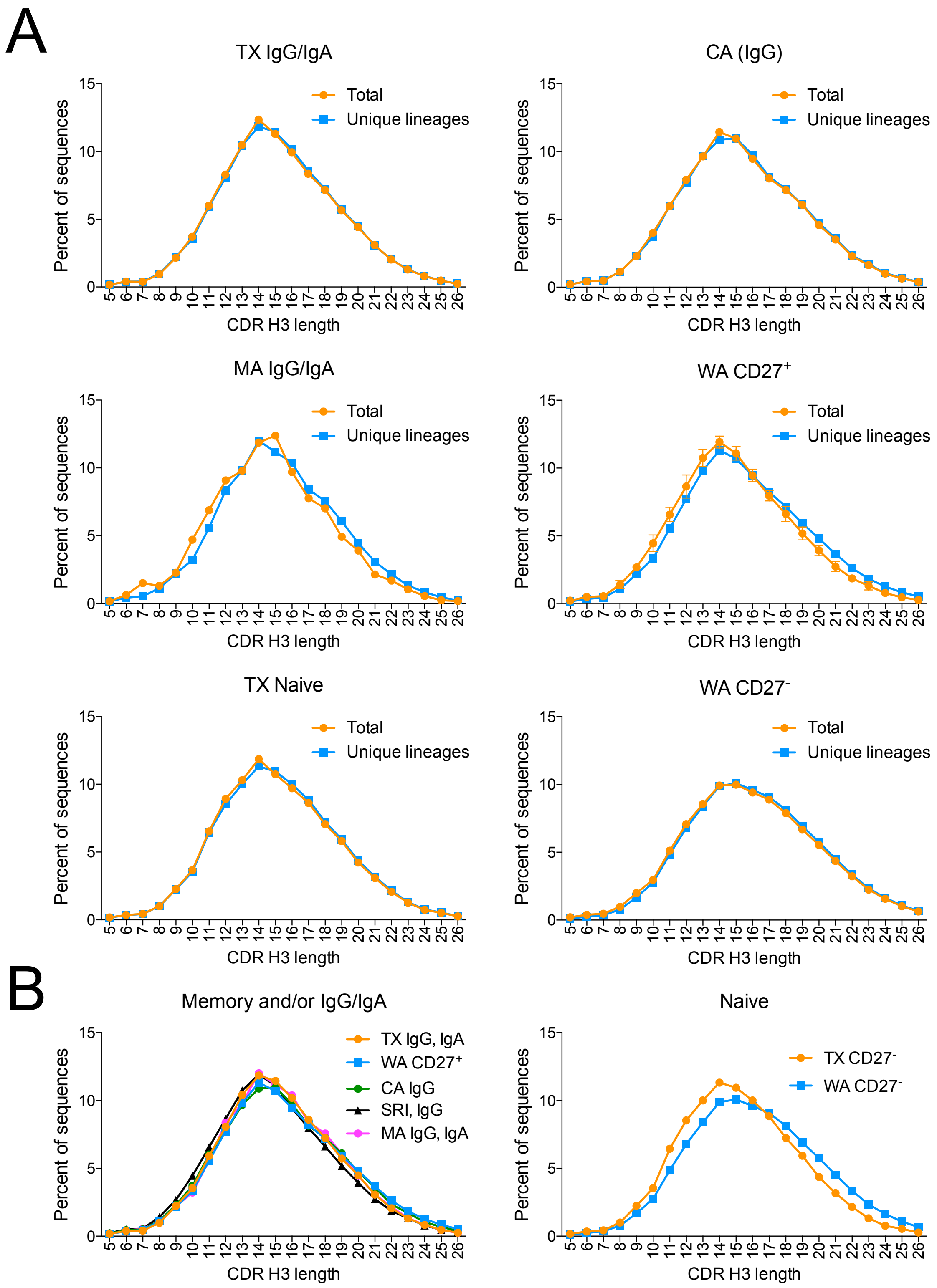


**Fig. S1.** (A) CDR H3 length distribution of raw, as published, data and unique lineage sequences as defined in main text. (B) Comparative CDR H3 length distributions of unique lineages (clonotypes) in the memory and/or IgG/IgA B cell and in the naïve compartments.

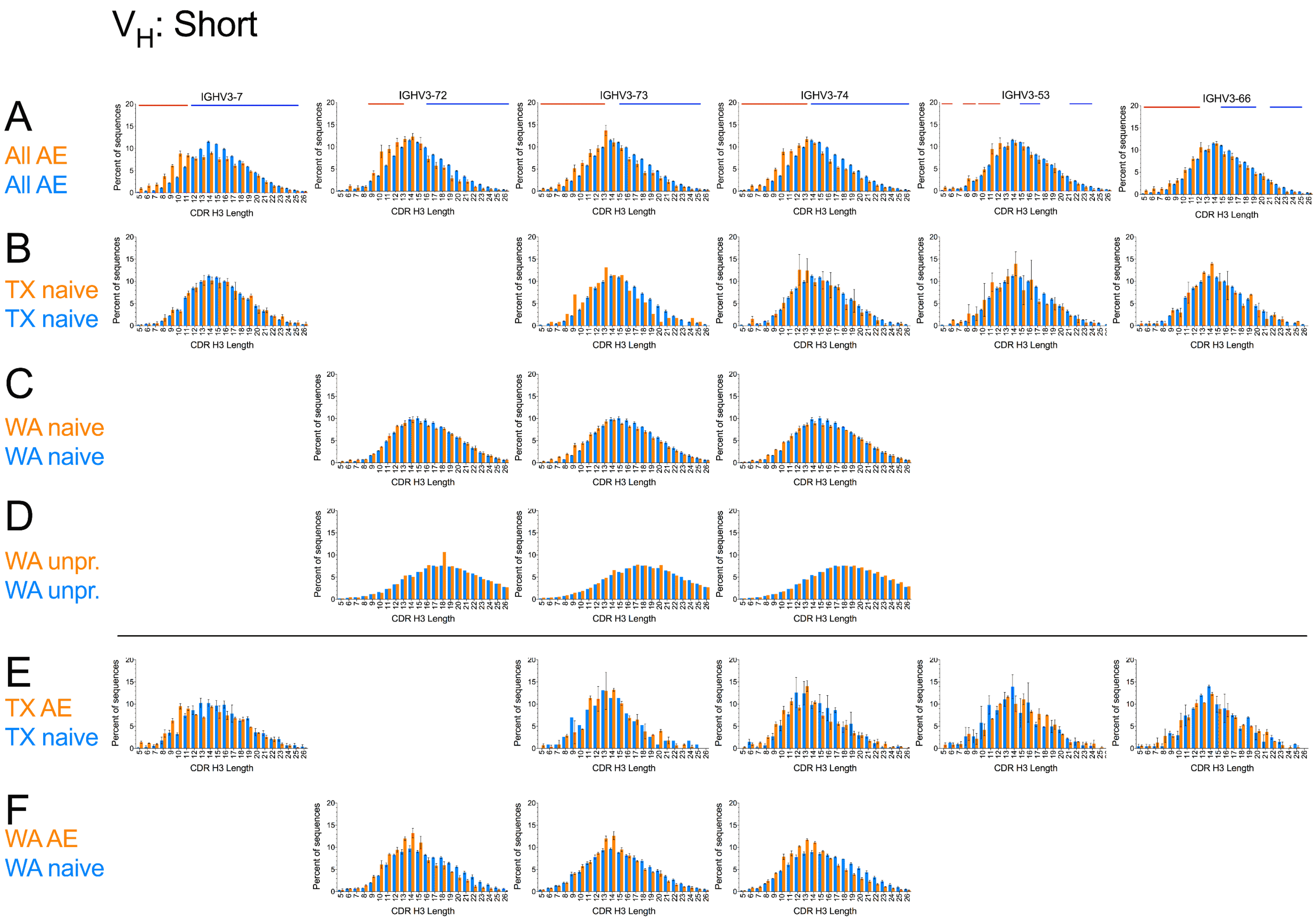
VH: Short
A
All AE
All AE
IGHV3-7
IGHV3-72
IGHV3-73
IGHV3-74
IGHV3-53
IGHV3-66
B
TX naive
TX naive
C
WA naive
WA naive
D
WA unpr.
WA unpr.
E
TX AE
TX naive
F
WA AE
WA naive
Percent of sequences
CDR H3 Length

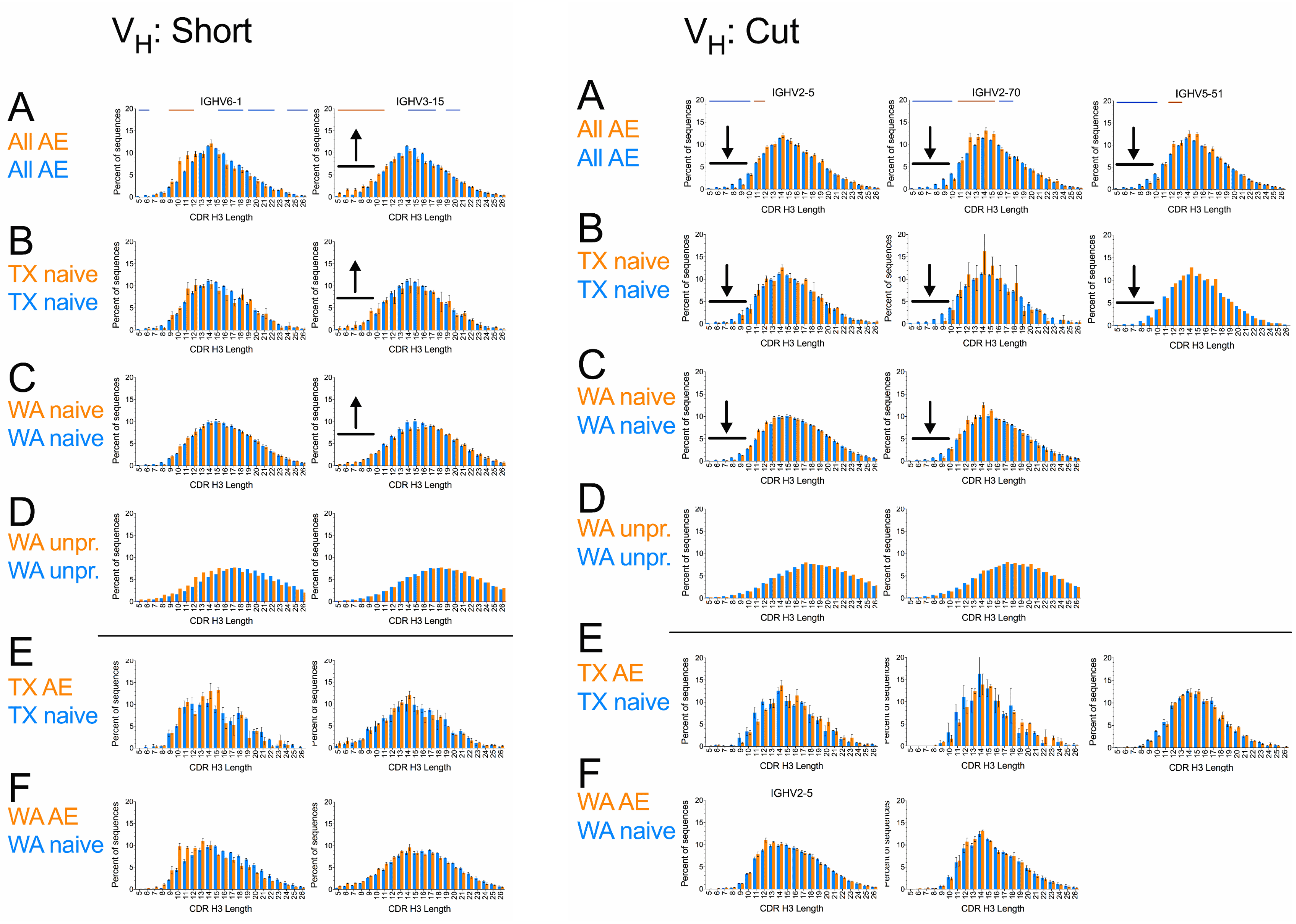

V$_H$: Short
V$_H$: Cut
A
All AE
All AE
B
TX naive
TX naive
C
WA naive
WA naive
D
WA unpr.
WA unpr.
E
TX AE
TX naive
F
WA AE
WA naive
IGHV6-1
IGHV3-15
IGHV2-5
IGHV2-70
IGHV5-51
IGHV2-5
Percent of sequences
CDR H3 Length

### $V_H$: Crested

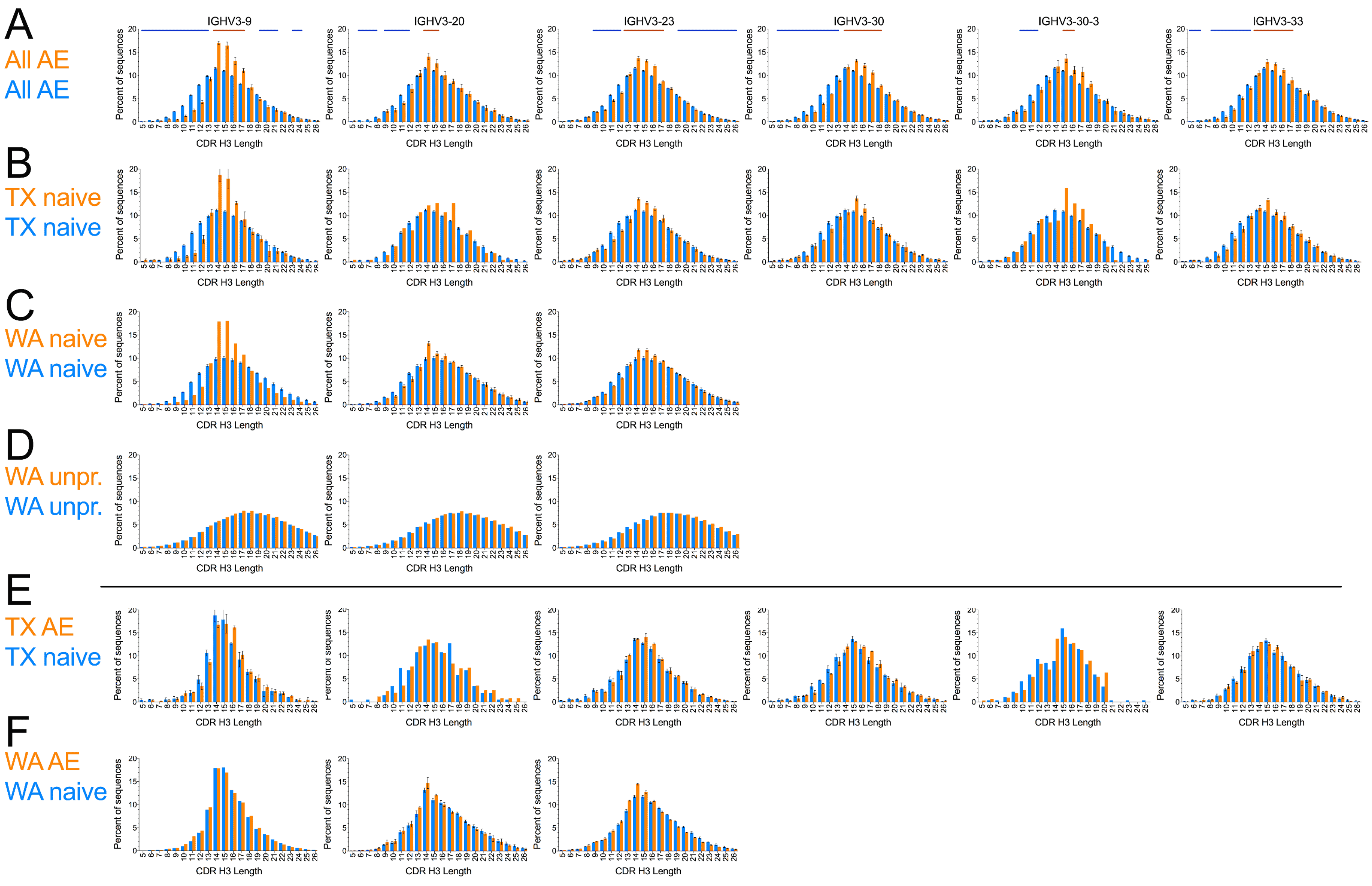

A
All AE
All AE
B
TX naive
TX naive
C
WA naive
WA naive
D
WA unpr.
WA unpr.
E
TX AE
TX naive
F
WA AE
WA naive
IGHV3-9
IGHV3-20
IGHV3-23
IGHV3-30
IGHV3-30-3
IGHV3-33
Percent of sequences
CDR H3 Length

### $V_H$: Crested

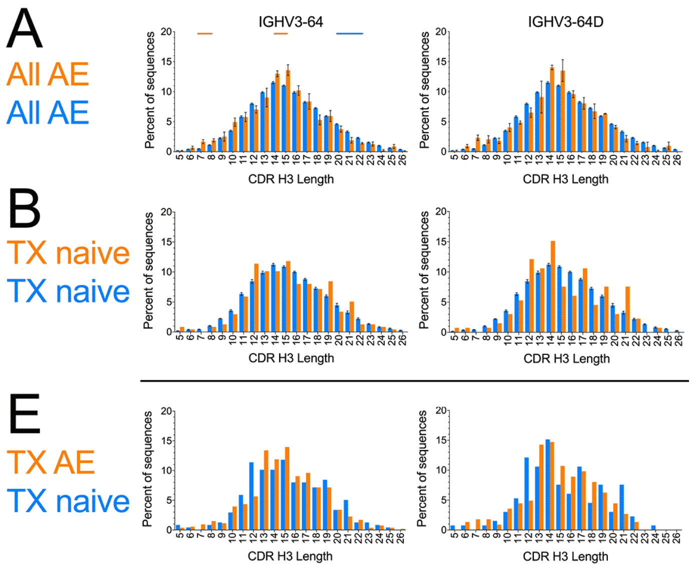
A
All AE
All AE
B
TX naive
TX naive
E
TX AE
TX naive
IGHV3-64
IGHV3-64D
Percent of sequences
CDR H3 Length

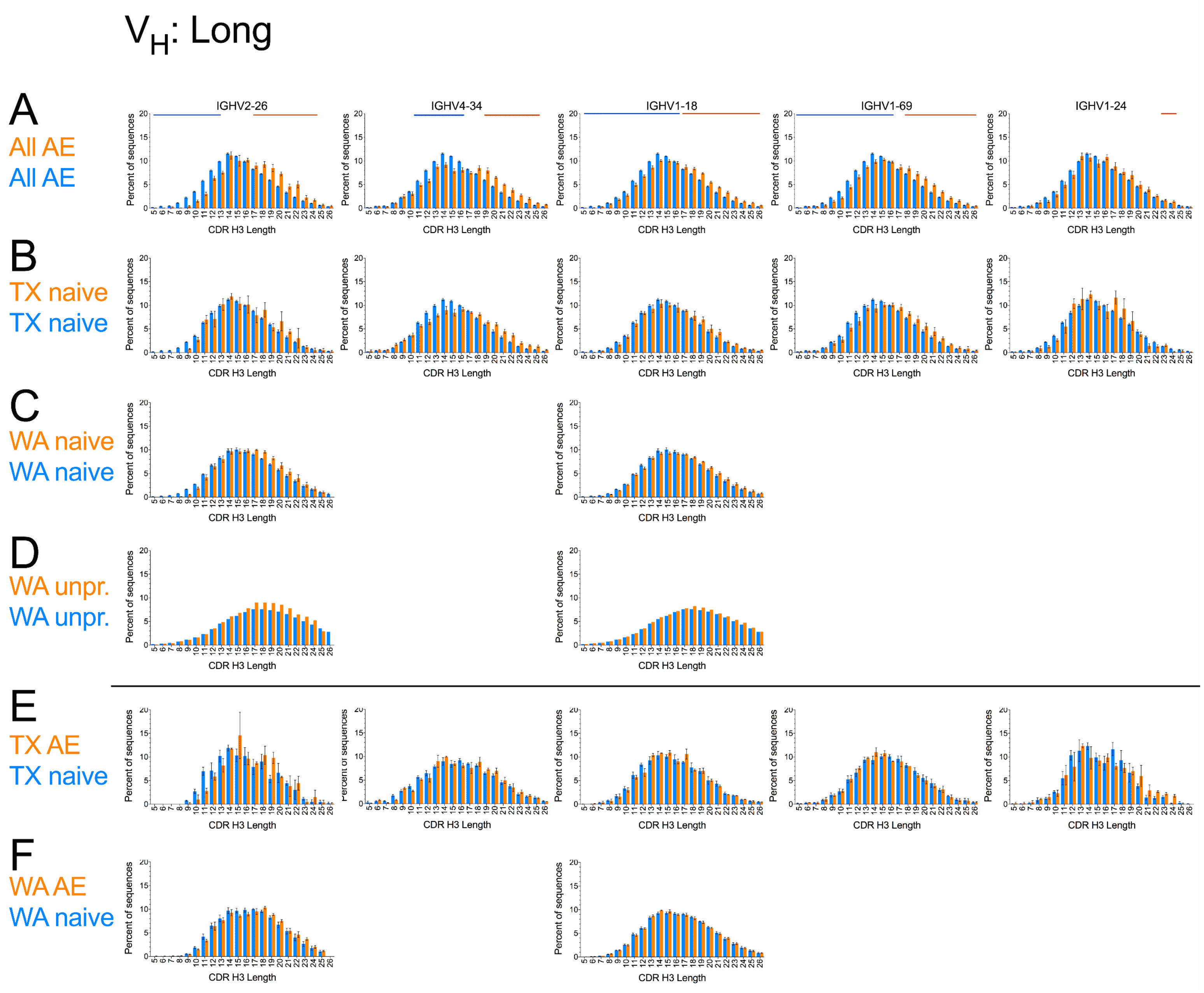

VH: Long
A
All AE
All AE
IGHV2-26
IGHV4-34
IGHV1-18
IGHV1-69
IGHV1-24
B
TX naive
TX naive
C
WA naive
WA naive
D
WA unpr.
WA unpr.
E
TX AE
TX naive
F
WA AE
WA naive
Percent of sequences
CDR H3 Length

### $V_H$: Neutral

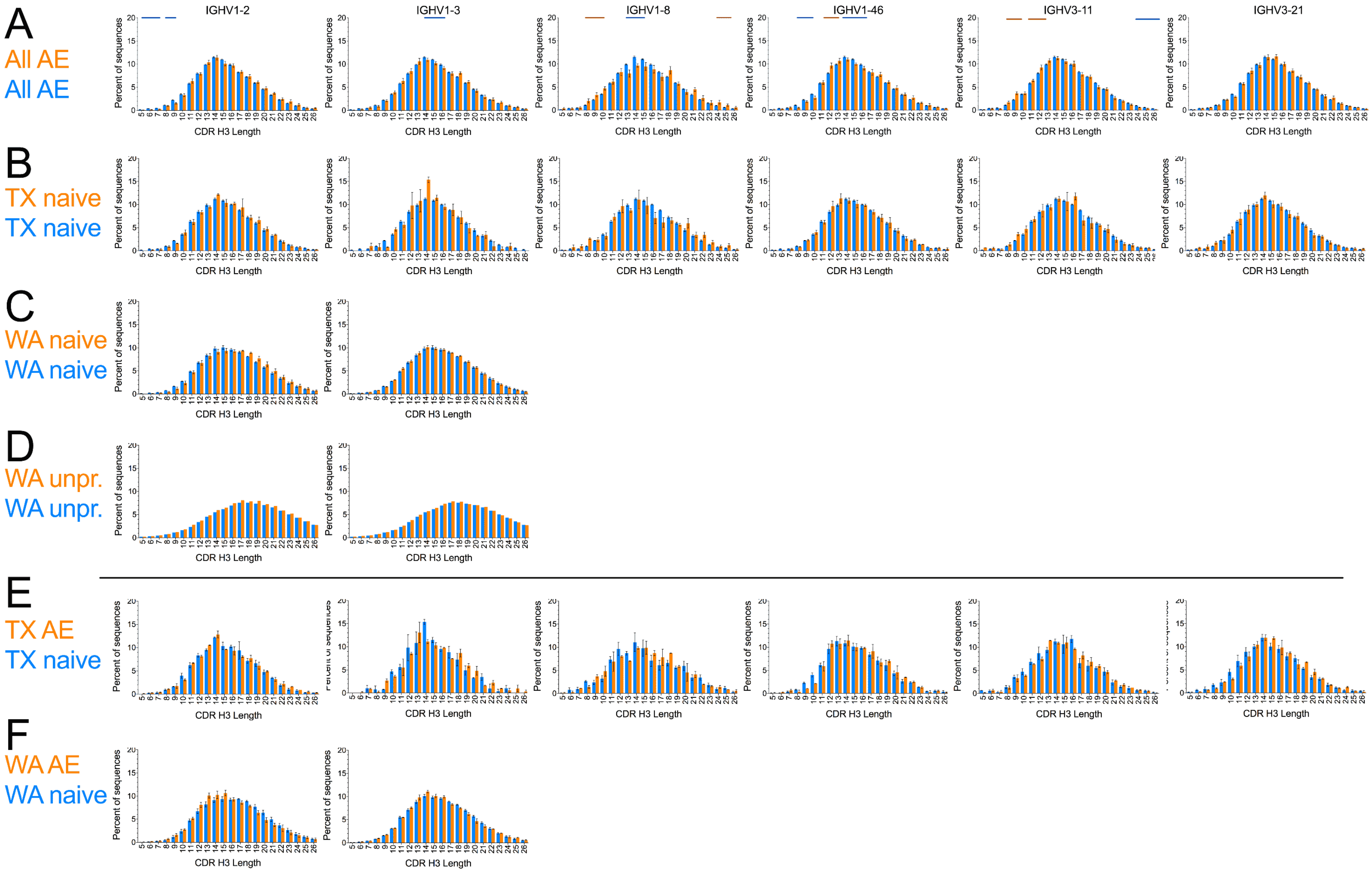
A
All AE
All AE
B
TX naive
TX naive
C
WA naive
WA naive
D
WA unpr.
WA unpr.
E
TX AE
TX naive
F
WA AE
WA naive
IGHV1-2
IGHV1-3
IGHV1-8
IGHV1-46
IGHV3-11
IGHV3-21
Percent of sequences
CDR H3 Length

$V_H$: Neutral

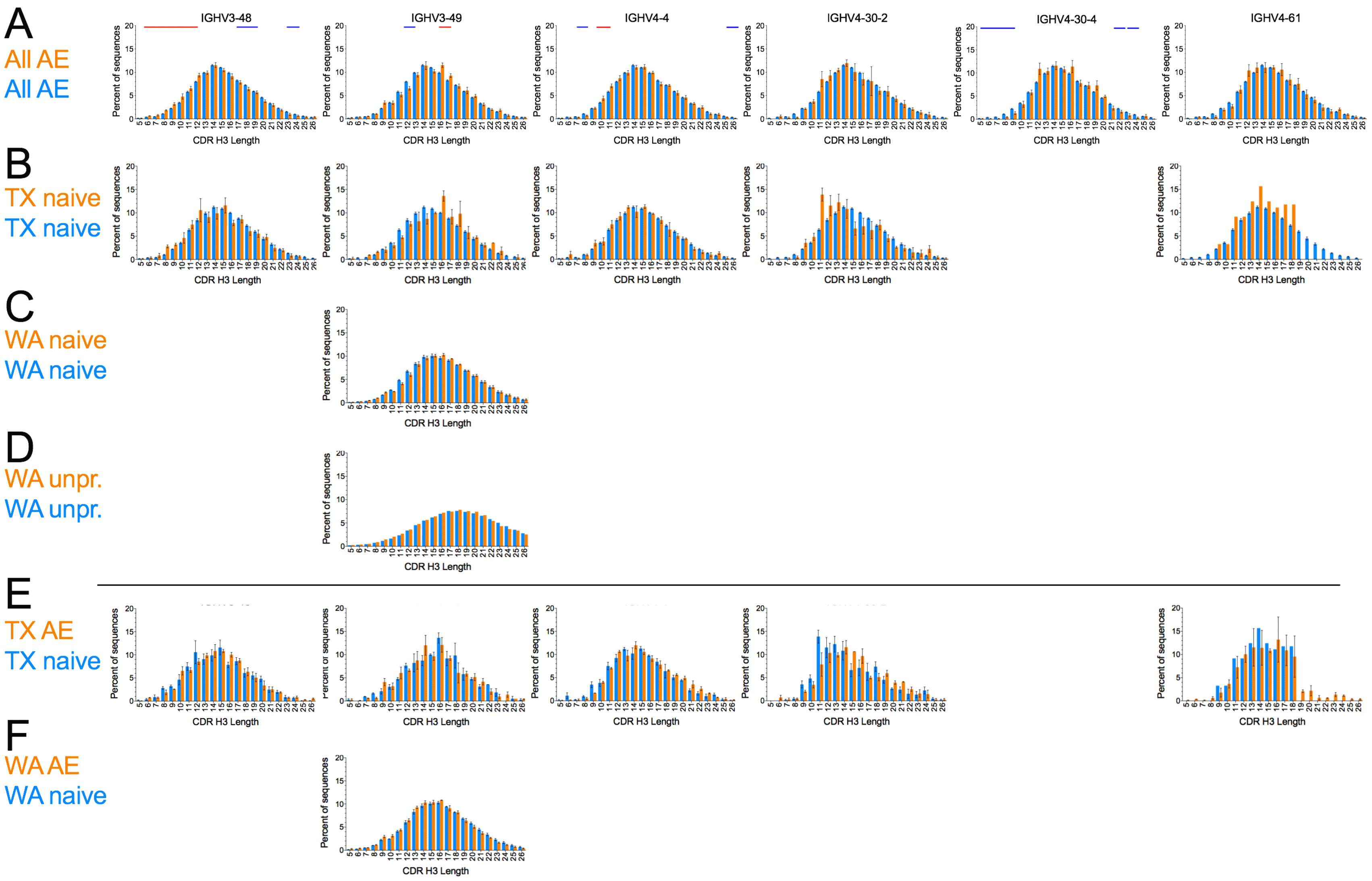

A
All AE
All AE
IGHV3-48
IGHV3-49
IGHV4-4
IGHV4-30-2
IGHV4-30-4
IGHV4-61
B
TX naive
TX naive
C
WA naive
WA naive
D
WA unpr.
WA unpr.
E
TX AE
TX naive
F
WA AE
WA naive
Percent of sequences
CDR H3 Length

### $V_H$: Neutral

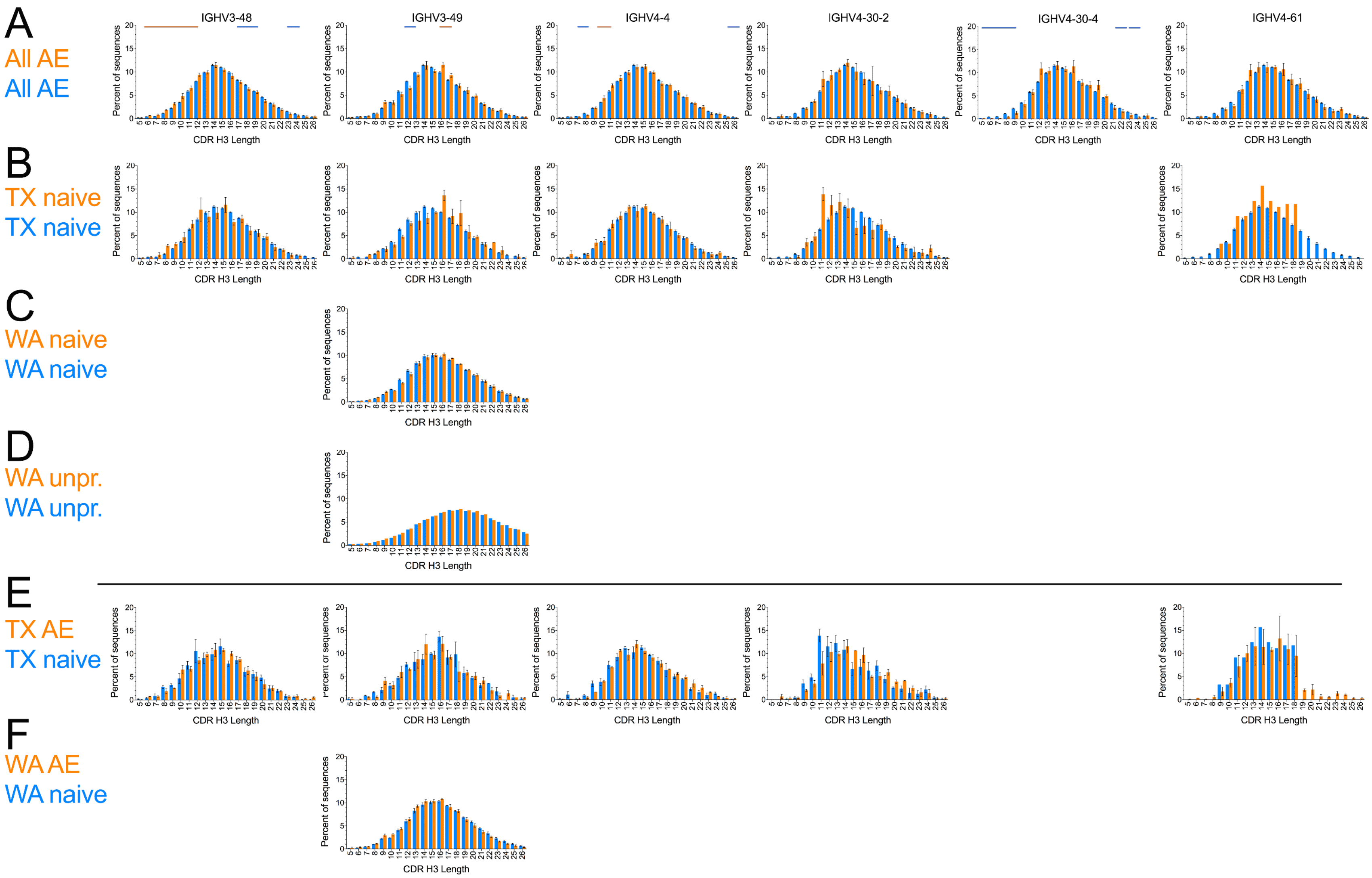

A
All AE
All AE
B
TX naive
TX naive
C
WA naive
WA naive
D
WA unpr.
WA unpr.
E
TX AE
TX naive
F
WA AE
WA naive
IGHV3-48
IGHV3-49
IGHV4-4
IGHV4-30-2
IGHV4-30-4
IGHV4-61
Percent of sequences
CDR H3 Length

### $V_H$: Neutral

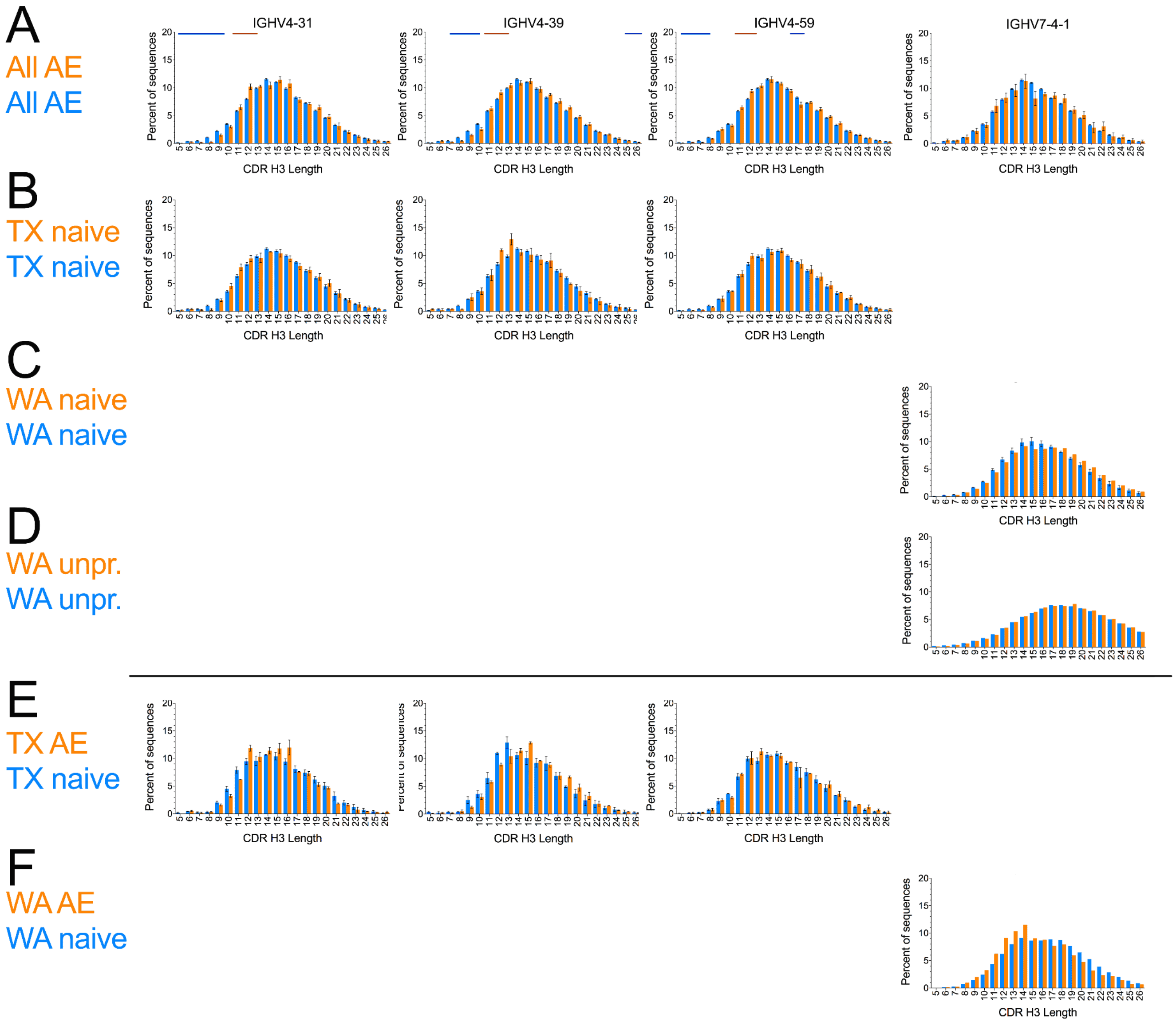
A
All AE
All AE
IGHV4-31
IGHV4-39
IGHV4-59
IGHV7-4-1
B
TX naive
TX naive
C
WA naive
WA naive
D
WA unpr.
WA unpr.
E
TX AE
TX naive
F
WA AE
WA naive
Percent of sequences
CDR H3 Length

**Fig. S2.** CDR H3 length distributions associated with $V_H$ germlines (orange bars) compared to overall distribution in the same samples and compartments (A-D, blue bars) or to corresponding naïve sequences (E and F):

(A) AE and/or IgG/IgA sequences, all donors pooled

(B) TX $CD27^{neg}$ (naïve) compartment

(C) WA $CD27^{neg}$ (naïve) compartment

(D) WA $V_H$ unproductive sequences

(E) TX IgG/IgA (orange bars, same as orange bars in Suppl. Fig. 3, TX) and naïve sequences (blue bars, same as orange bars B)

(F) WA $CD27^{pos}$ (orange bars, same as orange bars in Suppl. Fig. 3, WA) and naïve sequences (blue bars, same as orange bars C)

Red and blue horizontal bars above the histograms in (A) indicate statistically significant ($P < 10^{-4}$) differences between the germline-specific and overall distributions in a paired *t*-test (see main text for details). Error bars indicate S.E.M. (A, B, D, E and F) or range (panel C). Arrows indicate subtle deviations in the distributions. Germlines are noted in the IMGT naming convention.

### $V_H$: Short

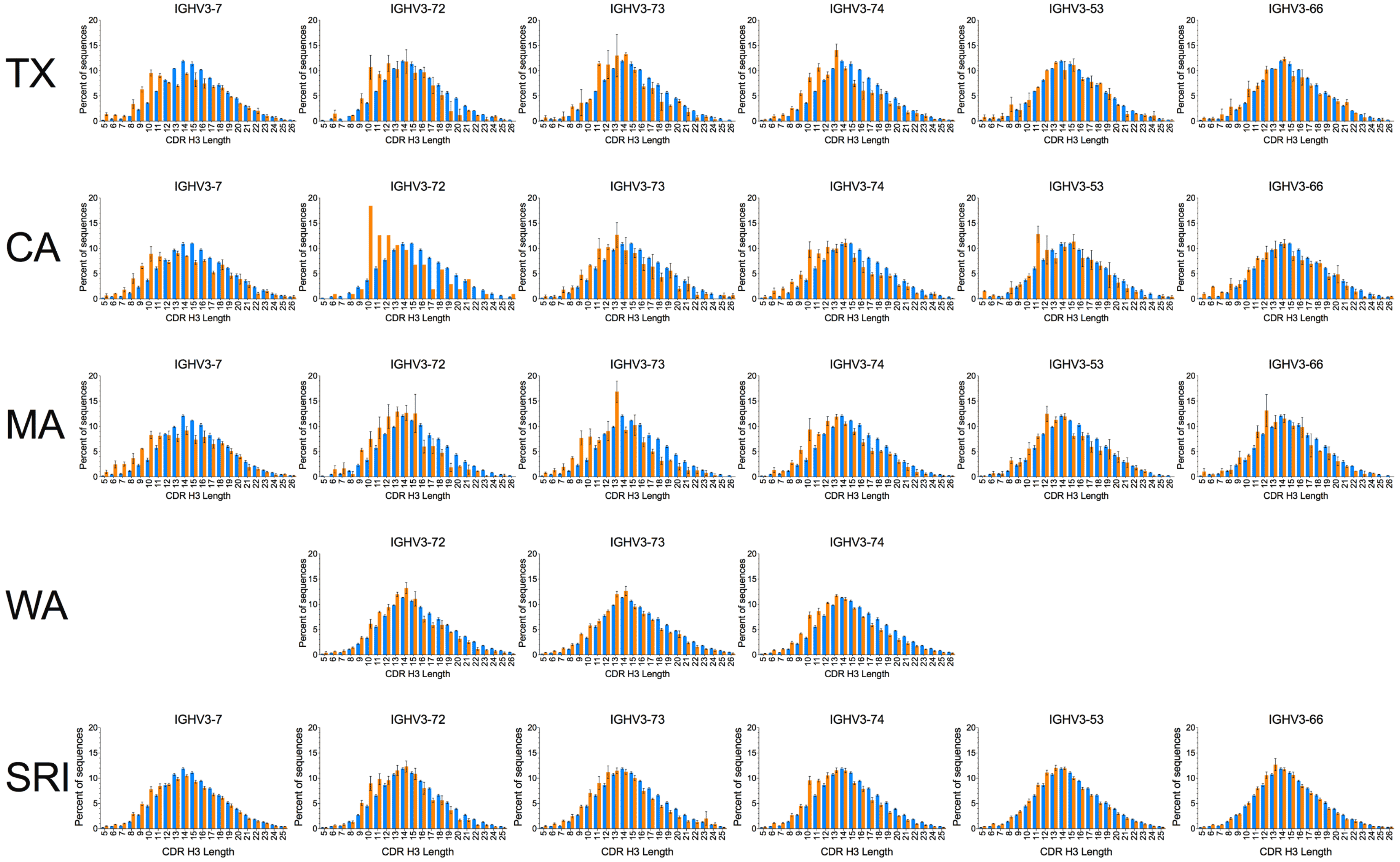
TX
IGHV3-7
IGHV3-72
IGHV3-73
IGHV3-74
IGHV3-53
IGHV3-66
CA
IGHV3-7
IGHV3-72
IGHV3-73
IGHV3-74
IGHV3-53
IGHV3-66
MA
IGHV3-7
IGHV3-72
IGHV3-73
IGHV3-74
IGHV3-53
IGHV3-66
WA
IGHV3-72
IGHV3-73
IGHV3-74
SRI
IGHV3-7
IGHV3-72
IGHV3-73
IGHV3-74
IGHV3-53
IGHV3-66
Percent of sequences
CDR H3 Length

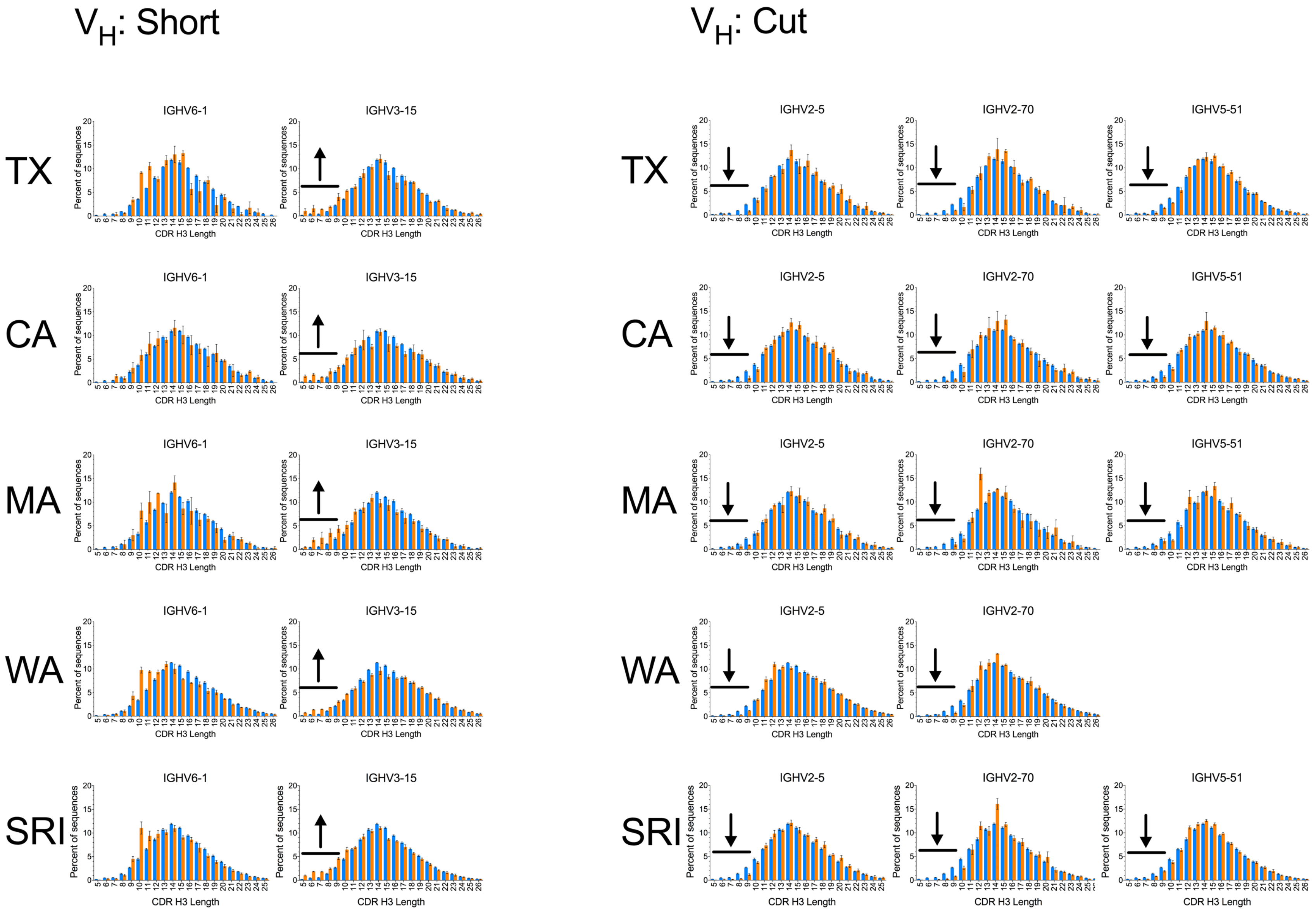

$V_H$: Short
$V_H$: Cut
TX
CA
MA
WA
SRI
IGHV6-1
IGHV3-15
IGHV2-5
IGHV2-70
IGHV5-51
Percent of sequences
CDR H3 Length

### $V_H$: Crested

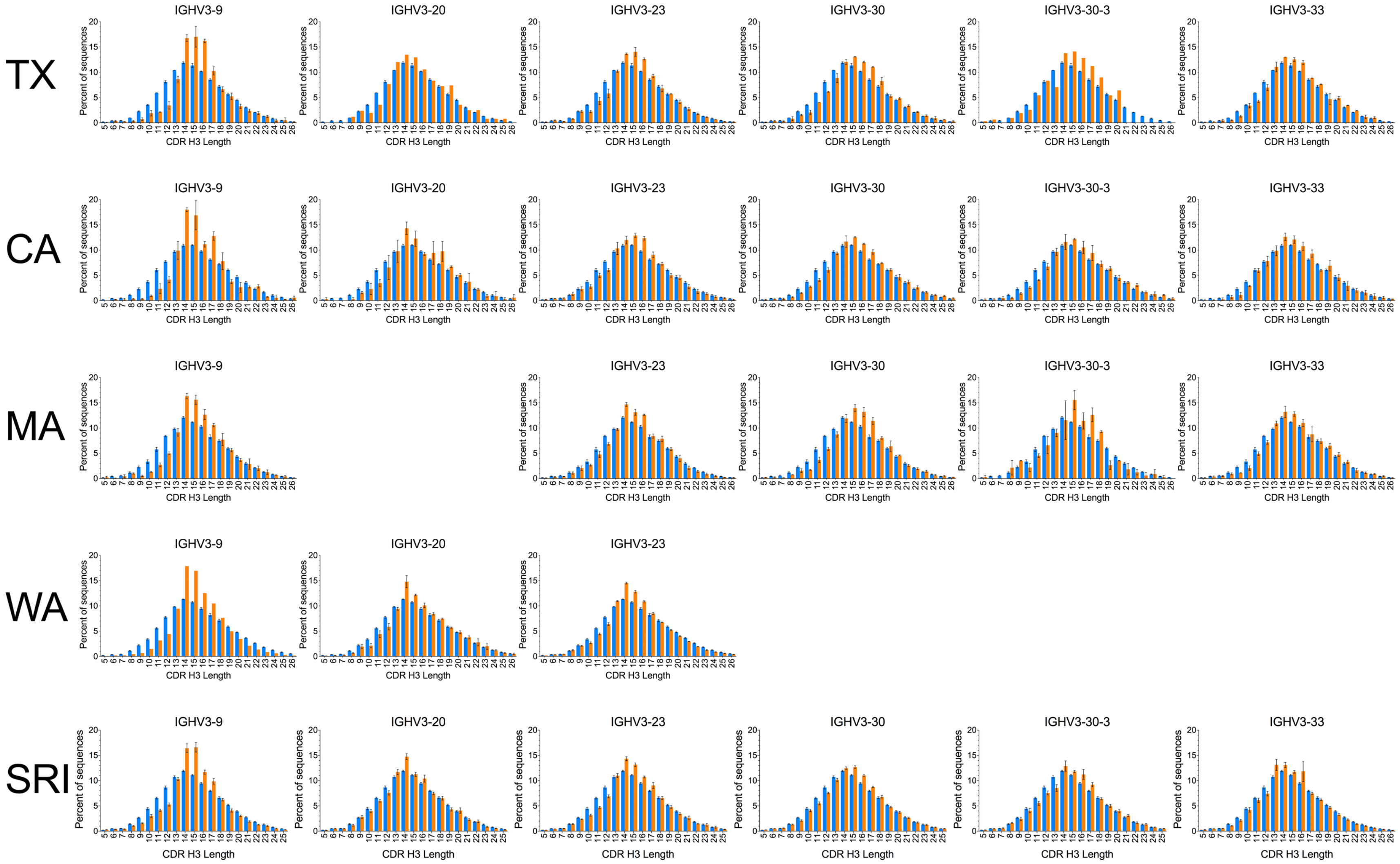
TX
CA
MA
WA
SRI
IGHV3-9
IGHV3-20
IGHV3-23
IGHV3-30
IGHV3-30-3
IGHV3-33
Percent of sequences
CDR H3 Length

### $V_H$: Crested

TX

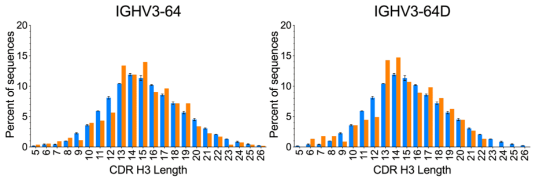


CA

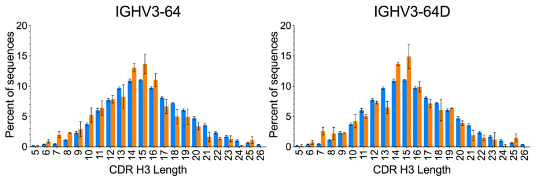


MA

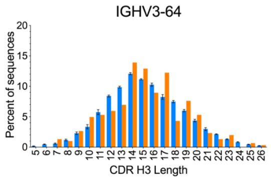


SRI

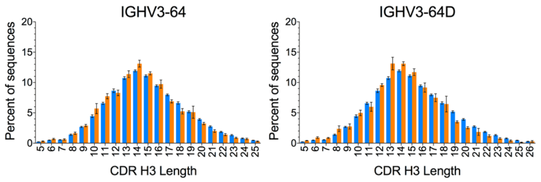

### $V_H$: Long

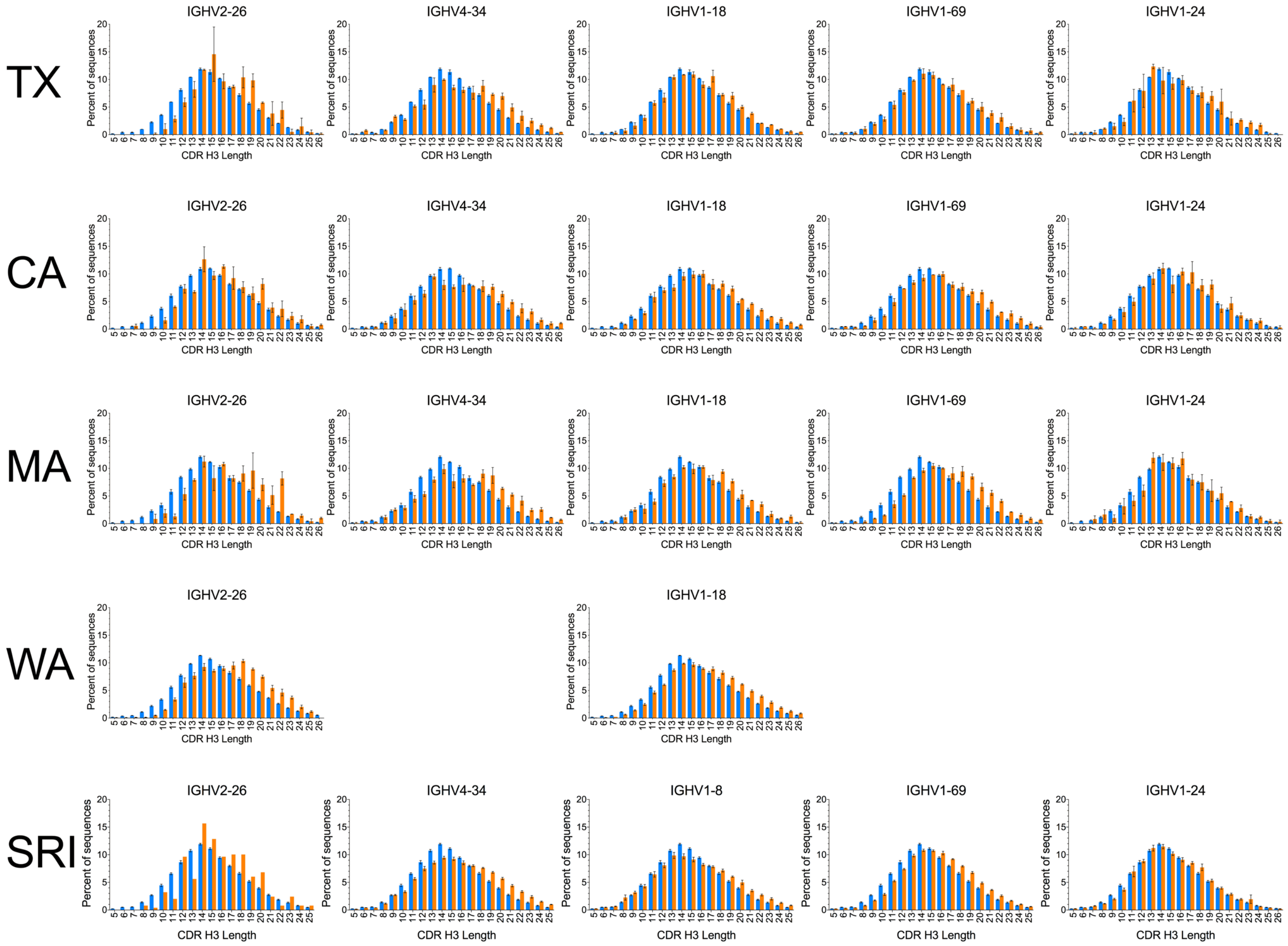
TX
IGHV2-26
IGHV4-34
IGHV1-18
IGHV1-69
IGHV1-24
CA
IGHV2-26
IGHV4-34
IGHV1-18
IGHV1-69
IGHV1-24
MA
IGHV2-26
IGHV4-34
IGHV1-18
IGHV1-69
IGHV1-24
WA
IGHV2-26
IGHV1-18
SRI
IGHV2-26
IGHV4-34
IGHV1-8
IGHV1-69
IGHV1-24
Percent of sequences
CDR H3 Length

### $V_H$: Neutral

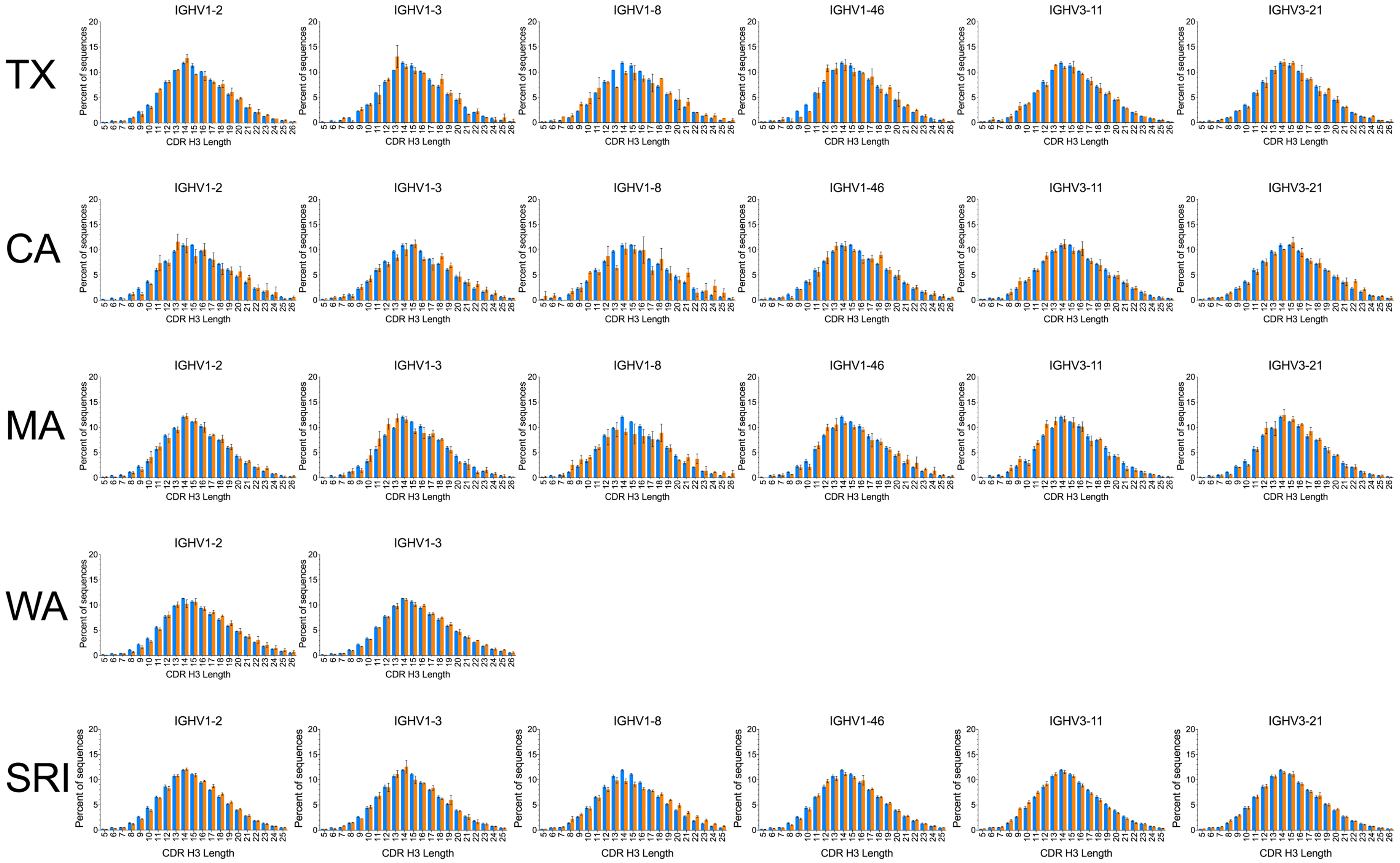
TX
CA
MA
WA
SRI
IGHV1-2
IGHV1-3
IGHV1-8
IGHV1-46
IGHV3-11
IGHV3-21
Percent of sequences
CDR H3 Length

#### $V_H$: Neutral

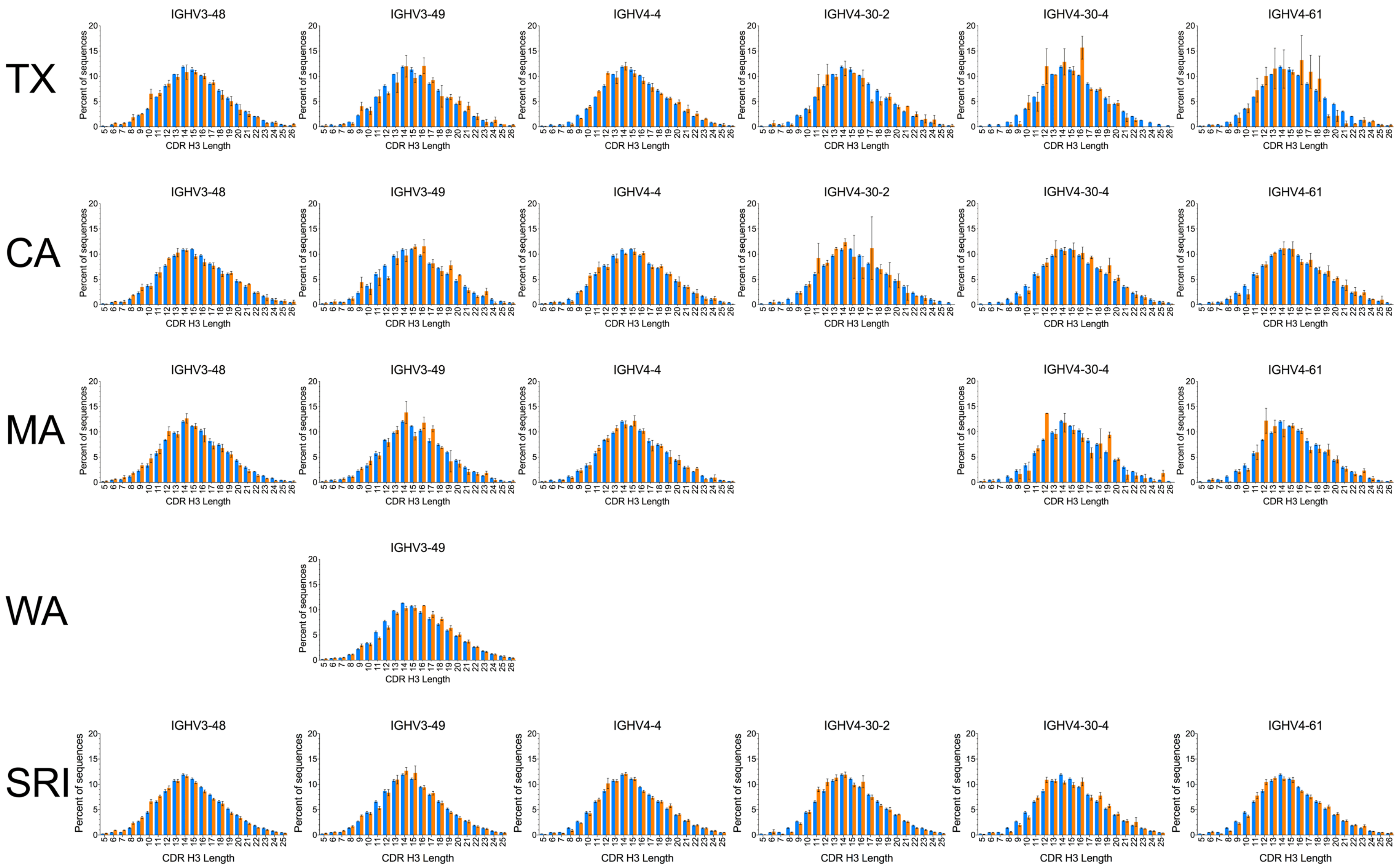

TX
CA
MA
WA
SRI
IGHV3-48
IGHV3-49
IGHV4-4
IGHV4-30-2
IGHV4-30-4
IGHV4-61
Percent of sequences
CDR H3 Length

### $V_H$: Neutral

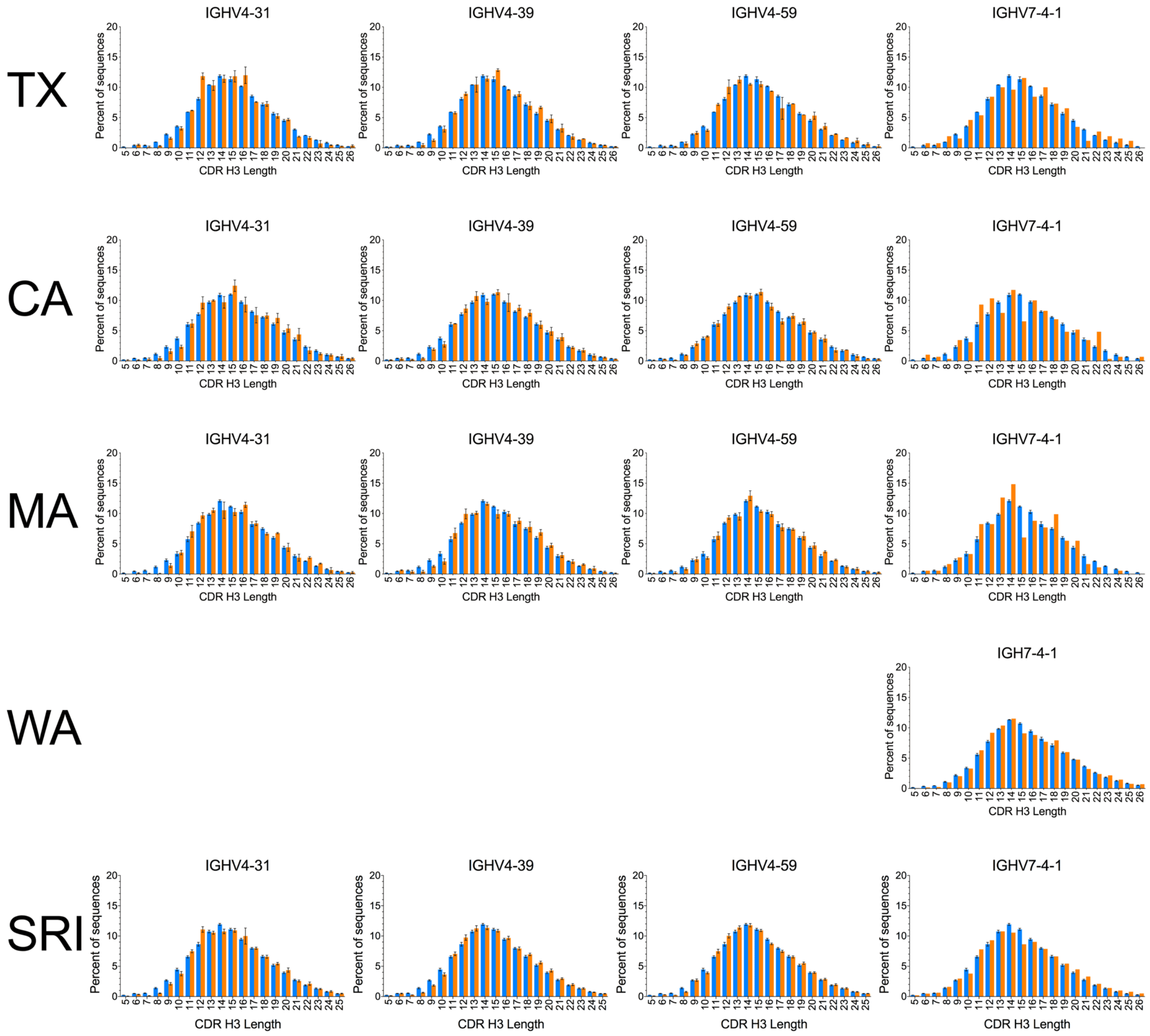
TX
CA
MA
WA
SRI
IGHV4-31
IGHV4-39
IGHV4-59
IGHV7-4-1
IGH7-4-1
Percent of sequences
CDR H3 Length

**Fig. S3.** Comparability of $V_H$-associated CDR H3 length biases among datasets. $V_H$ germline-specific CDR H3 length distributions (orange bars) compared to overall distribution (blue bars) in the AE and/or isotype-switched B cell compartments of the five datasets. Error bars indicate S.E.M. Arrows indicate subtle deviations in the distributions shared among datasets. Germlines are noted in the IMGT naming convention.

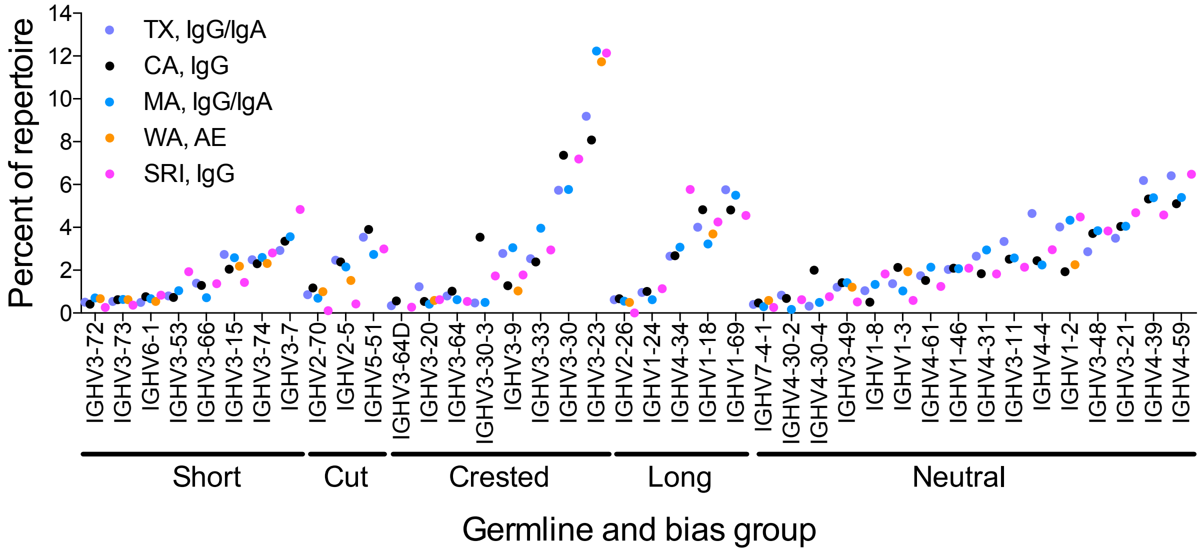


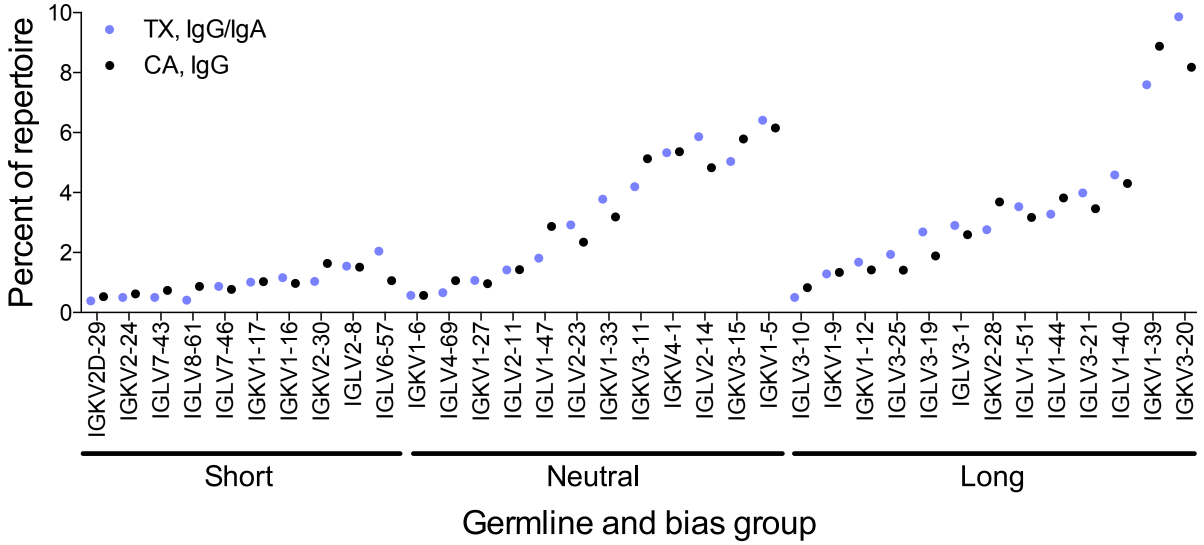


**Fig. S4.** Prevalence of $V_H$ (B) and $V_L$ (B) germlines in the AE compartment. Prevalence of germlines was calculated based on unique clonotypes.

A

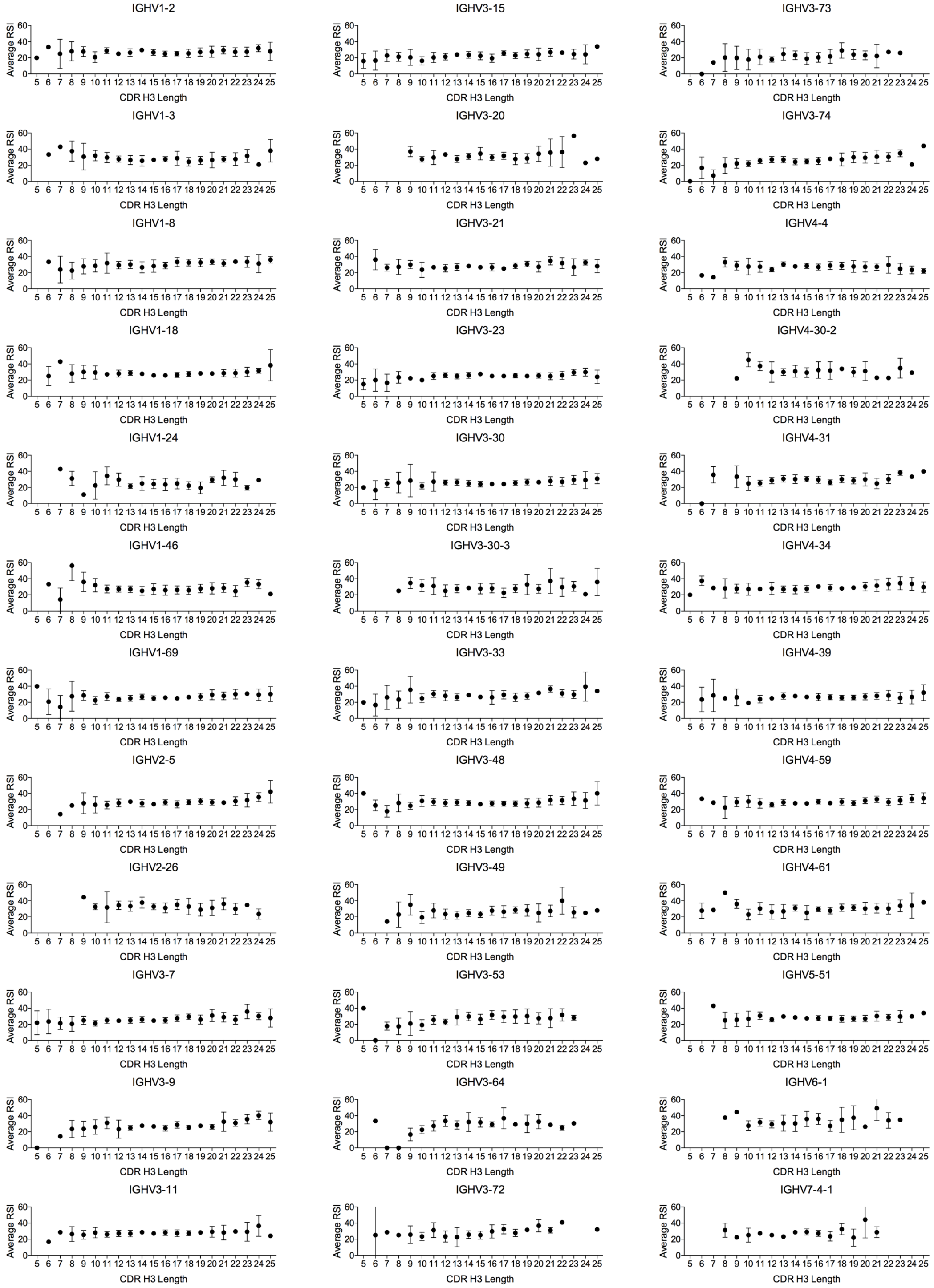
IGHV1-2
IGHV3-15
IGHV3-73
IGHV1-3
IGHV3-20
IGHV3-74
IGHV1-8
IGHV3-21
IGHV4-4
IGHV1-18
IGHV3-23
IGHV4-30-2
IGHV1-24
IGHV3-30
IGHV4-31
IGHV1-46
IGHV3-30-3
IGHV4-34
IGHV1-69
IGHV3-33
IGHV4-39
IGHV2-5
IGHV3-48
IGHV4-59
IGHV2-26
IGHV3-49
IGHV4-61
IGHV3-7
IGHV3-53
IGHV5-51
IGHV3-9
IGHV3-64
IGHV6-1
IGHV3-11
IGHV3-72
IGHV7-4-1
Average RSI
CDR H3 Length

B

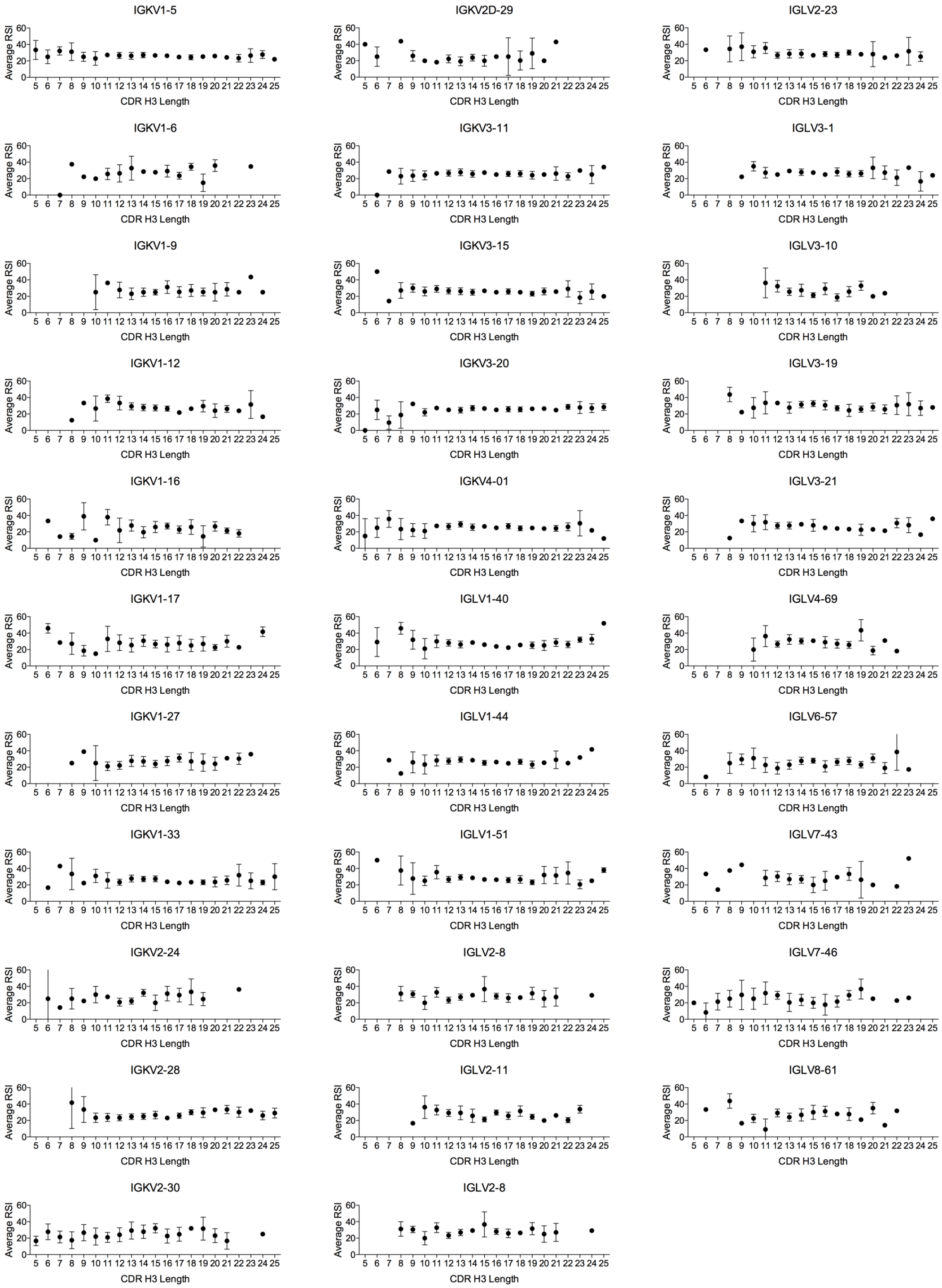


**Fig. S5.** RSI analysis of $V_H$ (A) and $V_L$ (B) germline-associated sequences. Points show average and standard deviation of the RSI value for sequences from each donor, germline and CDR H3 length. WA and SRI sequences were excluded due to large dataset sizes. Note that RSI values cannot be higher than 60 in the TX, CA and MA datasets due to clonotype definition. Germlines are noted in the IMGT naming convention.

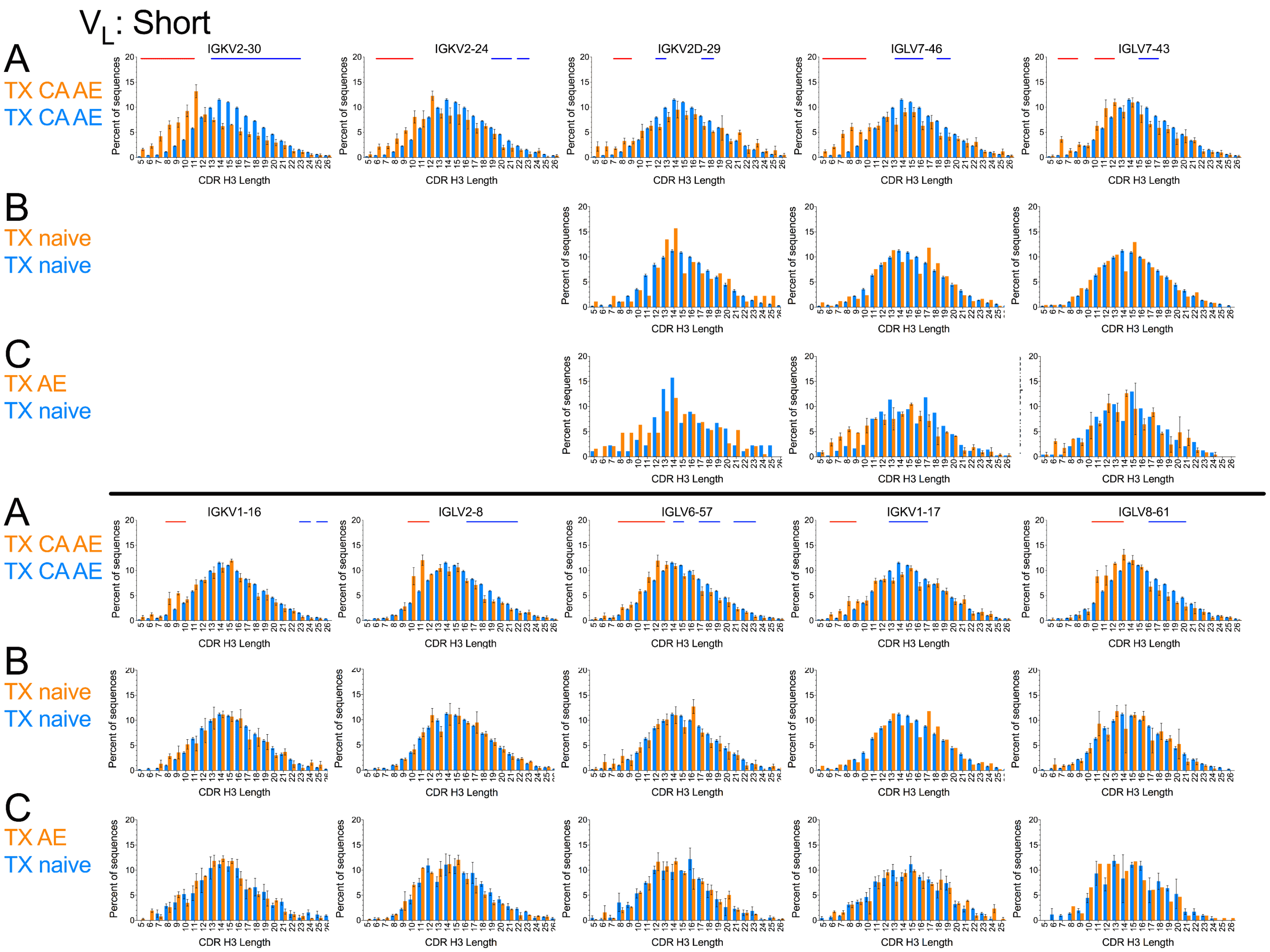
V$_L$: Short
A
TX CA AE
TX CA AE
B
TX naive
TX naive
C
TX AE
TX naive
IGKV2-30
IGKV2-24
IGKV2D-29
IGLV7-46
IGLV7-43
A
TX CA AE
TX CA AE
B
TX naive
TX naive
C
TX AE
TX naive
IGKV1-16
IGLV2-8
IGLV6-57
IGKV1-17
IGLV8-61
Percent of sequences
CDR H3 Length

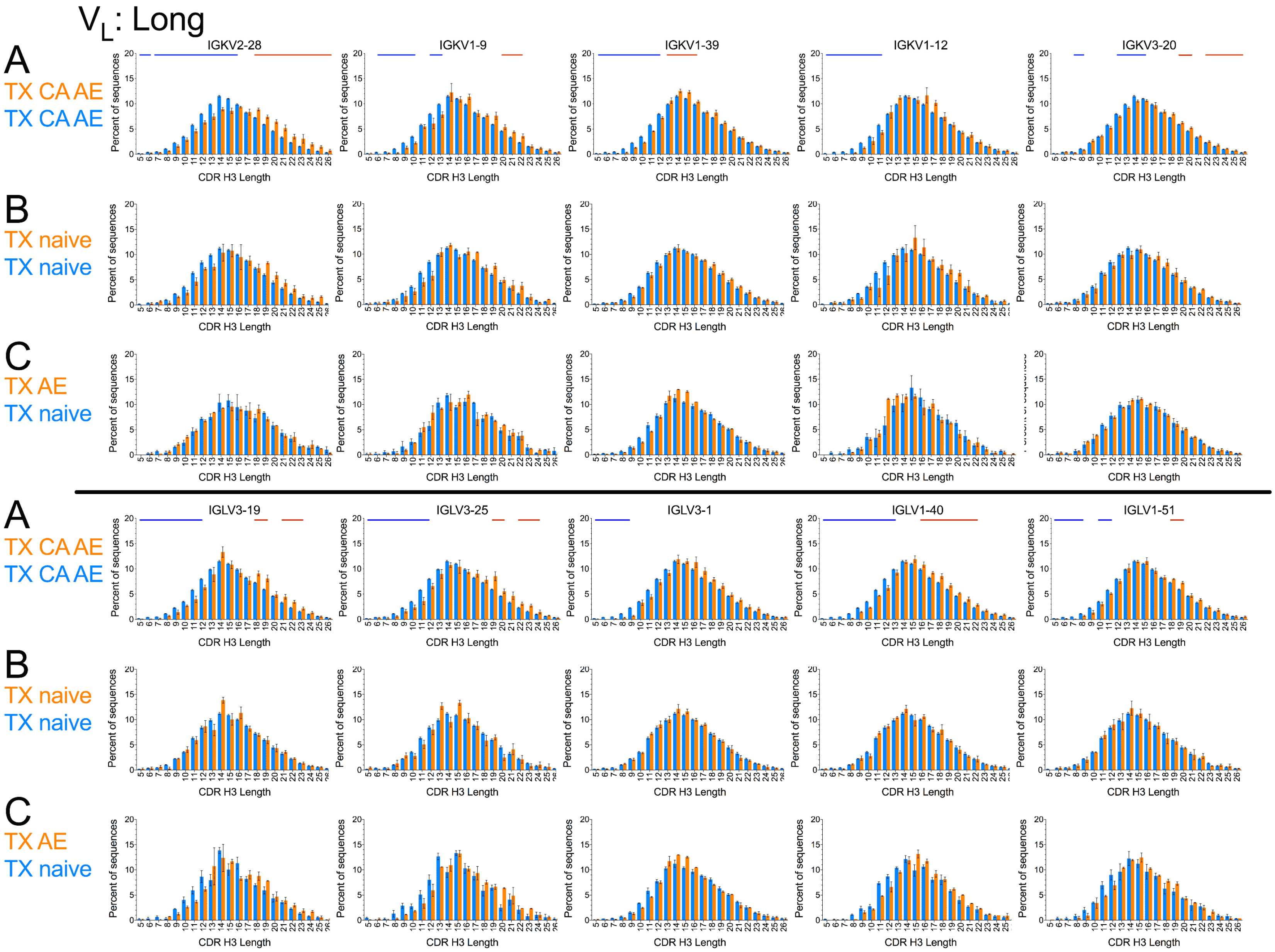
V$_L$: Long
A
TX CA AE
TX CA AE
B
TX naive
TX naive
C
TX AE
TX naive
IGKV2-28
IGKV1-9
IGKV1-39
IGKV1-12
IGKV3-20
IGLV3-19
IGLV3-25
IGLV3-1
IGLV1-40
IGLV1-51
Percent of sequences
CDR H3 Length

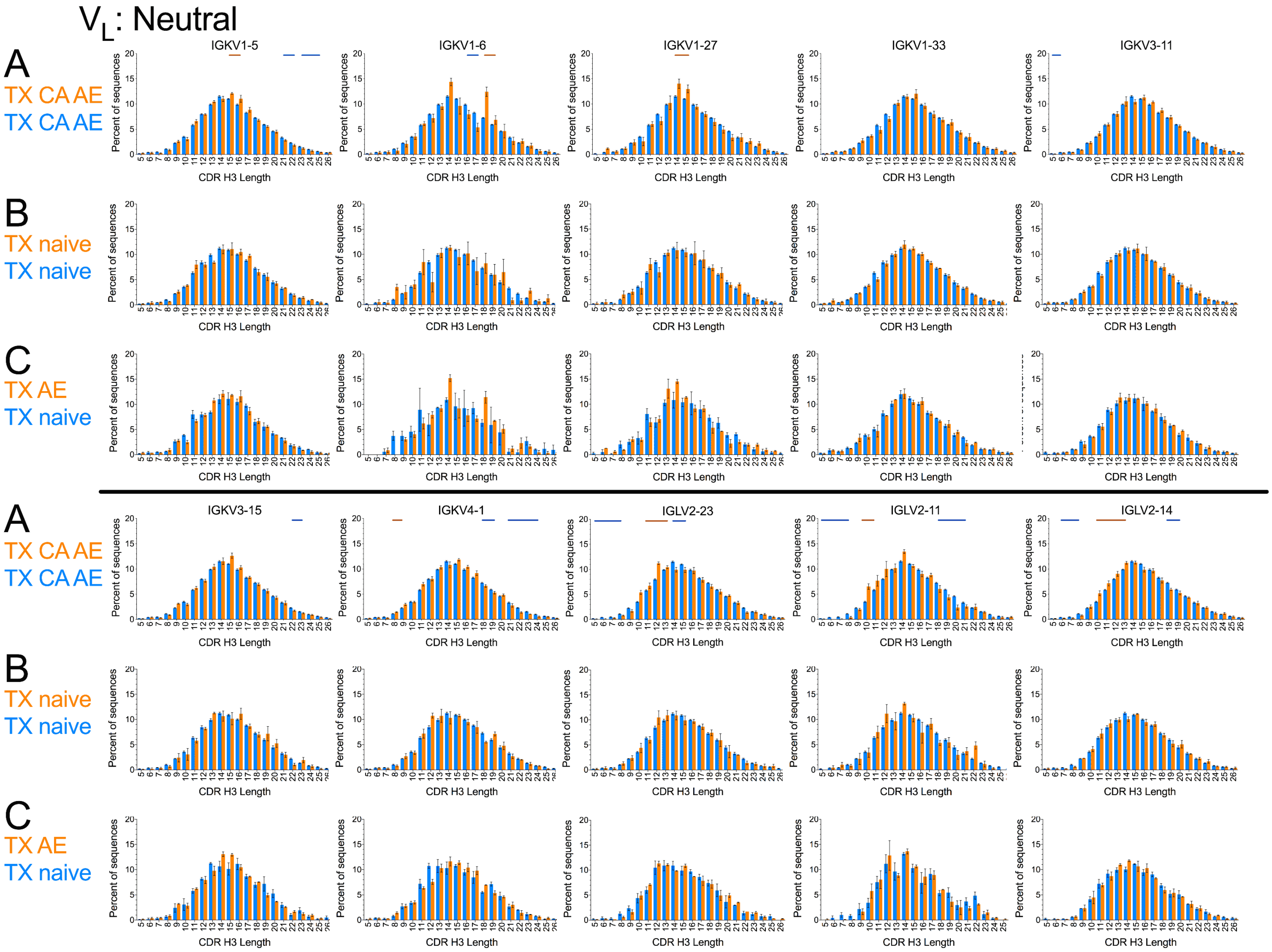

VL: Neutral
A
TX CA AE
TX CA AE
B
TX naive
TX naive
C
TX AE
TX naive
IGKV1-5
IGKV1-6
IGKV1-27
IGKV1-33
IGKV3-11
IGKV3-15
IGKV4-1
IGLV2-23
IGLV2-11
IGLV2-14
Percent of sequences
CDR H3 Length

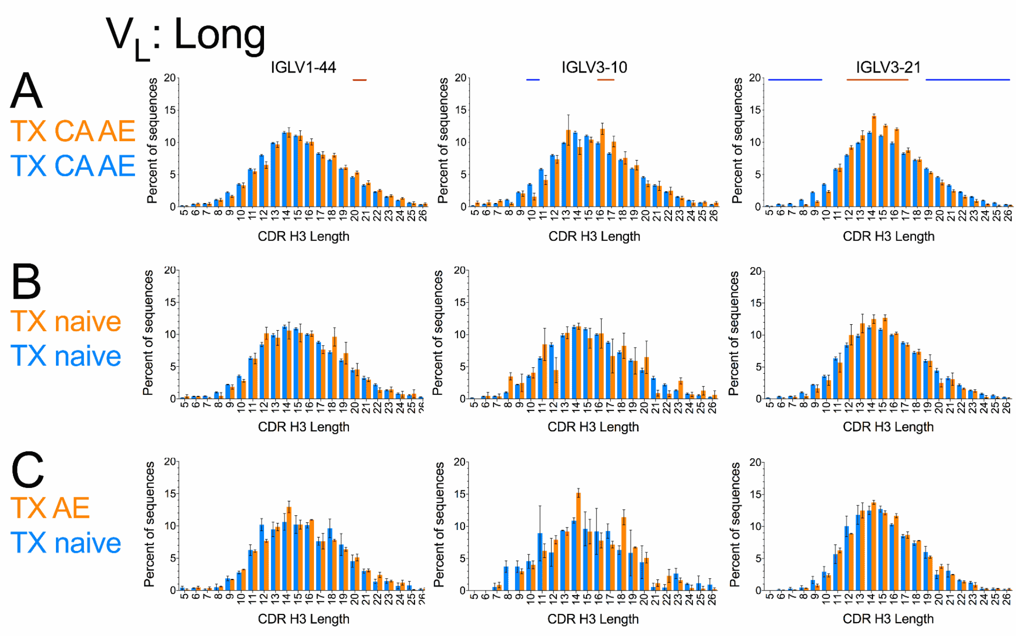


**Fig. S6.** CDR H3 length distributions associated with $V_L$ germlines (orange bars) compared to overall distribution in the same samples and compartments (A and B, blue bars) or to corresponding naïve sequences (C):

(A) AE and/or IgG/IgA sequences, CA and TX donors pooled

(B) TX CD27$^{neg}$ (naïve) compartment

(C) TX IgG/IgA (orange bars, same as orange bars in Suppl. Fig. 6 TX) and naïve (blue bars, same as orange bars in B) sequences of the indicated germline

Red and blue horizontal bars above the histograms in (A) indicate statistically significant ($P < 10^{-4}$) differences between the germline-specific and overall distributions in a paired *t*-test with a rolling window of 2 consecutive CDR H3 lengths (see main text for details). Error bars indicate S.E.M. (A and B) or range (C). Arrows indicate subtle deviations in the distributions. Germlines are noted in the IMGT naming convention.

#### $V_L$: Short

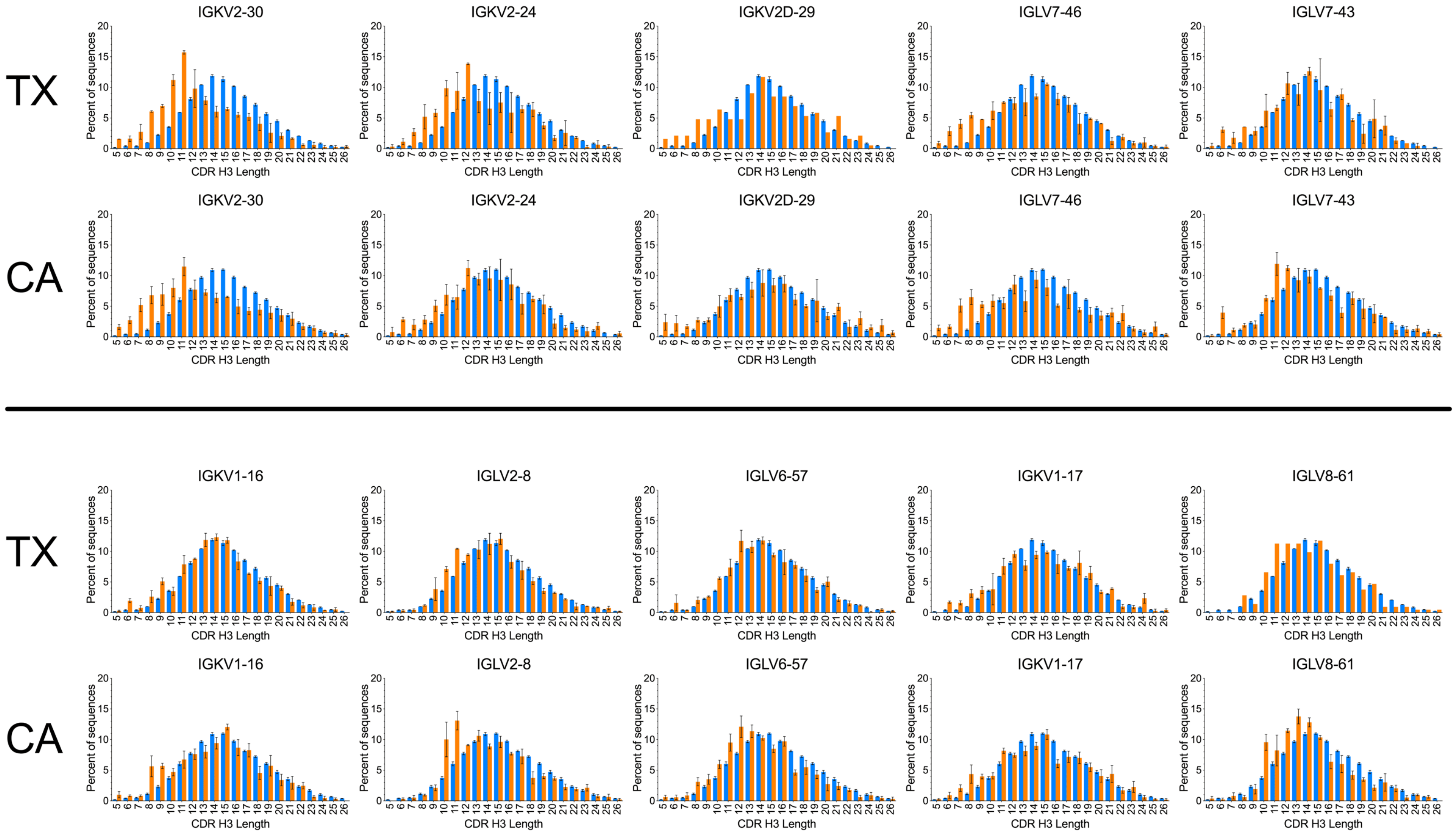

TX
CA
IGKV2-30
IGKV2-24
IGKV2D-29
IGLV7-46
IGLV7-43
IGKV1-16
IGLV2-8
IGLV6-57
IGKV1-17
IGLV8-61
Percent of sequences
CDR H3 Length

### $V_L$: Long

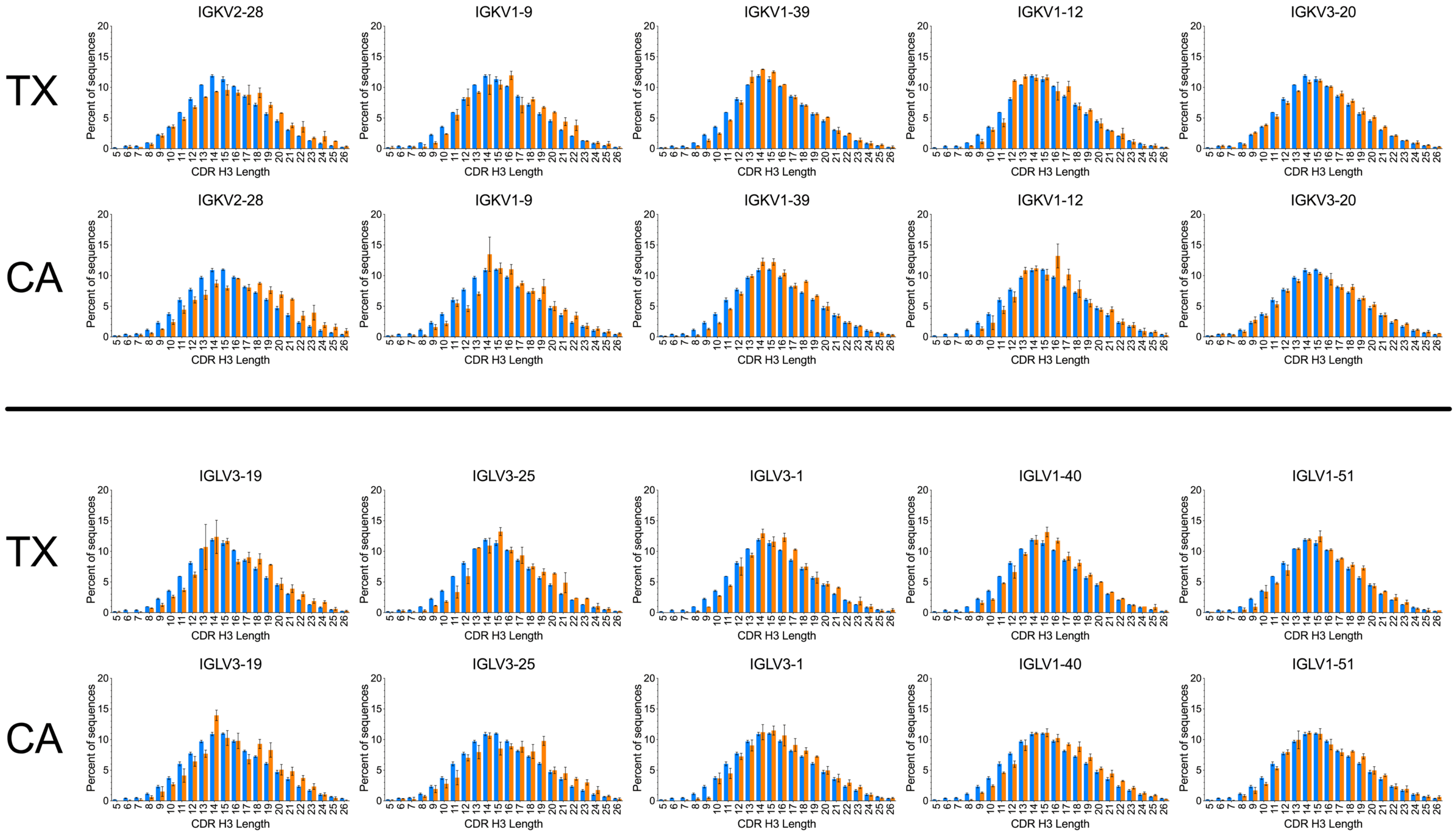
TX
CA
IGKV2-28
IGKV1-9
IGKV1-39
IGKV1-12
IGKV3-20
IGLV3-19
IGLV3-25
IGLV3-1
IGLV1-40
IGLV1-51
Percent of sequences
CDR H3 Length

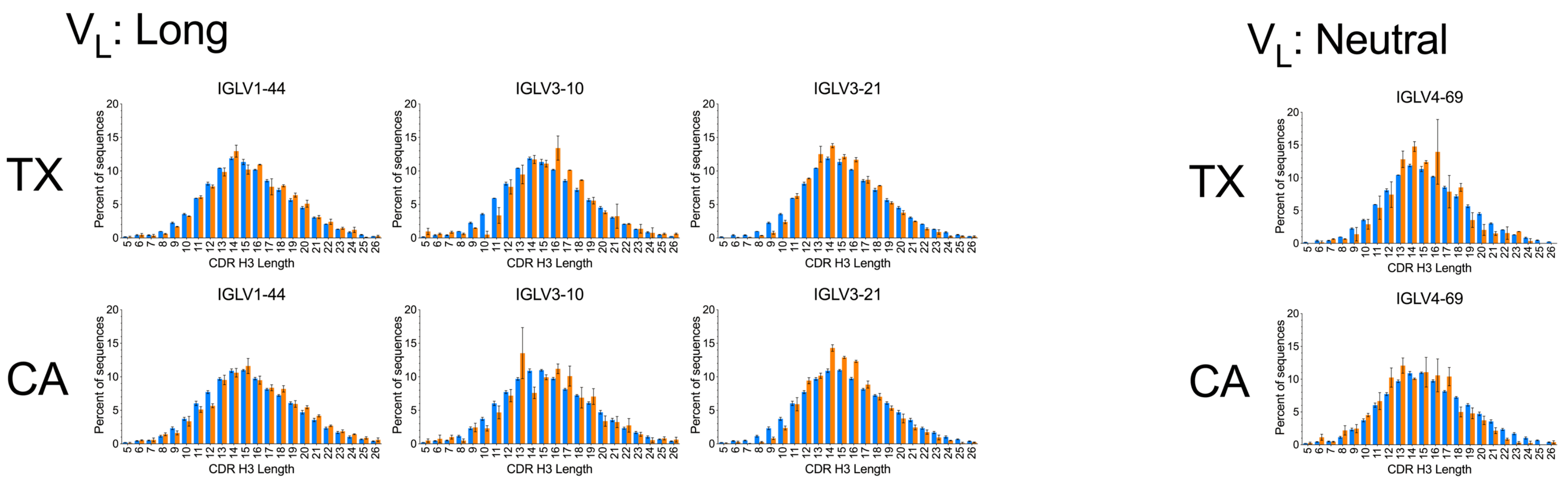

$V_L$: Long
$V_L$: Neutral
TX
CA
IGLV1-44
IGLV3-10
IGLV3-21
IGLV4-69
Percent of sequences
CDR H3 Length

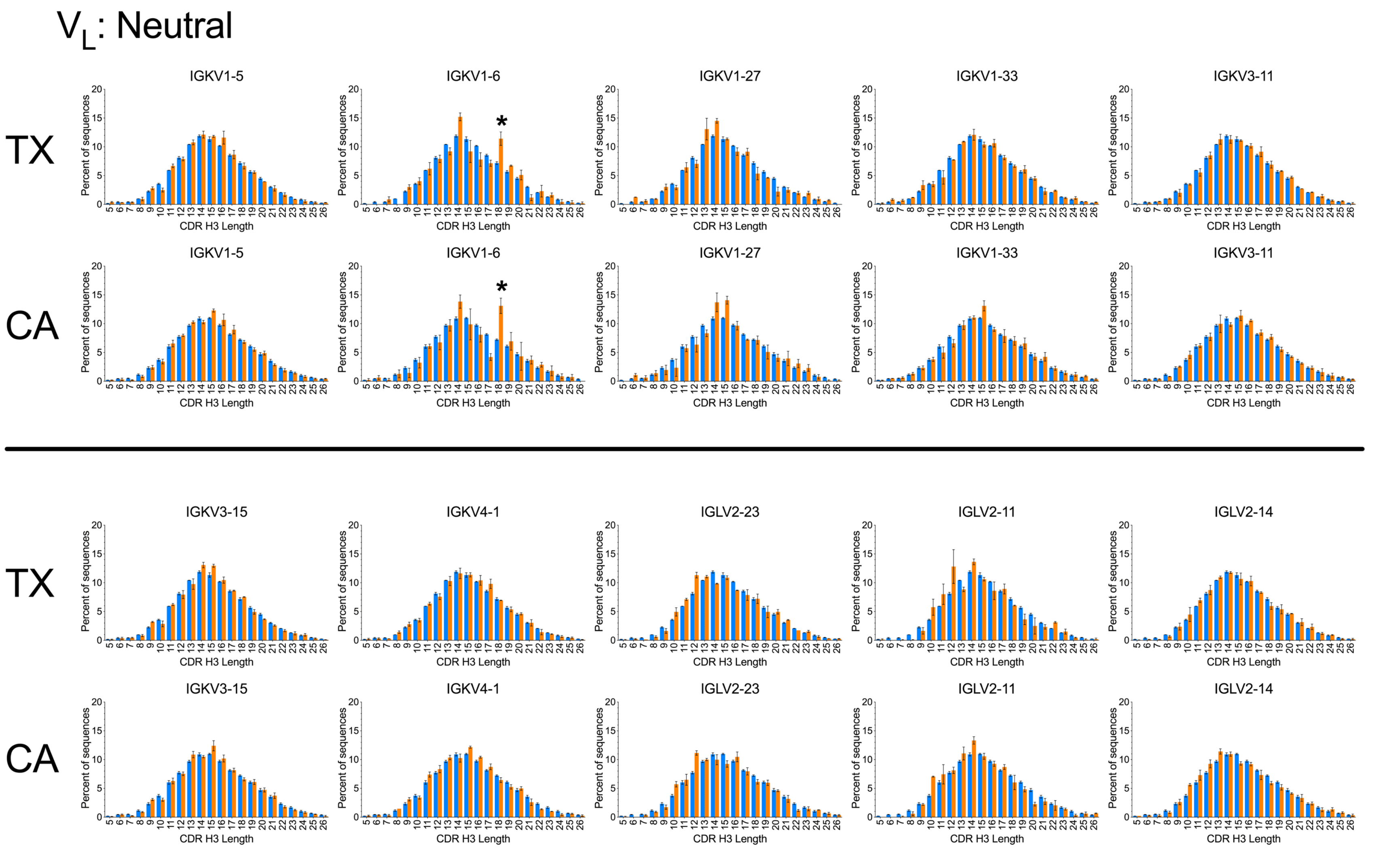


**Fig. S7.** $V_L$-associated CDR H3 length biases are comparable between the CA and TX (AE compartment) datasets. $V_L$ germline-specific CDR H3 length distributions (orange bars) compared to overall distribution (blue bars) in the four datasets. Error bars indicate S.E.M. (CA) and range (TX). Germlines are named in the IMGT convention. Asterisks indicate spikes at length 18 in the IGKV1-6 distributions of TX and CA datasets with high prevalence of convergent clones with IGHV3-7, IGHJ3, CDR H3 consensus sequence (A/V)RD$X_7$(L/I/V)(W/Y)YDAFDI and light chain Arg-116 in CDR L3, observed in all CA and TX donors.

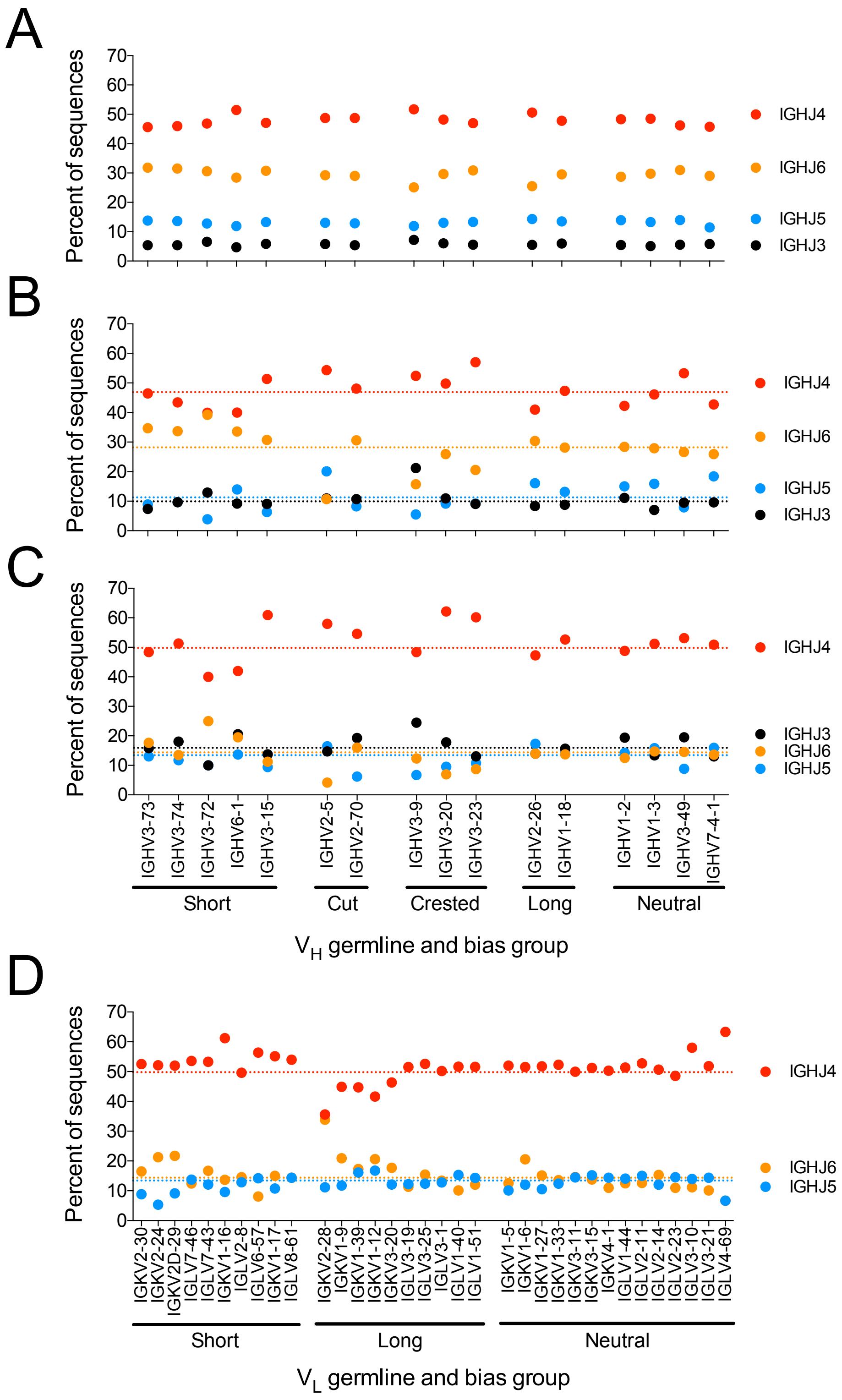


**Fig. S8.** $J_H$ prevalence as a function of $V_H$ (A-C) and $V_L$ (D) germline in WA unproductive sequences (A) and WA (B) and TX (C and D) naïve compartment sequences. Dotted lines indicate overall average $J_H$ prevalence. Germlines are noted in the IMGT naming convention.

WA: Unproductive

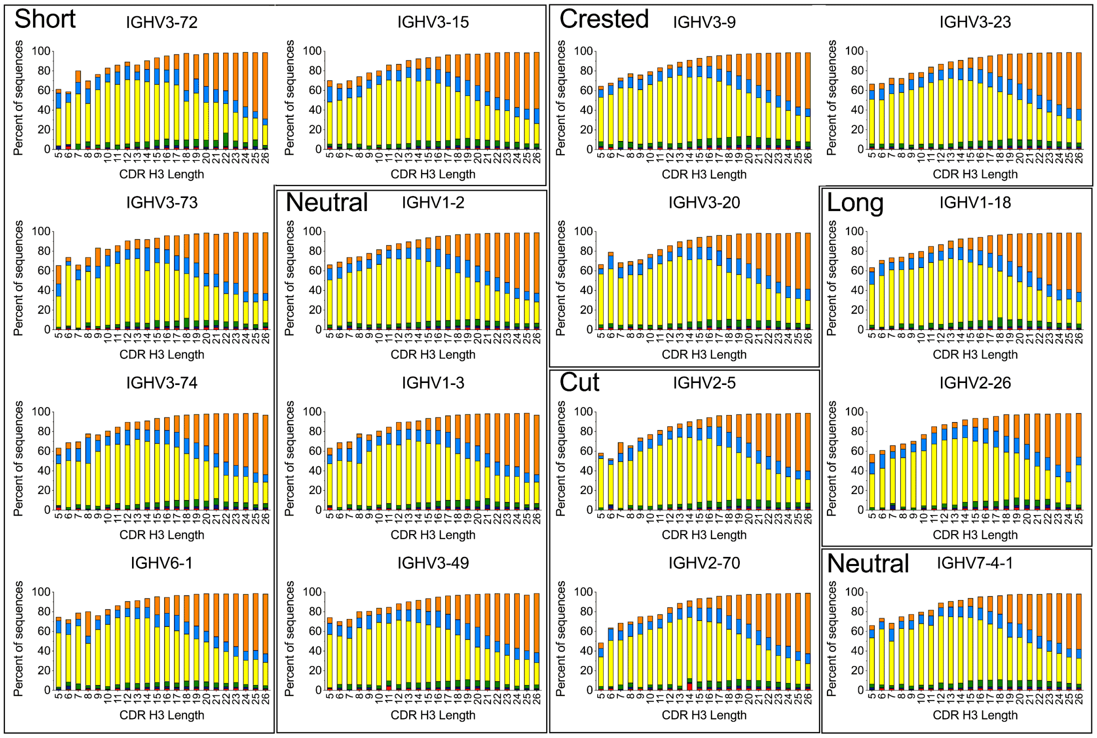


WA: Naive

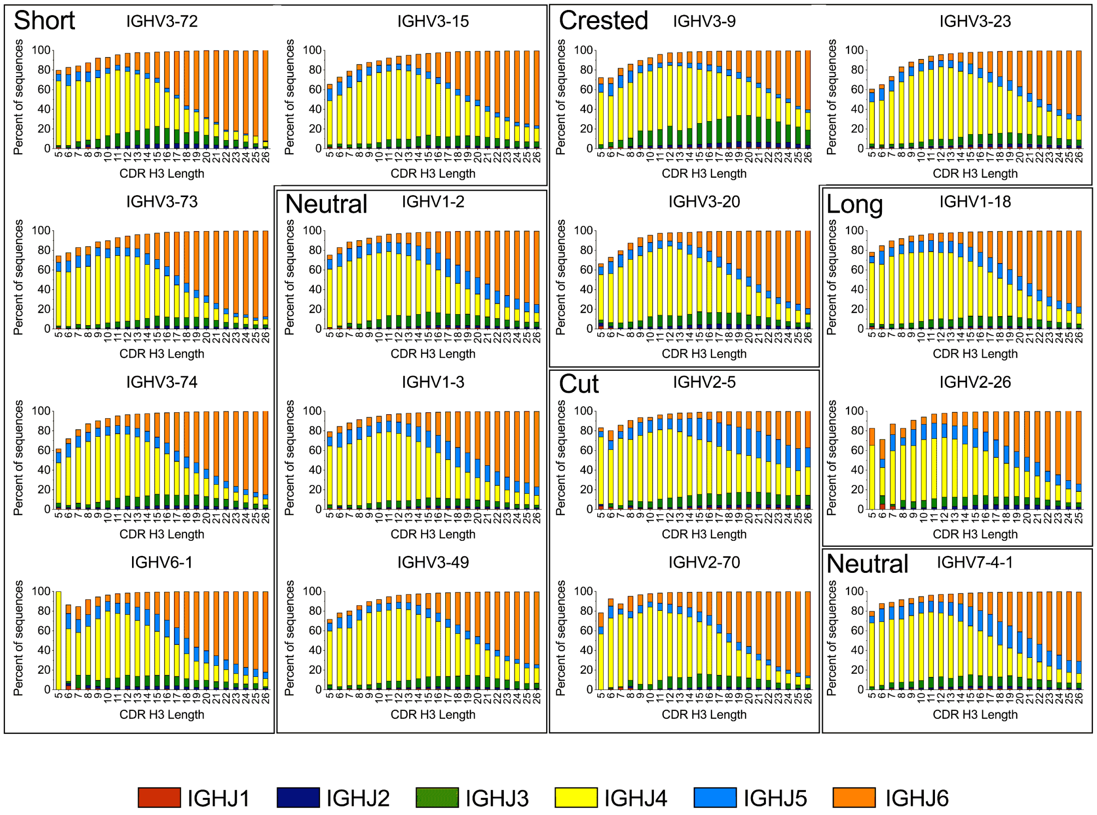

**Fig. S9.** $J_H$ germline distribution as a function of $V_H$ germline and CDR H3 length in unproductive sequences and in the naïve compartments of the WA dataset. Data is averaged for all donors. Germlines are noted in the IMGT naming convention. A fraction of the sequences does not have an unambiguously annotated $J_H$ segment, particularly in the short end of the spectrum.

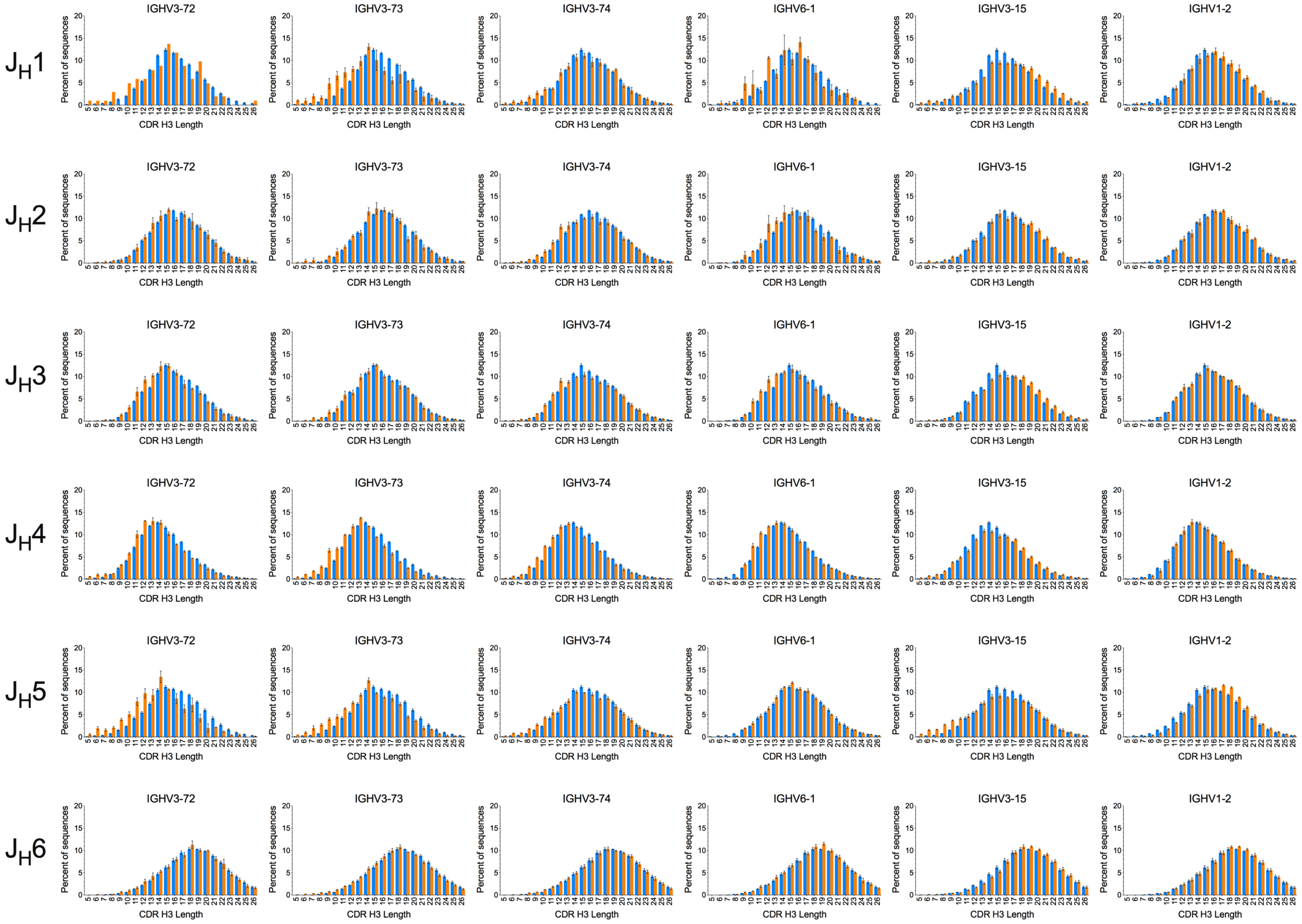

JH1
JH2
JH3
JH4
JH5
JH6
IGHV3-72
IGHV3-73
IGHV3-74
IGHV6-1
IGHV3-15
IGHV1-2
Percent of sequences
CDR H3 Length

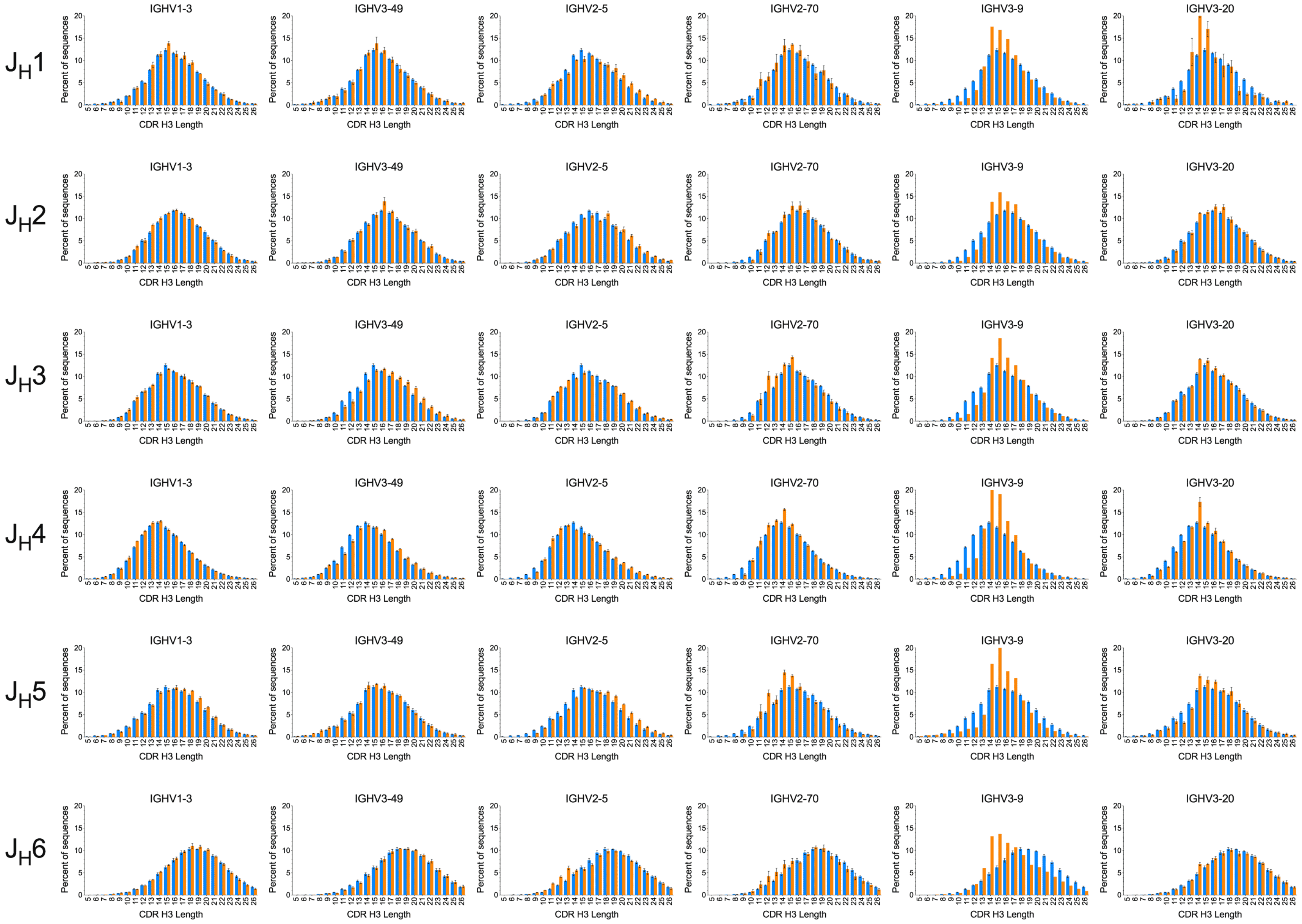

$J_H1$
$J_H2$
$J_H3$
$J_H4$
$J_H5$
$J_H6$
IGHV1-3
IGHV3-49
IGHV2-5
IGHV2-70
IGHV3-9
IGHV3-20
Percent of sequences
CDR H3 Length

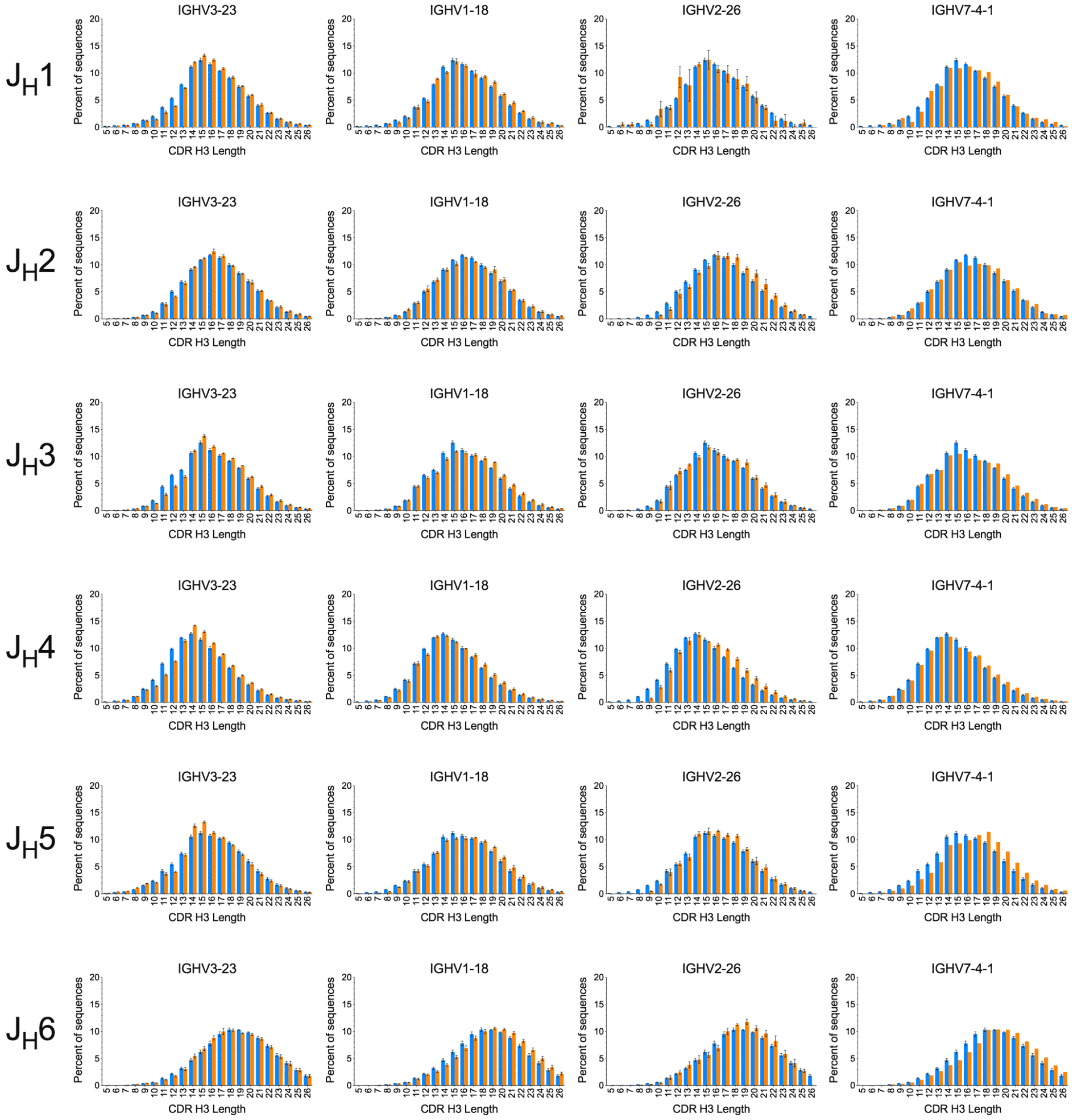


**Fig. S10.** CDR H3 distribution as a function of $V_H$ and $J_H$ germline in the WA naïve compartment. $V_H/J_H$ combination-specific distributions are shown in orange and JH-specific distributions in blue. $V_H$ germlines are noted in the IMGT naming convention. The number of CDR H3 amino acid residues potentially encoded by $J_H1$-$J_H6$ are 6, 7, 6, 4, 6 and 9 respectively.

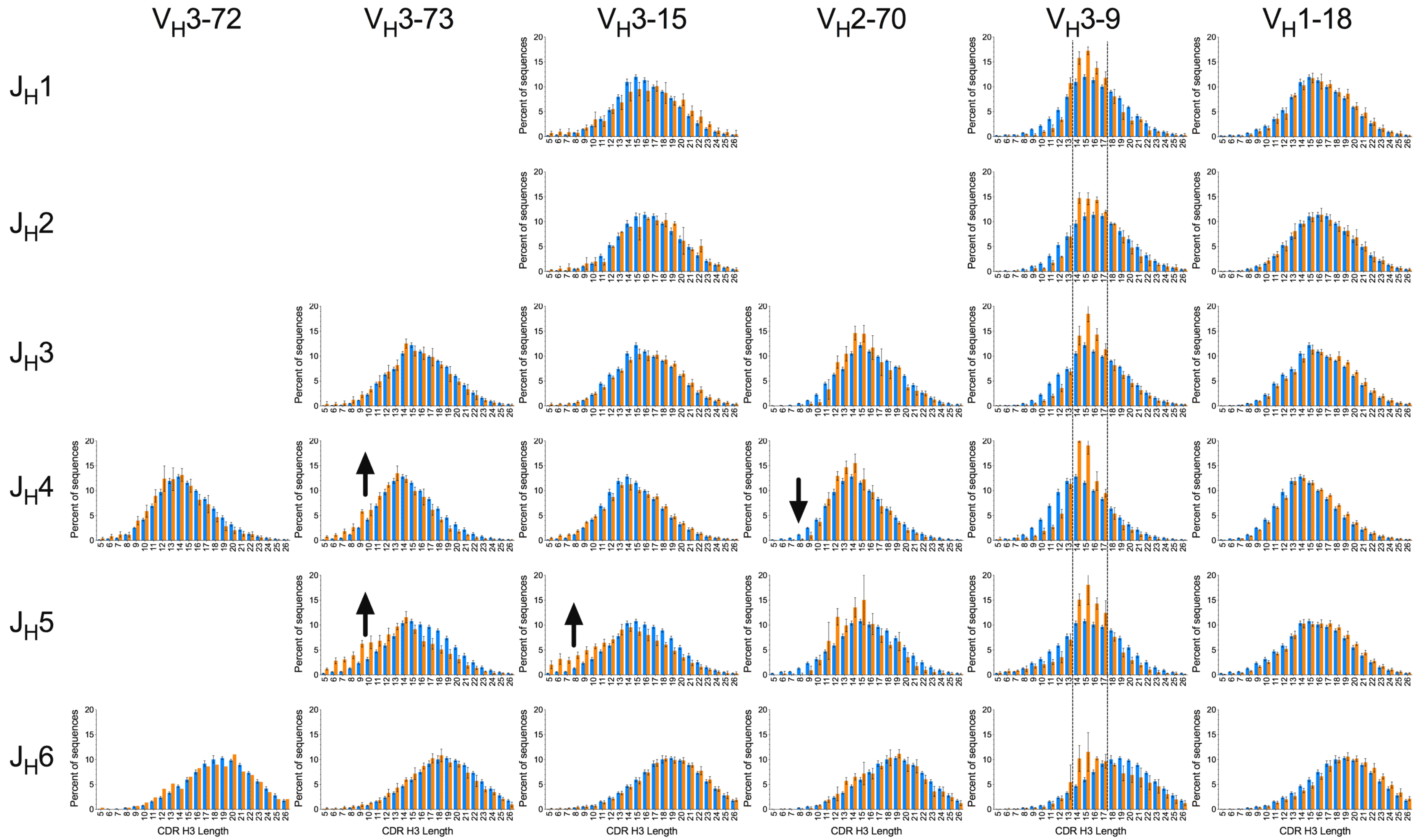


**Fig. S11.** Modulation of VH-associated CDR H3 length biases by JH segments in the SRI naïve (unmutated IgM, see Table S1) compartment. Colors, arrows and panel organization is the same as in Figure 5 for ease of comparison. Omitted panels had low sequence counts.

## IGHJ4: YFDY (IMGT 114-117)    WA, nonproductive sequences

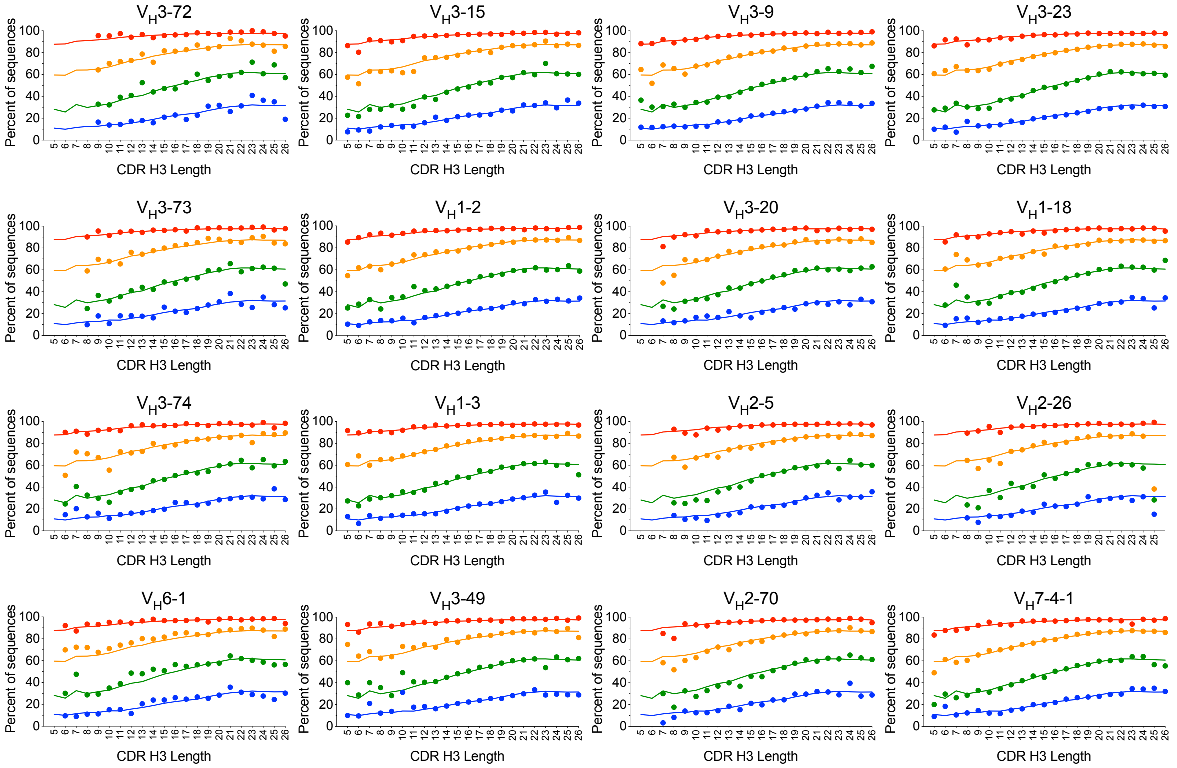


## IGHJ5: NWFDP (IMGT 113-117)    Nonproductive sequences

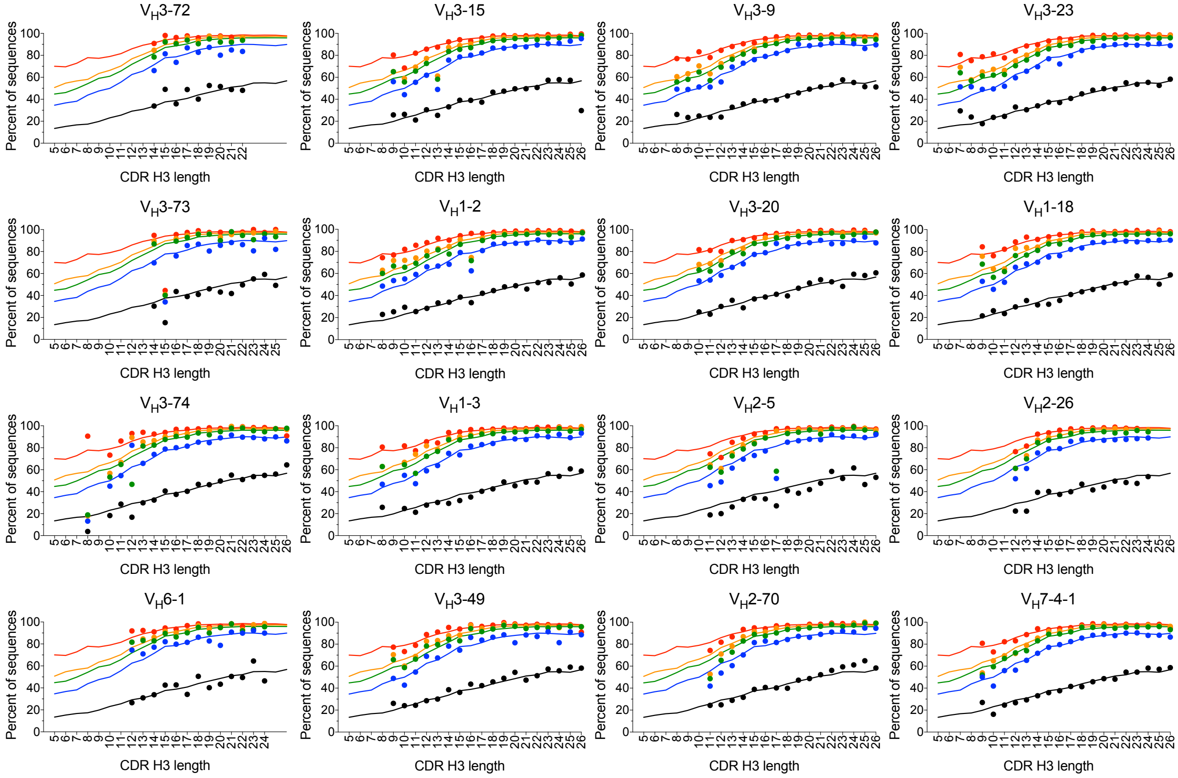

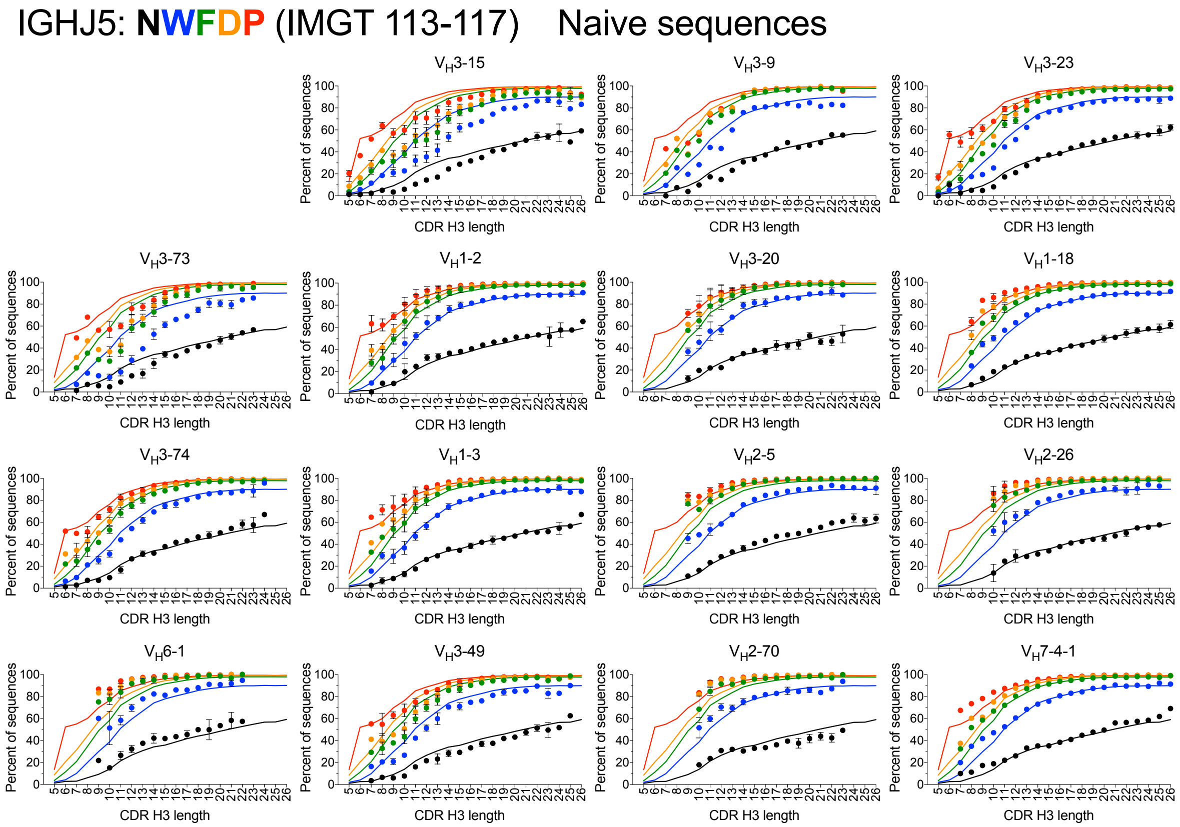


**Fig. S12.** Analysis of $J_H4$ and $J_H5$ sequence trimming as a function of $V_H$ germline use and CDR H3 length in WA unproductive and naïve compartment sequences. Occupancy of $J_H$ germline-encoded residues in the last CDR H3 positions are shown. $J_H$ residues are color-coded. Solid lines indicate residue occupancy for all sequences in the unproductive and naïve repertoire with a given $J_H$ segment. Dots indicate average residue occupancy with each $V_H/J_H$ combination CDR H3 length. Dots above and below the corresponding solid line indicate reduced and increased $J_H$ trimming relative to the overall repertoire. Note that trimming of $J_H$ segments in the non-productive sequences is not biased by $V_H$. Bars indicate S.E.M. for three donors except for $V_H3$-9 and $V_H7$-4-1, which are present in one donor each, and the nonproductive sequences, which were pooled before analysis. Data points with fewer than 60 sequences (30 for $V_H3$-9 and $V_H7$-4-1) were excluded. Panels are organized in the same order as in Fig. S11.

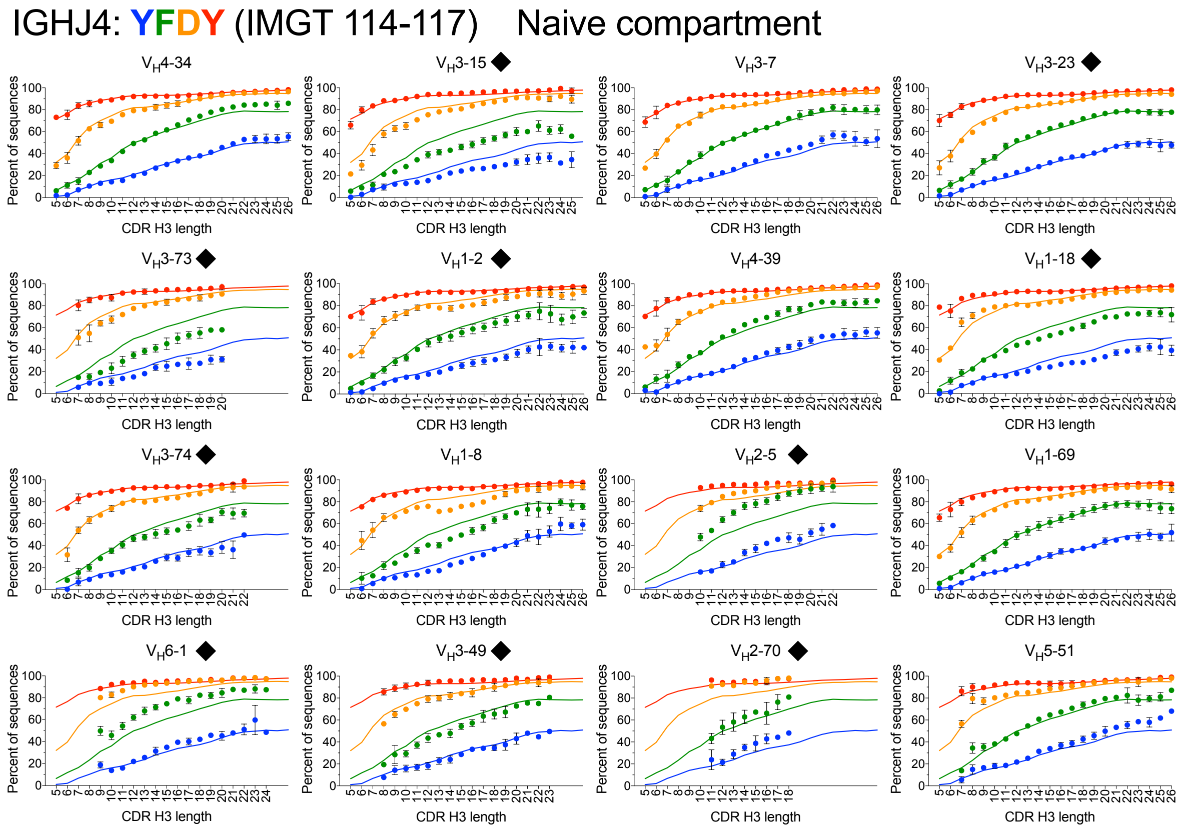


**Fig. S13.** Analysis of $J_H4$ sequence trimming as a function of $V_H$ germline use and CDR H3 length in SRI unmutated IgM (naïve) compartment sequences. Symbols shown as in Figure 6. The panels highlighted with black diamonds indicate germlines in the same place as in Figure 6. Other panels show different germlines according to data availability in the dataset. Data points with fewer than 60 sequences were excluded.

IGHJ3

| D germline length (nt) | Total | VH3-72 | VH3-73 | VH3-74 | VH6-1 | VH3-15 | VH1-2 | VH1-3 | VH3-49 | VH3-9 | VH3-20 | VH2-5 | VH2-70 | VH3-23 | VH1-18 | VH2-26 | VH7-4-1 |
|---|---|---|---|---|---|---|---|---|---|---|---|---|---|---|---|---|---|
| 37 | 1.7% | 2.1% | 1.9% | 1.5% | 2.7% | 1.5% | 2.2% | 1.7% | 1.4% | 1.8% | 1.8% | 1.5% | 1.9% | 1.4% | 1.8% | 1.6% | 1.8% |
| 31 | 24.7% | 27.4% | 25.2% | 23.0% | 23.4% | 25.9% | 27.9% | 23.5% | 24.1% | 26.8% | 23.9% | 20.0% | 22.1% | 23.9% | 26.2% | 18.3% | 23.3% |
| 28 | 0.5% | 0.6% | 0.4% | 0.5% | 0.4% | 0.7% | 0.7% | 0.6% | 0.6% | 0.1% | 0.5% | 0.5% | 0.4% | 0.5% | 0.6% | 0.5% | 1.5% |
| 23 | 8.1% | 8.3% | 7.1% | 8.1% | 6.6% | 8.6% | 7.4% | 11.0% | 8.1% | 9.6% | 8.4% | 9.1% | 8.1% | 8.3% | 8.3% | 9.5% | 8.0% |
| 21 | 15.1% | 15.3% | 12.4% | 15.6% | 12.7% | 15.3% | 12.5% | 14.4% | 15.8% | 12.2% | 13.3% | 13.9% | 14.9% | 17.4% | 14.8% | 16.7% | 5.7% |
| 20 | 19.1% | 19.5% | 23.1% | 20.5% | 11.3% | 24.2% | 16.7% | 17.8% | 23.6% | 13.4% | 21.3% | 24.6% | 16.5% | 19.2% | 20.0% | 22.0% | 26.7% |
| 19 | 1.8% | 1.6% | 2.1% | 1.7% | 1.4% | 1.3% | 1.6% | 1.1% | 1.4% | 1.9% | 1.5% | 1.4% | 2.2% | 1.5% | 1.5% | 2.0% | 0.5% |
| 18 | 3.0% | 2.7% | 2.5% | 2.7% | 3.2% | 2.6% | 2.9% | 3.0% | 3.0% | 2.7% | 3.4% | 2.9% | 3.0% | 2.5% | 2.6% | 3.7% | 4.7% |
| 17 | 9.7% | 7.7% | 8.2% | 8.9% | 9.5% | 6.9% | 9.4% | 9.3% | 8.6% | 13.8% | 9.0% | 9.6% | 11.1% | 9.4% | 8.3% | 9.9% | 6.8% |
| 16 | 9.8% | 7.7% | 10.0% | 9.0% | 16.6% | 7.2% | 10.4% | 11.0% | 8.4% | 11.7% | 10.8% | 8.5% | 11.6% | 10.3% | 9.8% | 9.3% | 13.8% |
| 11 | 1.3% | 1.2% | 1.3% | 1.5% | 4.0% | 1.0% | 1.9% | 1.1% | 0.9% | 1.6% | 1.0% | 1.7% | 1.6% | 1.1% | 1.2% | 1.4% | 1.1% |
| Weighted average (nt) | 21.5 | 21.9 | 21.5 | 21.0 | 20.4 | 22.0 | 21.6 | 21.4 | 21.8 | 21.7 | 21.5 | 20.8 | 20.8 | 21.6 | 21.8 | 20.9 | 21.1 |

IGHJ4

| D germline length (nt) | Total | VH3-72 | VH3-73 | VH3-74 | VH6-1 | VH3-15 | VH1-2 | VH1-3 | VH3-49 | VH3-9 | VH3-20 | VH2-5 | VH2-70 | VH3-23 | VH1-18 | VH2-26 | VH7-4-1 |
|---|---|---|---|---|---|---|---|---|---|---|---|---|---|---|---|---|---|
| 37 | 4.4% | 4.6% | 4.1% | 4.5% | 4.1% | 4.7% | 4.5% | 4.6% | 4.2% | 3.0% | 4.7% | 4.8% | 4.1% | 4.2% | 4.4% | 4.5% | 5.5% |
| 31 | 40.3% | 37.6% | 34.6% | 38.2% | 24.4% | 47.4% | 38.0% | 39.8% | 45.0% | 38.6% | 40.4% | 42.5% | 33.8% | 43.0% | 41.8% | 43.0% | 40.4% |
| 28 | 2.1% | 2.5% | 2.5% | 2.3% | 1.6% | 2.2% | 1.8% | 1.8% | 2.4% | 1.2% | 1.7% | 1.7% | 1.9% | 2.3% | 2.0% | 2.0% | 1.4% |
| 23 | 4.0% | 4.2% | 3.7% | 4.0% | 4.1% | 3.6% | 4.2% | 4.1% | 3.8% | 3.4% | 3.9% | 3.9% | 3.9% | 3.3% | 3.7% | 4.6% | 5.1% |
| 21 | 14.8% | 10.4% | 14.7% | 13.7% | 26.9% | 10.2% | 15.5% | 18.1% | 12.8% | 20.8% | 17.1% | 14.1% | 18.1% | 15.3% | 15.2% | 14.8% | 16.5% |
| 20 | 12.8% | 16.8% | 13.7% | 14.2% | 12.8% | 11.5% | 11.8% | 12.1% | 12.7% | 15.0% | 10.9% | 10.0% | 14.9% | 12.6% | 12.6% | 13.3% | 8.4% |
| 19 | 1.6% | 1.8% | 2.3% | 1.7% | 1.5% | 1.1% | 1.4% | 1.1% | 1.3% | 1.7% | 1.2% | 1.2% | 1.9% | 1.4% | 1.4% | 1.5% | 1.0% |
| 18 | 3.7% | 2.3% | 3.8% | 3.5% | 5.1% | 2.2% | 5.1% | 3.2% | 3.0% | 4.4% | 2.8% | 4.0% | 4.3% | 3.3% | 3.5% | 2.6% | 3.7% |
| 17 | 3.3% | 4.1% | 3.9% | 3.2% | 5.5% | 3.6% | 4.3% | 3.3% | 3.0% | 3.1% | 2.8% | 3.7% | 3.6% | 2.7% | 3.4% | 2.8% | 3.0% |
| 16 | 8.0% | 9.7% | 10.8% | 8.0% | 5.5% | 8.4% | 8.0% | 6.9% | 7.6% | 4.8% | 10.4% | 8.7% | 8.4% | 7.1% | 7.7% | 7.4% | 7.6% |
| 11 | 0.9% | 1.0% | 1.0% | 1.1% | 2.9% | 0.7% | 1.3% | 0.8% | 0.7% | 1.0% | 0.7% | 1.1% | 1.2% | 0.8% | 0.8% | 0.9% | 2.4% |
| Weighted average (nt) | 24.2 | 23.7 | 23.2 | 23.7 | 22.0 | 25.0 | 23.9 | 24.2 | 24.9 | 24.1 | 24.4 | 24.4 | 23.4 | 24.6 | 24.5 | 24.9 | 24.1 |

IGHJ5

| D germline length (nt) | Total | VH3-72 | VH3-73 | VH3-74 | VH6-1 | VH3-15 | VH1-2 | VH1-3 | VH3-49 | VH3-9 | VH3-20 | VH2-5 | VH2-70 | VH3-23 | VH1-18 | VH2-26 | VH7-4-1 |
|---|---|---|---|---|---|---|---|---|---|---|---|---|---|---|---|---|---|
| 37 | 4.2% | 5.6% | 4.1% | 4.3% | 4.1% | 4.3% | 4.2% | 4.2% | 3.6% | 3.3% | 4.8% | 4.2% | 4.8% | 3.9% | 4.1% | 4.1% | 3.9% |
| 31 | 47.9% | 38.8% | 42.5% | 47.2% | 32.1% | 53.7% | 47.4% | 49.3% | 50.7% | 42.4% | 43.7% | 52.5% | 41.0% | 47.6% | 49.1% | 46.6% | 49.8% |
| 28 | 1.9% | 3.2% | 2.2% | 1.9% | 2.1% | 2.2% | 1.4% | 1.7% | 2.1% | 1.1% | 1.5% | 1.5% | 2.1% | 1.9% | 1.6% | 1.8% | 1.2% |
| 23 | 2.4% | 3.3% | 2.7% | 2.5% | 3.0% | 2.4% | 2.5% | 2.6% | 2.4% | 2.2% | 2.2% | 2.1% | 2.6% | 2.2% | 2.3% | 2.5% | 3.2% |
| 21 | 13.5% | 10.2% | 14.2% | 13.7% | 24.2% | 9.1% | 13.6% | 15.7% | 12.6% | 20.0% | 16.3% | 12.6% | 15.4% | 14.9% | 13.7% | 16.1% | 16.0% |
| 20 | 7.9% | 11.6% | 9.3% | 8.9% | 8.9% | 7.6% | 7.4% | 7.2% | 8.3% | 10.8% | 7.6% | 5.9% | 8.8% | 8.3% | 8.0% | 8.2% | 4.8% |
| 19 | 1.0% | 1.1% | 1.2% | 1.0% | 1.0% | 0.8% | 0.9% | 0.8% | 0.9% | 1.0% | 0.9% | 0.7% | 1.2% | 0.9% | 0.9% | 1.2% | 0.8% |
| 18 | 4.1% | 2.0% | 3.6% | 3.4% | 5.1% | 2.2% | 5.5% | 3.4% | 3.3% | 4.3% | 3.1% | 4.0% | 4.7% | 3.7% | 3.8% | 3.4% | 4.2% |
| 17 | 4.0% | 5.5% | 3.9% | 3.3% | 7.2% | 3.8% | 5.1% | 3.9% | 3.7% | 4.6% | 3.9% | 3.9% | 4.7% | 3.6% | 4.2% | 3.8% | 4.7% |
| 16 | 7.5% | 10.5% | 9.3% | 6.7% | 5.3% | 6.8% | 7.0% | 6.4% | 7.1% | 5.0% | 11.0% | 7.2% | 8.5% | 7.5% | 7.0% | 7.2% | 6.2% |
| 11 | 0.7% | 0.7% | 0.7% | 0.7% | 1.6% | 0.6% | 0.8% | 0.5% | 0.6% | 0.8% | 0.8% | 0.7% | 0.9% | 0.7% | 0.6% | 0.7% | 0.8% |
| Weighted average (nt) | 24.8 | 23.5 | 23.9 | 24.5 | 23.0 | 25.2 | 24.8 | 25.2 | 25.1 | 24.2 | 24.4 | 25.3 | 24.0 | 24.8 | 24.9 | 24.8 | 25.1 |

IGHJ6

| D germline length (nt) | Total | VH3-72 | VH3-73 | VH3-74 | VH6-1 | VH3-15 | VH1-2 | VH1-3 | VH3-49 | VH3-9 | VH3-20 | VH2-5 | VH2-70 | VH3-23 | VH1-18 | VH2-26 | VH7-4-1 |
|---|---|---|---|---|---|---|---|---|---|---|---|---|---|---|---|---|---|
| 37 | 4.1% | 4.3% | 3.8% | 3.8% | 3.8% | 3.8% | 4.2% | 4.0% | 3.8% | 4.6% | 5.1% | 4.5% | 4.1% | 4.0% | 3.7% | 3.9% | 4.1% |
| 31 | 46.6% | 45.1% | 43.3% | 47.1% | 32.0% | 50.0% | 45.4% | 47.8% | 49.5% | 40.8% | 47.1% | 45.9% | 38.6% | 47.6% | 50.3% | 45.8% | 51.1% |
| 28 | 1.8% | 2.0% | 2.0% | 1.9% | 1.7% | 1.8% | 1.5% | 1.6% | 1.8% | 1.4% | 1.5% | 1.3% | 1.6% | 1.8% | 1.7% | 1.6% | 1.0% |
| 23 | 3.3% | 3.2% | 3.1% | 2.9% | 4.3% | 3.2% | 3.3% | 3.4% | 3.1% | 2.7% | 2.9% | 2.9% | 2.9% | 2.8% | 3.0% | 3.9% | 3.7% |
| 21 | 10.6% | 7.9% | 10.1% | 10.7% | 20.1% | 8.9% | 11.3% | 12.0% | 10.1% | 13.0% | 10.8% | 9.9% | 12.5% | 11.3% | 11.0% | 12.5% | 13.0% |
| 20 | 9.0% | 10.8% | 9.9% | 9.6% | 9.7% | 8.7% | 8.6% | 8.4% | 9.2% | 9.9% | 8.0% | 7.2% | 10.2% | 8.9% | 8.3% | 9.2% | 6.6% |
| 19 | 1.1% | 1.4% | 1.3% | 1.1% | 1.1% | 0.8% | 1.0% | 0.9% | 1.0% | 1.3% | 0.8% | 0.7% | 1.3% | 1.0% | 0.9% | 1.2% | 0.7% |
| 18 | 4.6% | 2.9% | 4.9% | 3.9% | 6.1% | 3.1% | 5.7% | 3.9% | 3.8% | 5.3% | 3.5% | 5.1% | 6.3% | 4.0% | 3.8% | 3.5% | 3.4% |
| 17 | 3.7% | 4.4% | 3.8% | 3.2% | 7.1% | 4.0% | 4.7% | 3.9% | 3.3% | 4.0% | 3.6% | 3.7% | 4.0% | 3.3% | 3.8% | 3.4% | 3.1% |
| 16 | 7.8% | 8.9% | 10.0% | 7.3% | 4.8% | 8.7% | 7.4% | 7.2% | 7.9% | 4.9% | 8.8% | 8.1% | 8.6% | 7.9% | 7.3% | 7.0% | 5.8% |
| 11 | 0.7% | 0.7% | 0.8% | 0.7% | 2.0% | 0.6% | 0.8% | 0.7% | 0.7% | 1.0% | 0.8% | 0.9% | 0.9% | 0.8% | 0.6% | 0.7% | 0.7% |
| Weighted average (nt) | 24.2 | 23.7 | 23.7 | 24.1 | 22.5 | 24.6 | 24.2 | 24.5 | 24.7 | 22.8 | 24.3 | 23.5 | 22.8 | 24.4 | 24.8 | 24.1 | 24.8 |

**Fig. S14.** D germline prevalence associated with different $V_H$ and $J_H$ germlines in the WA naïve compartment. D germlines are grouped by length. The D germline prevalence deviations from averages associated with $V_H$6-1 are boxed.

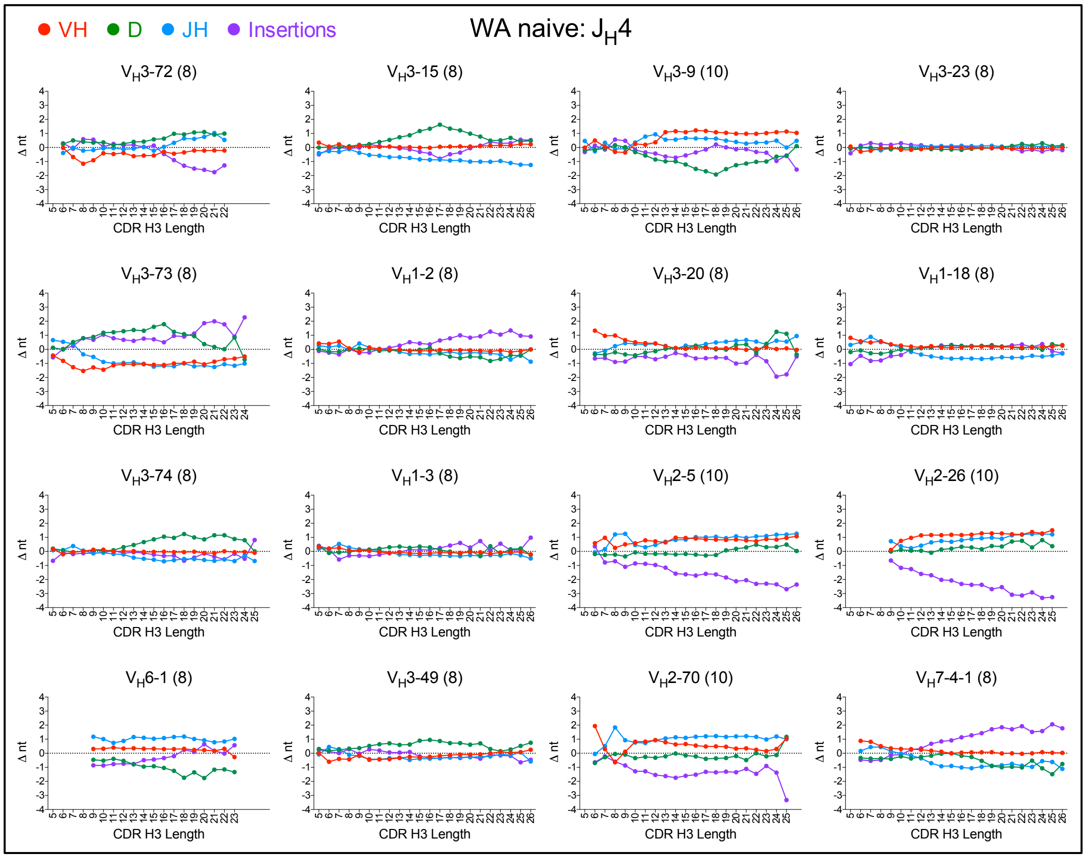
VH D JH Insertions
WA naive: $J_H4$
$V_H$3-72 (8)
$V_H$3-15 (8)
$V_H$3-9 (10)
$V_H$3-23 (8)
$V_H$3-73 (8)
$V_H$1-2 (8)
$V_H$3-20 (8)
$V_H$1-18 (8)
$V_H$3-74 (8)
$V_H$1-3 (8)
$V_H$2-5 (10)
$V_H$2-26 (10)
$V_H$6-1 (8)
$V_H$3-49 (8)
$V_H$2-70 (10)
$V_H$7-4-1 (8)
Δ nt
CDR H3 Length

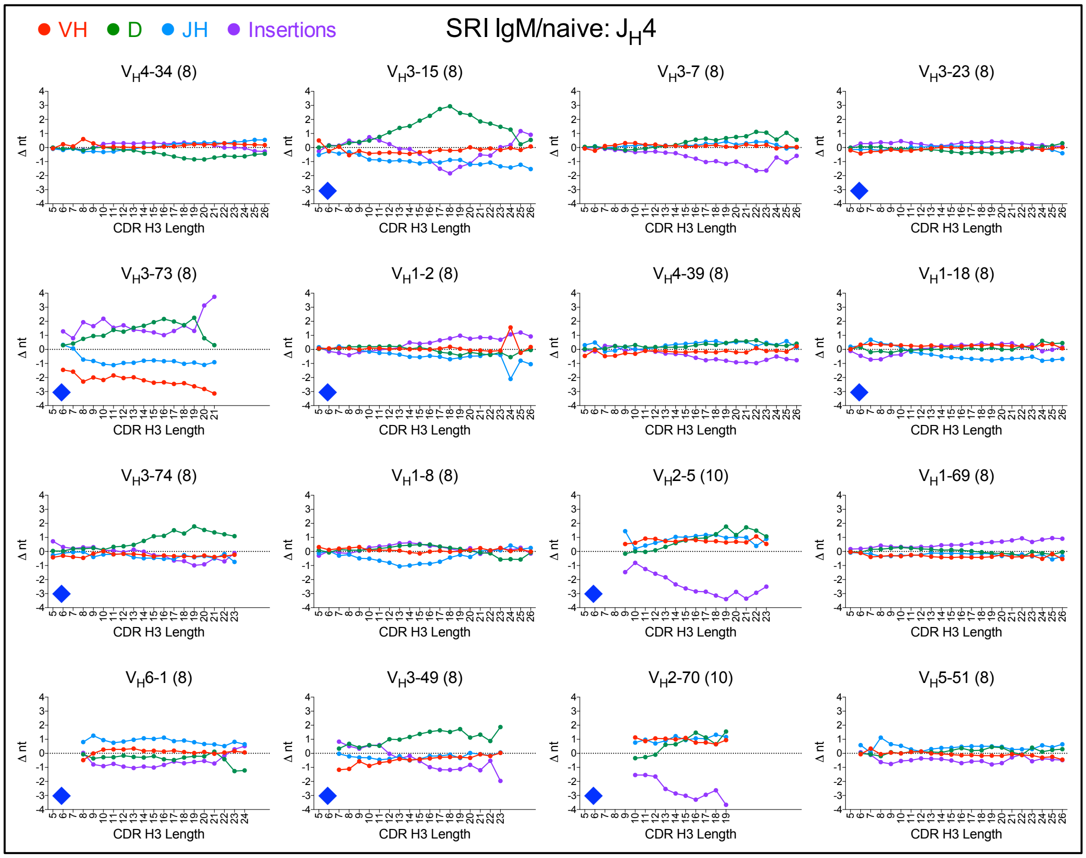
VH D JH Insertions
SRI IgM/naive: $J_H4$
$V_H$4-34 (8)
$V_H$3-15 (8)
$V_H$3-7 (8)
$V_H$3-23 (8)
$V_H$3-73 (8)
$V_H$1-2 (8)
$V_H$4-39 (8)
$V_H$1-18 (8)
$V_H$3-74 (8)
$V_H$1-8 (8)
$V_H$2-5 (10)
$V_H$1-69 (8)
$V_H$6-1 (8)
$V_H$3-49 (8)
$V_H$2-70 (10)
$V_H$5-51 (8)
Δ nt
CDR H3 Length

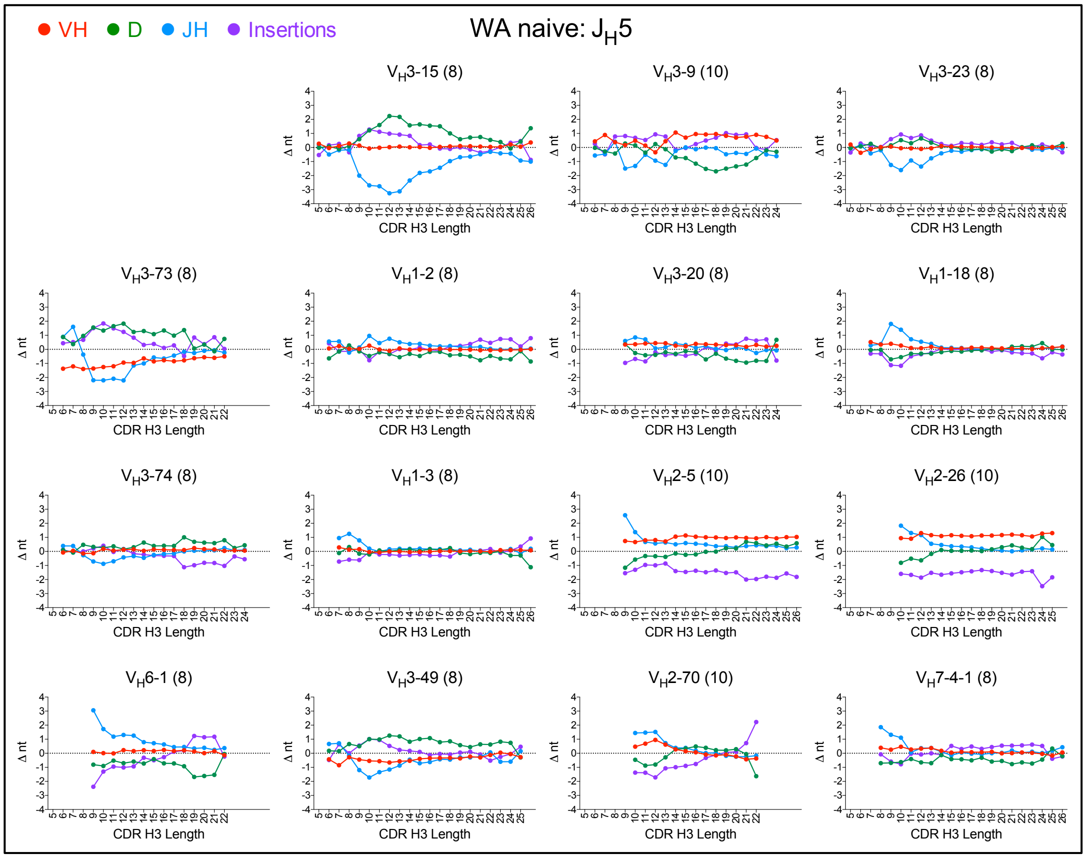
VH
D
JH
Insertions
WA naive: $J_H5$
$V_H3$-15 (8)
$V_H3$-9 (10)
$V_H3$-23 (8)
$V_H3$-73 (8)
$V_H1$-2 (8)
$V_H3$-20 (8)
$V_H1$-18 (8)
$V_H3$-74 (8)
$V_H1$-3 (8)
$V_H2$-5 (10)
$V_H2$-26 (10)
$V_H6$-1 (8)
$V_H3$-49 (8)
$V_H2$-70 (10)
$V_H7$-4-1 (8)
Δ nt
CDR H3 Length

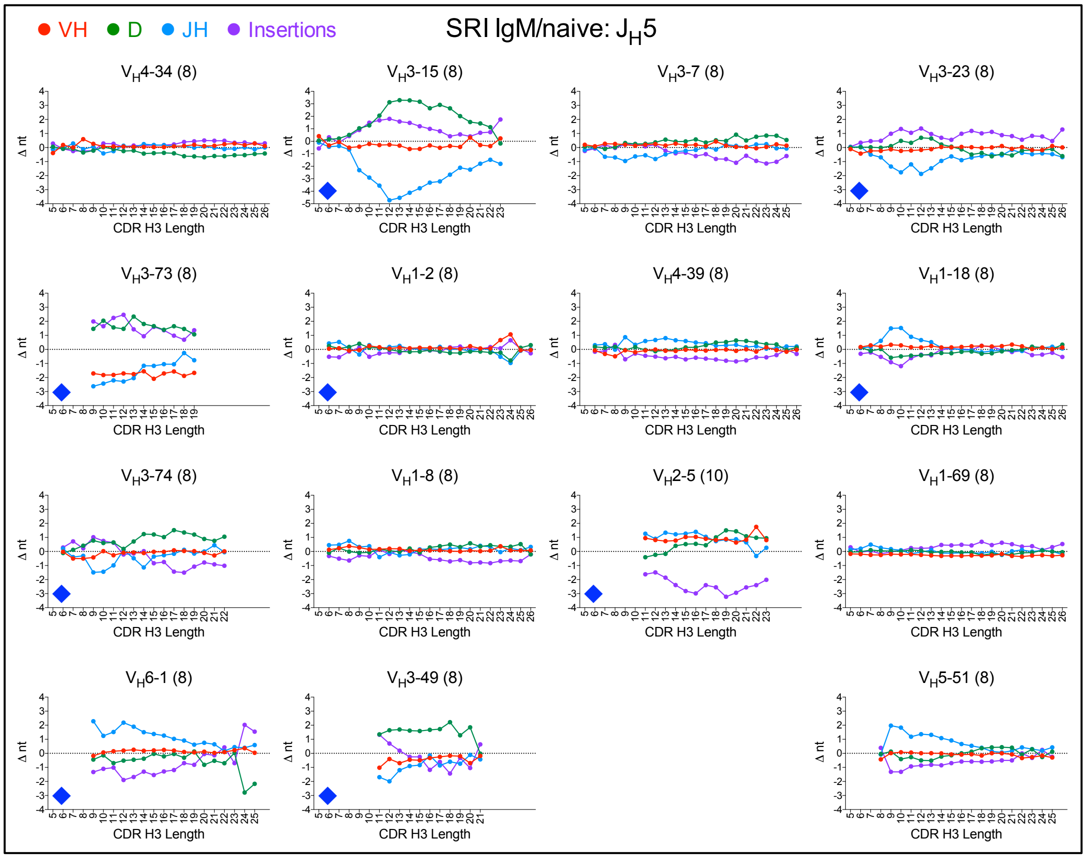
VH
D
JH
Insertions
SRI IgM/naive: $J_H5$
$V_H4$-34 (8)
$V_H3$-15 (8)
$V_H3$-7 (8)
$V_H3$-23 (8)
$V_H3$-73 (8)
$V_H1$-2 (8)
$V_H4$-39 (8)
$V_H1$-18 (8)
$V_H3$-74 (8)
$V_H1$-8 (8)
$V_H2$-5 (10)
$V_H1$-69 (8)
$V_H6$-1 (8)
$V_H3$-49 (8)
$V_H5$-51 (8)
Δ nt
CDR H3 Length

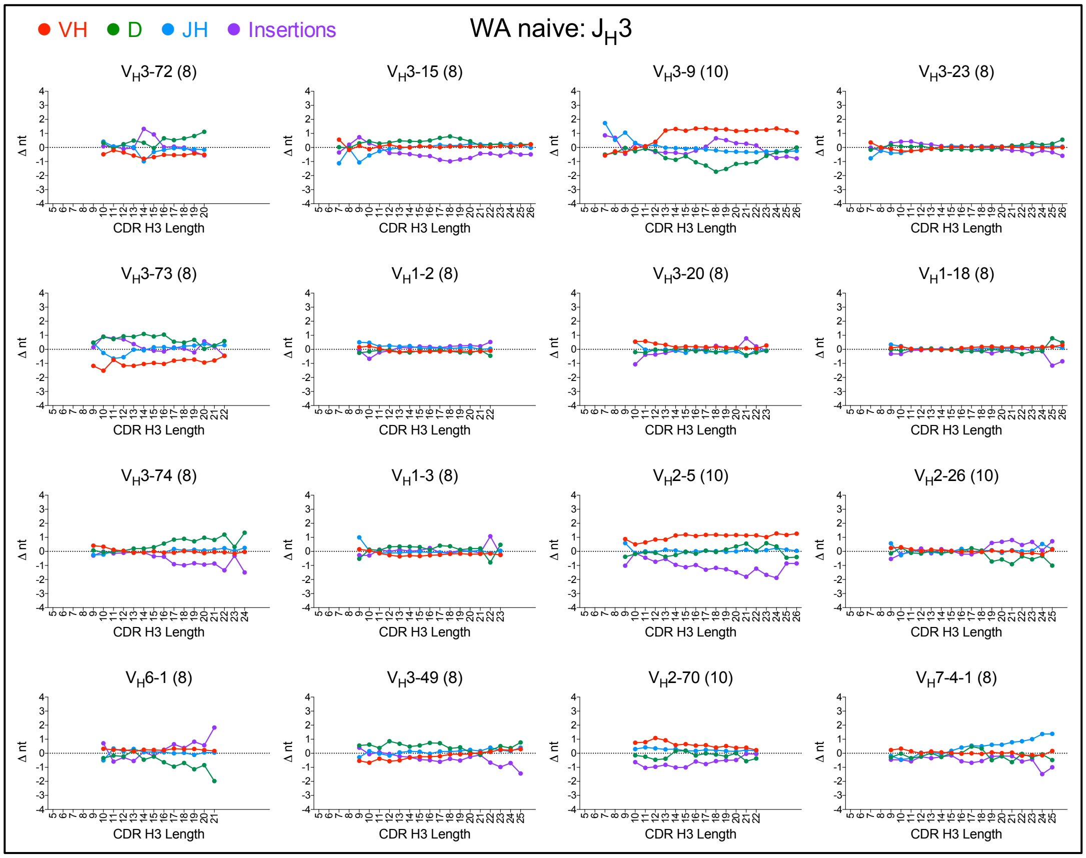
VH D JH Insertions
WA naive: $J_H3$
$V_H3$-72 (8)
$V_H3$-15 (8)
$V_H3$-9 (10)
$V_H3$-23 (8)
$V_H3$-73 (8)
$V_H1$-2 (8)
$V_H3$-20 (8)
$V_H1$-18 (8)
$V_H3$-74 (8)
$V_H1$-3 (8)
$V_H2$-5 (10)
$V_H2$-26 (10)
$V_H6$-1 (8)
$V_H3$-49 (8)
$V_H2$-70 (10)
$V_H7$-4-1 (8)
Δ nt
CDR H3 Length

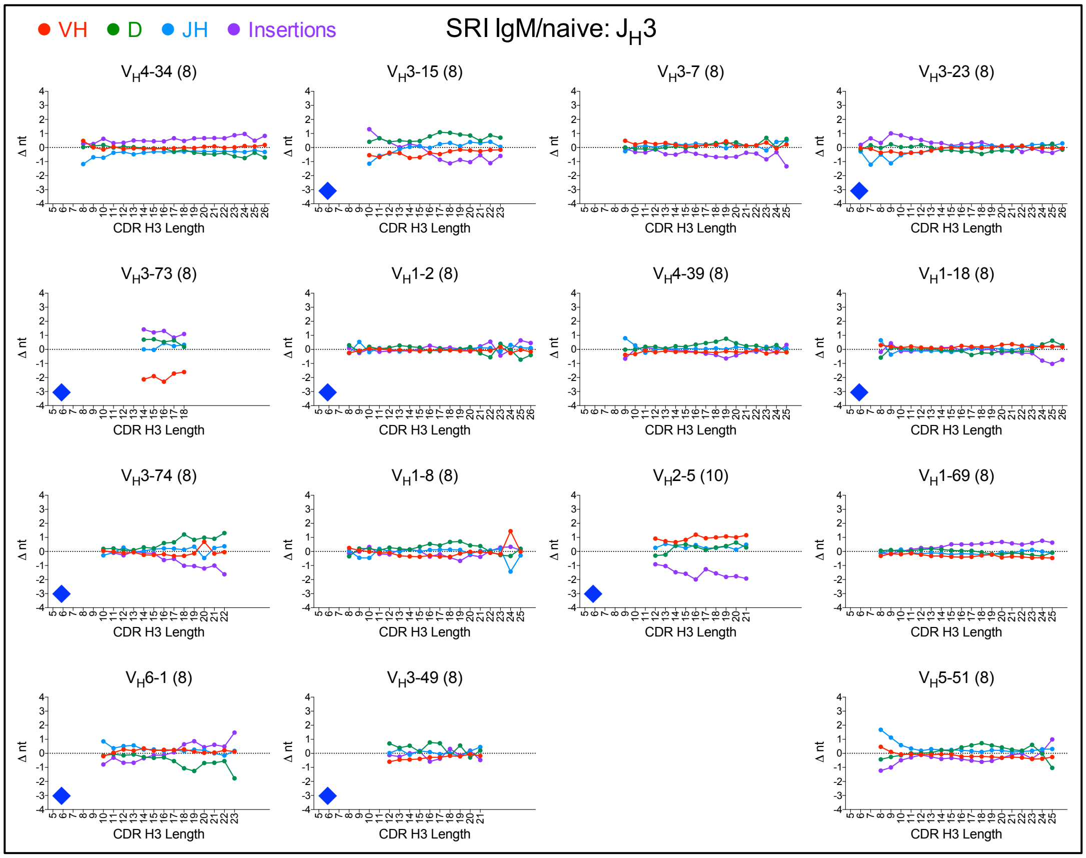
VH D JH Insertions
SRI IgM/naive: $J_H3$
$V_H4$-34 (8)
$V_H3$-15 (8)
$V_H3$-7 (8)
$V_H3$-23 (8)
$V_H3$-73 (8)
$V_H1$-2 (8)
$V_H4$-39 (8)
$V_H1$-18 (8)
$V_H3$-74 (8)
$V_H1$-8 (8)
$V_H2$-5 (10)
$V_H1$-69 (8)
$V_H6$-1 (8)
$V_H3$-49 (8)
$V_H5$-51 (8)
Δ nt
CDR H3 Length

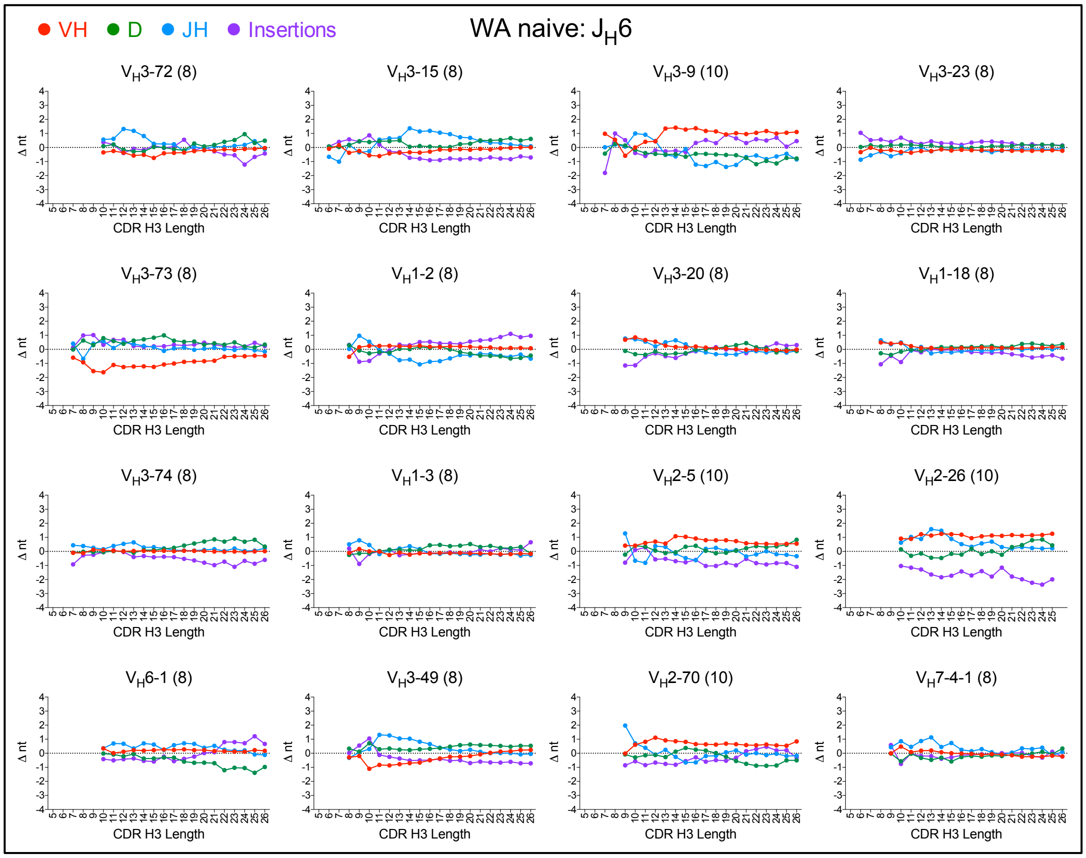
VH D JH Insertions
WA naive: $J_H6$
$V_H3$-72 (8)
$V_H3$-15 (8)
$V_H3$-9 (10)
$V_H3$-23 (8)
$V_H3$-73 (8)
$V_H1$-2 (8)
$V_H3$-20 (8)
$V_H1$-18 (8)
$V_H3$-74 (8)
$V_H1$-3 (8)
$V_H2$-5 (10)
$V_H2$-26 (10)
$V_H6$-1 (8)
$V_H3$-49 (8)
$V_H2$-70 (10)
$V_H7$-4-1 (8)
Δ nt
CDR H3 Length

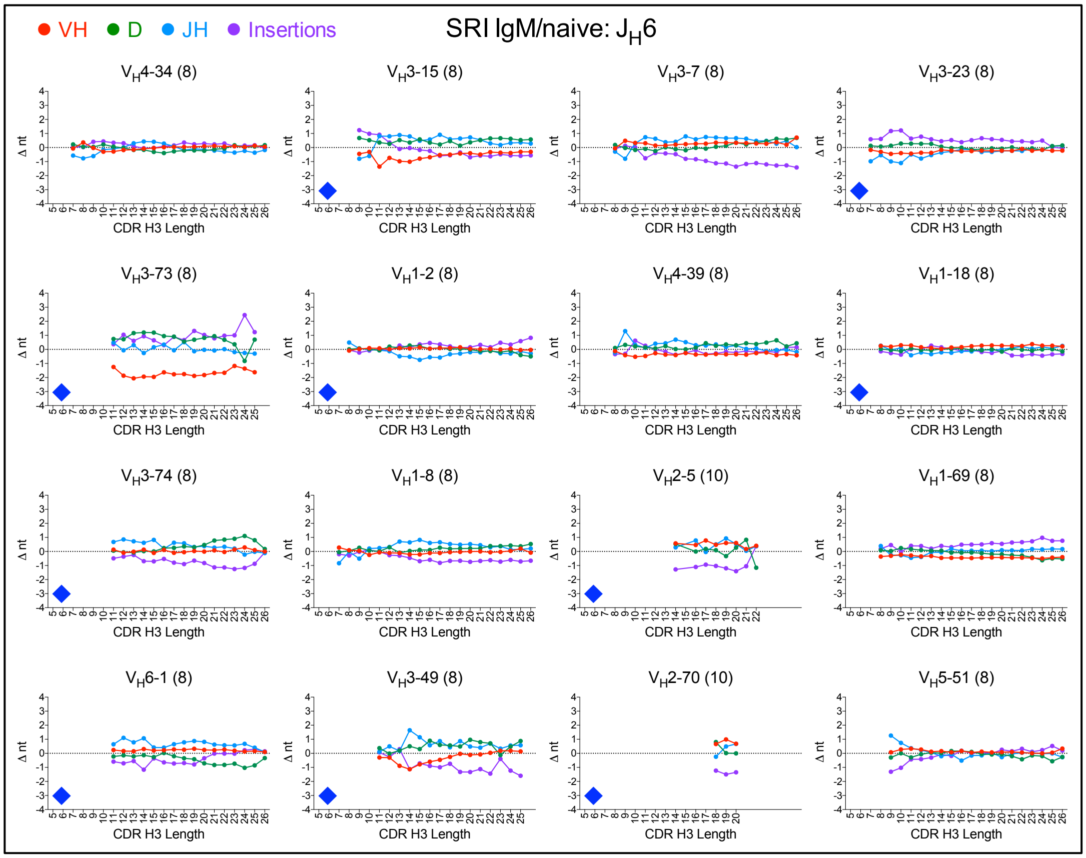
VH D JH Insertions
SRI IgM/naive: $J_H6$
$V_H4$-34 (8)
$V_H3$-15 (8)
$V_H3$-7 (8)
$V_H3$-23 (8)
$V_H3$-73 (8)
$V_H1$-2 (8)
$V_H4$-39 (8)
$V_H1$-18 (8)
$V_H3$-74 (8)
$V_H1$-8 (8)
$V_H2$-5 (10)
$V_H1$-69 (8)
$V_H6$-1 (8)
$V_H3$-49 (8)
$V_H2$-70 (10)
$V_H5$-51 (8)
Δ nt
CDR H3 Length

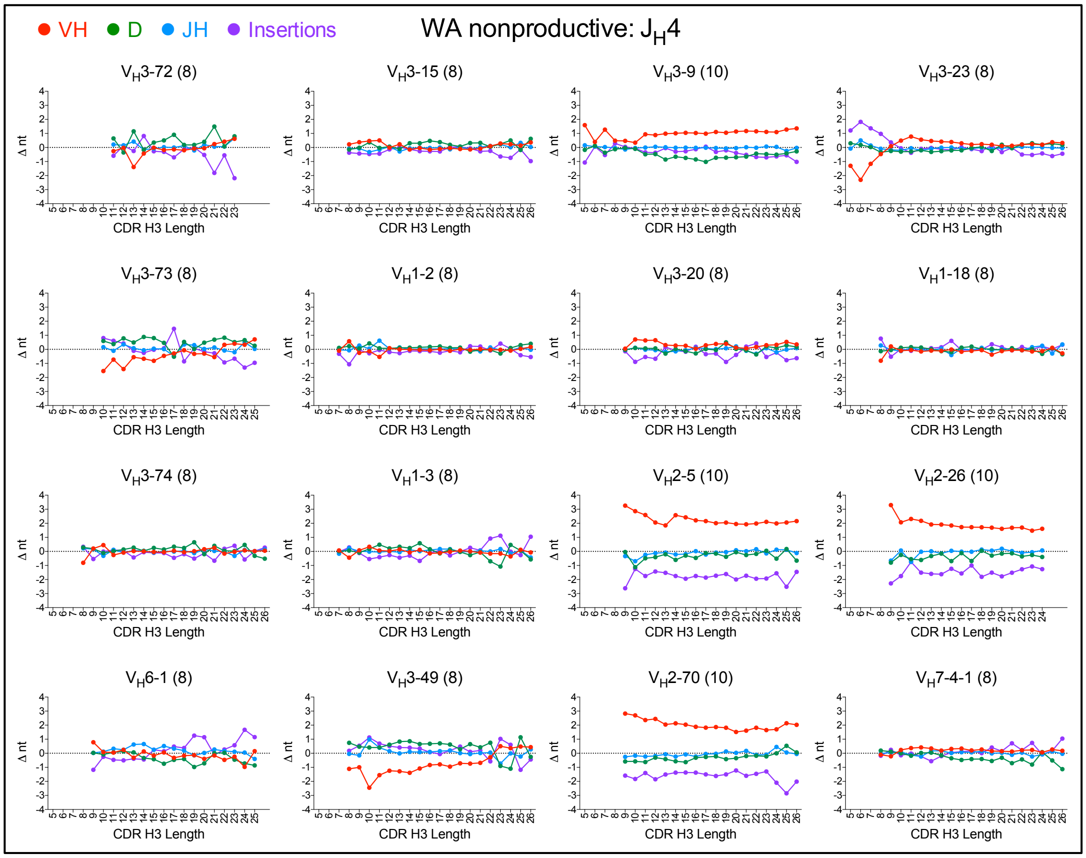
VH
D
JH
Insertions
WA nonproductive: JH4
VH3-72 (8)
VH3-15 (8)
VH3-9 (10)
VH3-23 (8)
VH3-73 (8)
VH1-2 (8)
VH3-20 (8)
VH1-18 (8)
VH3-74 (8)
VH1-3 (8)
VH2-5 (10)
VH2-26 (10)
VH6-1 (8)
VH3-49 (8)
VH2-70 (10)
VH7-4-1 (8)
Δ nt
CDR H3 Length

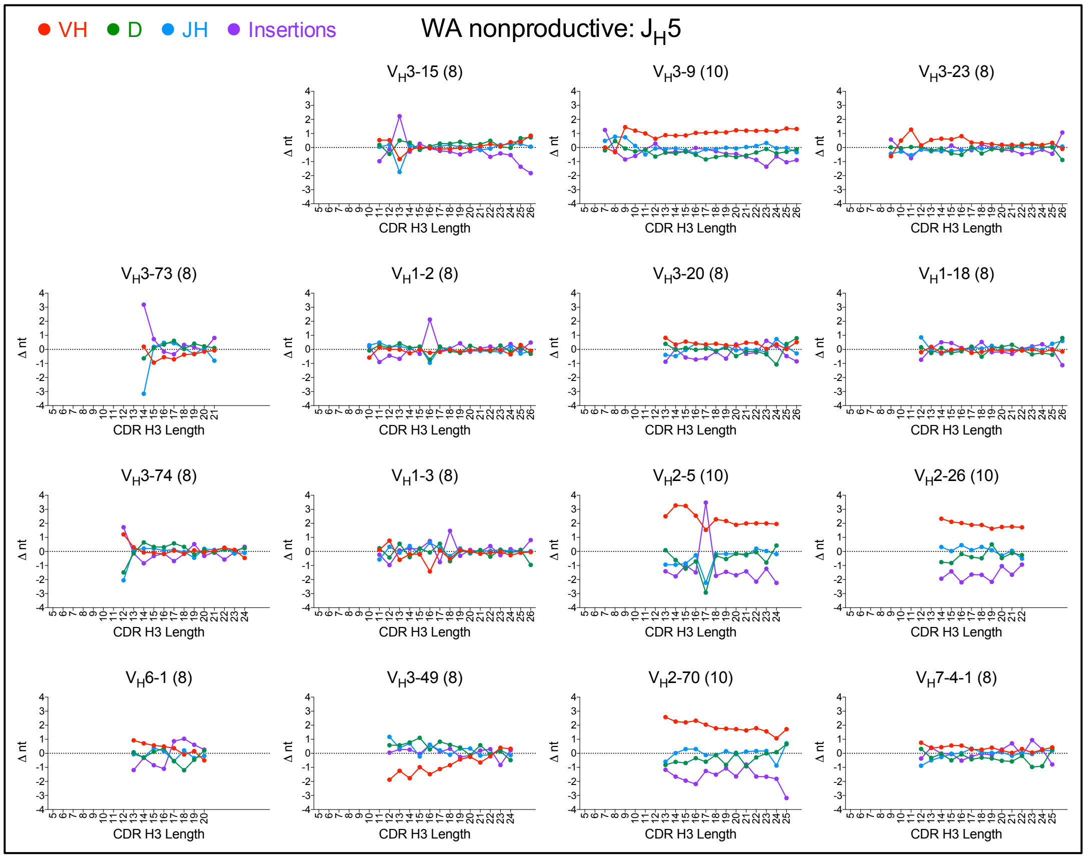
VH
D
JH
Insertions
WA nonproductive: JH5
VH3-15 (8)
VH3-9 (10)
VH3-23 (8)
VH3-73 (8)
VH1-2 (8)
VH3-20 (8)
VH1-18 (8)
VH3-74 (8)
VH1-3 (8)
VH2-5 (10)
VH2-26 (10)
VH6-1 (8)
VH3-49 (8)
VH2-70 (10)
VH7-4-1 (8)
Δ nt
CDR H3 Length

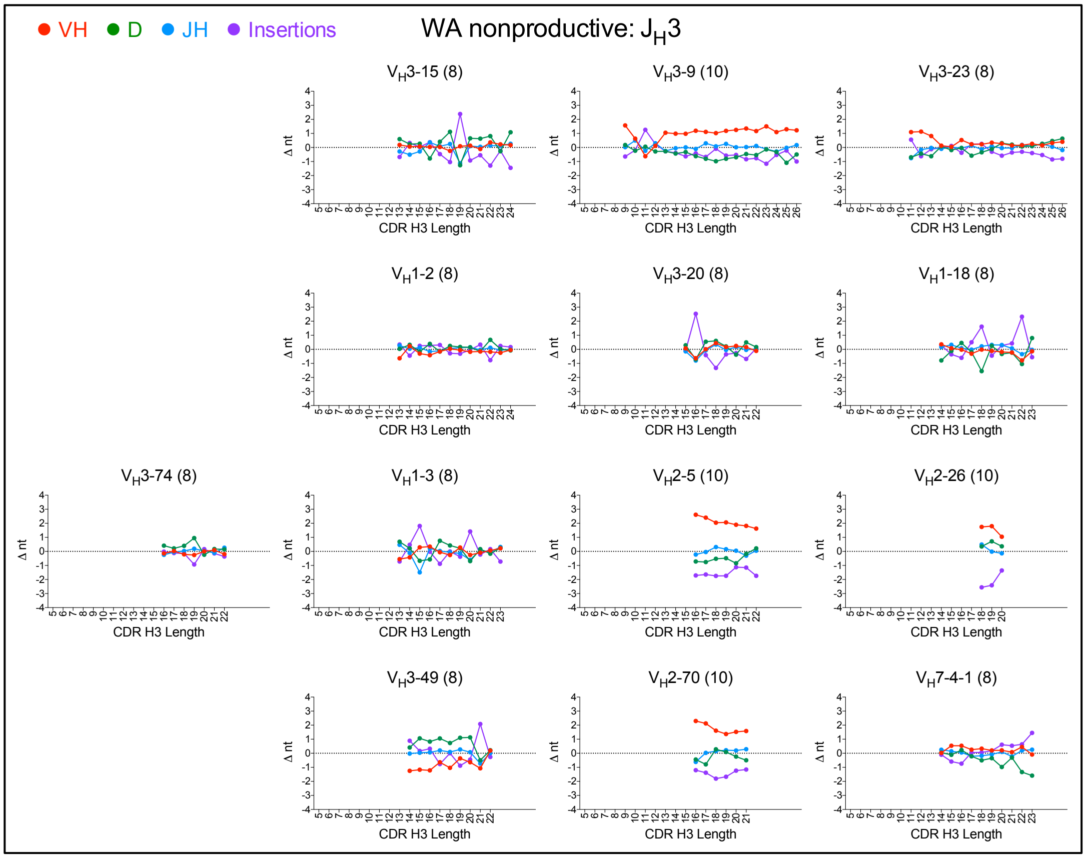
VH
D
JH
Insertions
WA nonproductive: JH3
VH3-15 (8)
VH3-9 (10)
VH3-23 (8)
VH1-2 (8)
VH3-20 (8)
VH1-18 (8)
VH3-74 (8)
VH1-3 (8)
VH2-5 (10)
VH2-26 (10)
VH3-49 (8)
VH2-70 (10)
VH7-4-1 (8)
Δ nt
CDR H3 Length

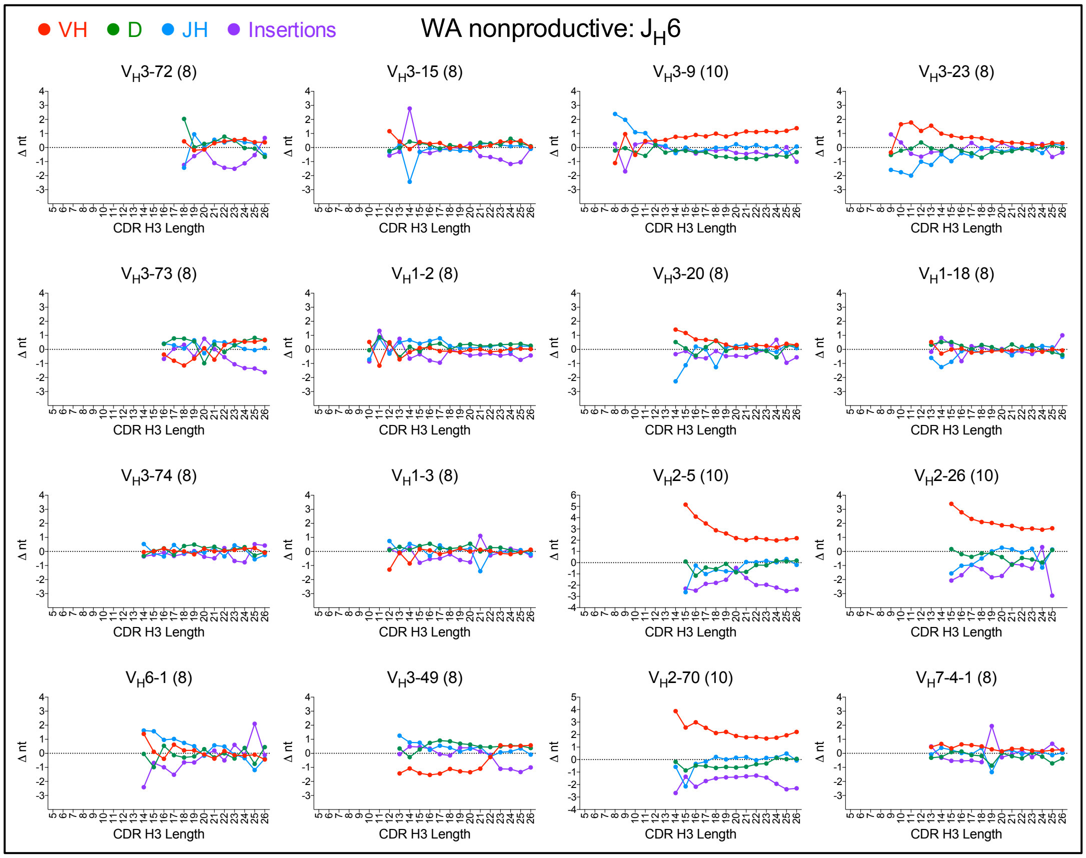
VH
D
JH
Insertions
WA nonproductive: JH6
VH3-72 (8)
VH3-15 (8)
VH3-9 (10)
VH3-23 (8)
VH3-73 (8)
VH1-2 (8)
VH3-20 (8)
VH1-18 (8)
VH3-74 (8)
VH1-3 (8)
VH2-5 (10)
VH2-26 (10)
VH6-1 (8)
VH3-49 (8)
VH2-70 (10)
VH7-4-1 (8)
Δ nt
CDR H3 Length

**Fig. S15.** Deviations from average length for $V_H$, D, $J_H$ and NP-regions (insertions) in clones segregated by $V_H$ and $J_H$ germlines. Deviations are calculated by subtracting average values for each region from overall repertoire averages within donors. Data points show averages of at least 3 donors from each data set for $V_H$/$J_H$ combinations with at least 60 counts. SRI panels highlighted by blue diamonds indicate $V_H$/$J_H$ combinations also analyzed in the WA naïve dataset. Error bars omitted for clarity. Note similarity of trends between WA and SRI datasets and lack of trends in WA unproductive sequences except for longer $V_H$ sequences for $V_H$2 and $V_H$3-9, as expected, and a trend for shorter than expected lengths of nucleotide insertions in $V_H$2 sequences.

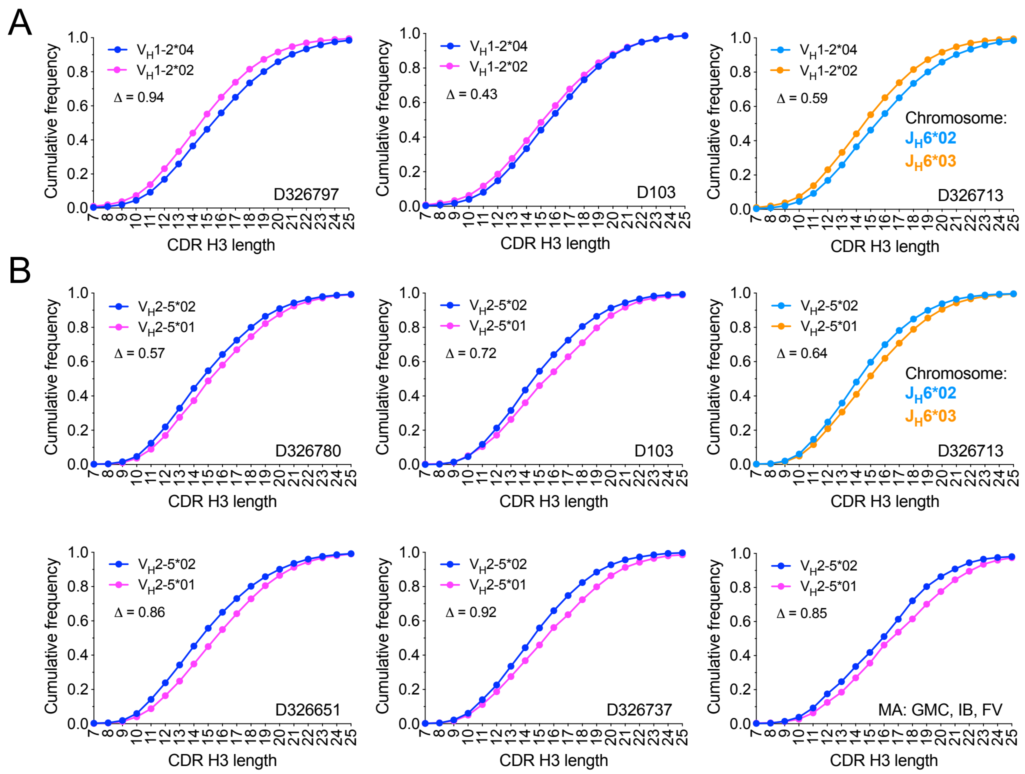


**Fig. S16.** CDR H3 length distributions associated with $V_H$1-2 (A) and $V_H$2-5 (B) allelic variants in the SRI and MA datasets. All panels except the lower right are SRI dataset donors. Only heterozygous donors for these germlines are shown, indicated in each panel. The 3 MA donors are heterozygous for $V_H$2-5 and were pooled in the lower right panel due to relatively low counts for each allele in individual donors. For SRI donors, only IgM/naïve sequences are included. For the MA donors, only IgM sequences with up to 1 amino acid mutation in IMGT® positions 1 to 104 were included. Light blue and orange symbols in donor D326713 distributions indicate haplotypes based on $J_H$6 alleles (1) associated with each $V_H$1-2 and $V_H$2-5 allele for donor D326713. Dark blue and magenta symbols indicate alleles for which haplotype analysis based on $J_H$6 alleles was not possible. Note the opposite biases of $V_H$1-2 and $V_H$2-5 alleles in the same chromosome for donor D326713, indicating that haplotype-associated variations of D germlines do not directly determine allele-associated CDR H3 length biases. The difference in average CDR H3 length between alleles is shown in each panel. All CDR H3 distribution differences are statistically significant in a Mann-Whitney test ($P < 10^{-4}$). $V_H$2-5 alleles vary by Asn/Asp-59 and $V_H$1-2 alleles by Arg/Trp-75. Additional variations in the framework region 1 may occur but are not covered in the SRI dataset. Other allelic variants were not analyzed due to lack of sufficient number of heterozygous donors with same alleles or insufficient allele-specific sequence counts.

| Germline | IMGT position 105 | 106 | 107 | Bias group |
|---|---|---|---|---|
| IGHV3-66 | A | R | (D) | Short |
| IGHV3-7 | A | R | (D) | |
| IGHV3-72 | A | R | (D) | |
| IGHV3-74 | A | R | (D) | |
| IGHV6-1 | A | R | (D) | |
| IGHV3-53 | A | R | (D) | |
| IGHV3-73 | T | R | (Q) | |
| IGHV3-15 | T | T | (D) | |
| IGHV2-5 | A | H | **R** | Cut |
| IGHV2-70 | A | R | I | |
| IGHV5-51 | A | R | (Q) | |
| IGHV3-20 | A | R | (D) | Crested |
| IGHV3-30 | A | R | (D) | |
| IGHV3-30-3 | A | R | (D) | |
| IGHV3-33 | A | R | (D) | |
| IGHV3-64 | A | R | (D) | |
| IGHV3-64D | A | R | (D) | |
| IGHV3-23 | A | K | (D) | |
| IGHV3-9 | A | K | **D** | |
| IGHV1-18 | A | R | (D) | Long |
| IGHV1-69 | A | R | (D) | |
| IGHV2-26 | A | R | I | |
| IGHV4-34 | A | R | (G) | |
| IGHV1-24 | A | T | (D) | |
| IGHV1-2 | A | R | (D) | Neutral |
| IGHV1-3 | A | R | (D) | |
| IGHV3-11 | A | R | (D) | |
| IGHV3-21 | A | R | (D) | |
| IGHV3-48 | A | R | (D) | |
| IGHV3-49 | T | R | (D) | |
| IGHV4-30-2 | A | R | (D) | |
| IGHV4-30-4 | A | R | (D) | |
| IGHV4-4 | A | R | (D) | |
| IGHV4-61 | A | R | (D) | |
| IGHV4-31 | A | R | (D) | |
| IGHV4-59 | A | R | (D) | |
| IGHV1-8 | A | R | (G) | |
| IGHV4-39 | A | R | (Q) | |
| IGHV7-4-1 | A | R | X | |

**Fig. S17.** CDR H3 residues encoded by $V_H$ germlines in the absence of nucleotide trimming. Residues in parentheses show partial codons and favored encoded residue in the absence of nucleotide trimming.

**Table S1. B cell subsets and dataset sample size before and after filtering by clonotype**

| | Dataset | | | | | | | |
|---|---|---|---|---|---|---|---|---|
| | CA | MA | TX | | | WA | | SRI |
| Donors | 3 | 3 | 3[a] | 2[a] | 2[a] | 3 | 3 | 8 |
| CD27 marker | NA[b] | NA | CD27$^{neg}$ | CD27$^{pos}$ | CD27$^{pos}$ | CD27$^{neg}$ | CD27$^{pos}$ | NA |
| Isotype | IgG | IgG, IgA[c] | IgM/Null[d] | IgG, IgA | IgM | NA | NA | IgG/IgM |
| Clonotype definition germlines | $V_H + V_L$ | $V_H + J_H$ | $V_H + V_L$ | $V_H + V_L$ | $V_H + V_L$ | $V_H + J_H$[e] | $V_H + J_H$[e] | $V_H + J_H$ |
| Clonotype CDR H3 amino acid identity[f] | 60% | 60% | 60% | 60% | 60% | 100% | 100% | 100% |
| Raw counts | 68,727 | 233,724 | 55,210 | 81,786 | 26,447 | $2.2 \times 10^7$ | $1.9 \times 10^7$ | $1.9 \times 10^8$ |
| Unique clonotypes[g] | 63,503 | 63,051 | 52,993 | 67,158 | 22,616 | $1.7 \times 10^7$ | $8.3 \times 10^6$ | IgG/IgA: $5.9 \times 10^6$<br>IgM/naïve[h]: $2.5 \times 10^7$ |
| Unproductive | NA | NA | NA | NA | NA | $3.1 \times 10^6$ | Not used | NA |

[a] Dataset with 3 donors, only 2 of which were processed for CD27$^{pos}$ B cells.

[b] Not available.

[c] IgM sequences were not included in analyses for the MA dataset.

[d] "Null" isotype sequences in the CD27$^{neg}$ compartment were assumed to be IgM.

[e] Sequences with undefined $V_H$ or $J_H$ germlines grouped by clonotype as described in methods.

[f] Average 60% amino acid identity across CDR H3 lengths achieved by using a nominal 57% identity threshold.

[g] Number of sequences after filtering for clonotypes as described in methods.

[h] IgM/naïve sequences in the SRI dataset defined as IgM sequences without amino acid mutations between IMGT® Cys-23 and Cys-104.

**Table S2. Ambiguity in $V_H$ germline calls in the AE and naïve B cell compartments of the WA dataset[a]**

| Germline | AE compartment: | | | Naïve compartment: | | | Included in analyses |
|---|---|---|---|---|---|---|---|
| | Unambiguous | In ties | Ambiguous | Unambiguous | In ties | Ambiguous | |
| IGHV1-2 | 186646 | 26646 | 12% | 492797 | 7003 | 1% | Yes |
| IGHV1-3 | 159549 | 15495 | 9% | 321611 | 2513 | 1% | Yes |
| IGHV1-18 | 305053 | 34857 | 10% | 768812 | 7397 | 1% | Yes |
| IGHV2-5 | 125943 | 11499 | 8% | 312664 | 9487 | 3% | Yes |
| IGHV2-26 | 40792 | 4713 | 10% | 137338 | 3120 | 2% | Yes |
| IGHV2-70 | 82501 | 14979 | 15% | 208336 | 12288 | 6% | Yes |
| IGHV3-9 | 87467 | 8318 | 9% | 225624 | 14103 | 6% | Yes |
| IGHV3-15 | 180906 | 6485 | 3% | 611241 | 1474 | 0% | Yes |
| IGHV3-20 | 48984 | 6588 | 12% | 146903 | 1395 | 1% | Yes |
| IGHV3-23 | 970420 | 69298 | 7% | 1815159 | 4669 | 0% | Yes |
| IGHV3-49 | 100337 | 3985 | 4% | 279063 | 1171 | 0% | Yes |
| IGHV3-72 | 55488 | 2988 | 5% | 35571 | 224 | 1% | Yes |
| IGHV3-73 | 50904 | 1195 | 2% | 108340 | 353 | 0% | Yes |
| IGHV3-74 | 192071 | 10882 | 5% | 230877 | 957 | 0% | Yes |
| IGHV6-1 | 44906 | 862 | 2% | 52641 | 319 | 1% | Yes |
| IGHV7-4-1 | 50503 | 1376 | 3% | 106564 | 731 | 1% | Yes |
| IGHV3-43 | 49473 | 5122 | 9% | 119476 | 13034 | 10% | No |
| IGHV1-8 | 43443 | 66316 | 60% | 122649 | 205349 | 63% | No |
| IGHV1-14 | 750 | 1371 | 65% | 736 | 894 | 55% | No |
| IGHV1-17 | 10177 | 306 | 3% | 27385 | 17 | 0% | No |
| IGHV1-24 | 20909 | 21051 | 50% | 55192 | 42559 | 44% | No |
| IGHV1-45 | 2467 | 655 | 21% | 5136 | 31 | 1% | No |
| IGHV1-46 | 215404 | 70218 | 25% | 403610 | 184105 | 31% | No |
| IGHV1-58 | 14300 | 1445 | 9% | 23185 | 277 | 1% | No |
| IGHV1-67 | 5968 | 6063 | 50% | 1653 | 140 | 8% | No |
| IGHV1-68 | 305 | 96 | 24% | 298 | 0 | 0% | No |
| IGHV1-69 | 235691 | 107375 | 31% | 938608 | 286200 | 23% | No |
| IGHV1-c | 2341 | 53 | 2% | 4840 | 2 | 0% | No |

| | | | | | | | |
|---|---|---|---|---|---|---|---|
| IGHV1-f | 21621 | 21123 | 49% | 45142 | 42554 | 49% | No |
| IGHV2-10 | 359 | 1 | 0% | 435 | 0 | 0% | No |
| IGHV3-6 | 45 | 25 | 36% | 3 | 0 | 0% | No |
| IGHV3-7 | 373 | 824141 | 100% | 55 | 2410458 | 100% | No |
| IGHV3-11 | 52183 | 539883 | 91% | 82250 | 1534126 | 95% | No |
| IGHV3-13 | 16956 | 1480 | 8% | 20999 | 247 | 1% | No |
| IGHV3-16 | 252 | 150 | 37% | 2 | 72 | 97% | No |
| IGHV3-19 | 3775 | 186 | 5% | 4308 | 73 | 2% | No |
| IGHV3-21 | 6904 | 821948 | 99% | 5107 | 2405759 | 100% | No |
| IGHV3-22 | 4942 | 600 | 11% | 9275 | 47 | 1% | No |
| IGHV3-25 | 1111 | 656 | 37% | 221 | 12 | 5% | No |
| IGHV3-30 | 47023 | 708534 | 94% | 2756 | 1891771 | 100% | No |
| IGHV3-32 | 27 | 11 | 29% | 1 | 0 | 0% | No |
| IGHV3-33 | 50506 | 211019 | 81% | 2409 | 303610 | 99% | No |
| IGHV3-35 | 2242 | 515 | 19% | 405 | 49 | 11% | No |
| IGHV3-37 | 56 | 24 | 30% | 9 | 1 | 10% | No |
| IGHV3-38 | 939 | 627 | 40% | 33 | 25 | 43% | No |
| IGHV3-41 | 2940 | 83 | 3% | 4487 | 5 | 0% | No |
| IGHV3-42 | 221 | 112 | 34% | 17 | 11 | 39% | No |
| IGHV3-47 | 4546 | 399 | 8% | 3400 | 9 | 0% | No |
| IGHV3-48 | 148865 | 692459 | 82% | 205941 | 1987958 | 91% | No |
| IGHV3-52 | 3981 | 131 | 3% | 6272 | 7 | 0% | No |
| IGHV3-53 | 171951 | 56317 | 25% | 244472 | 25071 | 9% | No |
| IGHV3-54 | 259 | 45 | 15% | 2 | 7 | 78% | No |
| IGHV3-57 | 76 | 182 | 71% | 1 | 4 | 80% | No |
| IGHV3-60 | 1100 | 1077 | 49% | 14 | 30 | 68% | No |
| IGHV3-62 | 64 | 203 | 76% | 1 | 2 | 67% | No |
| IGHV3-64 | 60179 | 273510 | 82% | 82328 | 777265 | 90% | No |
| IGHV3-65 | 2369 | 783 | 25% | 2427 | 72 | 3% | No |
| IGHV3-66 | 106999 | 77989 | 42% | 149442 | 87494 | 37% | No |
| IGHV3-71 | 26736 | 82189 | 75% | 11087 | 196702 | 95% | No |
| IGHV3-75 | 112 | 8 | 7% | 5 | 2 | 29% | No |

| | | | | | | | |
|---|---|---|---|---|---|---|---|
| IGHV3-76 | 127 | 27 | 18% | 103 | 0 | 0% | No |
| IGHV3-79 | 5 | 35 | 88% | 1 | 0 | 0% | No |
| IGHV3-d | 1511 | 1513 | 50% | 1338 | 184 | 12% | No |
| IGHV4-4 | 877 | 271677 | 100% | 74 | 447152 | 100% | No |
| IGHV4-28 | 3205 | 3043 | 49% | 2689 | 1058 | 28% | No |
| IGHV4-30-2 | 15651 | 350263 | 96% | 28860 | 932945 | 97% | No |
| IGHV4-30-4 | 3664 | 60737 | 94% | 267 | 198674 | 100% | No |
| IGHV4-31 | 3542 | 204323 | 98% | 159 | 367420 | 100% | No |
| IGHV4-34 | 207848 | 205207 | 50% | 382365 | 333301 | 47% | No |
| IGHV4-39 | 109736 | 377346 | 77% | 171350 | 698882 | 80% | No |
| IGHV4-55 | 26072 | 99668 | 79% | 24046 | 129608 | 84% | No |
| IGHV4-59 | 34296 | 693791 | 95% | 11570 | 1719475 | 99% | No |
| IGHV4-61 | 21706 | 642300 | 97% | 3538 | 1596968 | 100% | No |
| IGHV4-80 | 48 | 9 | 16% | 2 | 0 | 0% | No |
| IGHV4-b | 28075 | 110644 | 80% | 53745 | 229522 | 81% | No |
| IGHV5-51 | 360757 | 44331 | 11% | 829787 | 124676 | 13% | No |
| IGHV5-78 | 3751 | 712 | 16% | 44 | 18 | 29% | No |
| IGHV5-a | 159275 | 43948 | 22% | 401534 | 124666 | 24% | No |
| IGHV7-27 | 238 | 9 | 4% | 337 | 0 | 0% | No |
| IGHV7-34-1 | 25 | 98 | 80% | 6 | 12 | 67% | No |
| IGHV7-40 | 1177 | 677 | 37% | 1785 | 458 | 20% | No |
| IGHV7-56 | 198 | 35 | 15% | 58 | 1 | 2% | No |
| IGHV7-81 | 249 | 389 | 61% | 31 | 121 | 80% | No |

[a] Data from donors 1, 2 and 3 pooled and analyzed for ambiguous ("in ties") germline calls after clonotype clustering. Minor germlines excluded for brevity.

**Table S3. MA dataset sample source from the SRA archive**

| Run | BioSample | Sample name | Experiment | Donor | Timepoint[a] |
|---|---|---|---|---|---|
| SRR4431793 | SAMN05924285 | FV_before_8d | SRX2251687 | Fv | -8d |
| SRR4431792 | SAMN05924284 | FV_before_2d | SRX2251686 | Fv | -2d |
| SRR4431787 | SAMN05924283 | FV_before_1h | SRX2251681 | Fv | -1h |
| SRR4431789 | SAMN05924277 | FV_after_1h | SRX2251683 | Fv | +1h |
| SRR4431788 | SAMN05924276 | FV_after_1d | SRX2251682 | Fv | +1d |
| SRR4431784 | SAMN05924280 | FV_after_3d | SRX2251678 | Fv | +3d |
| SRR4431790 | SAMN05924278 | FV_after_1w | SRX2251684 | Fv | +1w |
| SRR4431791 | SAMN05924279 | FV_after_2w | SRX2251685 | Fv | +2w |
| SRR4431785 | SAMN05924281 | FV_after_3w | SRX2251679 | Fv | +3w |
| SRR4431786 | SAMN05924282 | FV_after_4w | SRX2251680 | Fv | +4w |
| SRR4431783 | SAMN05924295 | GMC_before_8d | SRX2251677 | GMC | -8d |
| SRR4431782 | SAMN05924294 | GMC_before_2d | SRX2251676 | GMC | -2d |
| SRR4431775 | SAMN05924293 | GMC_before_1h | SRX2251669 | GMC | -1h |
| SRR4431781 | SAMN05924287 | GMC_after_1h | SRX2251675 | GMC | +1h |
| SRR4431780 | SAMN05924286 | GMC_after_1d | SRX2251674 | GMC | +1d |
| SRR4431776 | SAMN05924290 | GMC_after_3d | SRX2251670 | GMC | +3d |
| SRR4431778 | SAMN05924288 | GMC_after_1w | SRX2251672 | GMC | +1w |
| SRR4431779 | SAMN05924289 | GMC_after_2w | SRX2251673 | GMC | +2w |
| SRR4431777 | SAMN05924291 | GMC_after_3w | SRX2251671 | GMC | +3w |
| SRR4431774 | SAMN05924292 | GMC_after_4w | SRX2251668 | GMC | +4w |
| SRR4431772 | SAMN05924305 | IB_before_8d | SRX2251666 | IB | -8d |
| SRR4431773 | SAMN05924304 | IB_before_2d | SRX2251667 | IB | -2d |
| SRR4431768 | SAMN05924303 | IB_before_1h | SRX2251662 | IB | -1h |
| SRR4431766 | SAMN05924297 | IB_after_1h | SRX2251660 | IB | +1h |
| SRR4431767 | SAMN05924296 | IB_after_1d | SRX2251661 | IB | +1d |
| SRR4431771 | SAMN05924300 | IB_after_3d | SRX2251665 | IB | +3d |
| SRR4431765 | SAMN05924298 | IB_after_1w | SRX2251659 | IB | +1w |
| SRR4431764 | SAMN05924299 | IB_after_2w | SRX2251658 | IB | +2w |
| SRR4431770 | SAMN05924301 | IB_after_3w | SRX2251664 | IB | +3w |
| SRR4431769 | SAMN05924302 | IB_after_4w | SRX2251663 | IB | +4w |

[a] Sample time relative to vaccination as described by Laserson *et al*. (2).
